# Complex bacterial diversity of Guaymas Basin hydrothermal sediments revealed by synthetic long-read sequencing (LoopSeq)

**DOI:** 10.1101/2024.04.12.589229

**Authors:** John E. Hinkle, Jeffrey P. Chanton, Molly A. Moynihan, S. Emil Ruff, Andreas Teske

## Abstract

Hydrothermal sediments host phylogenetically diverse and physiologically complex microbial communities. Previous studies of microbial community structure in hydrothermal sediments have typically used short-read sequencing approaches. To improve on these approaches, we use LoopSeq, a high-throughput synthetic long-read sequencing method that has yielded promising results in analyses of microbial ecosystems, such as the human gut microbiome. In this study, LoopSeq is used to obtain near-full length (approximately 1400 - 1500 nucleotides) bacterial 16S rRNA gene sequences from hydrothermal sediments in Guaymas Basin. Based on these sequences, high-quality alignments and phylogenetic analyses provided new insights into previously unrecognized taxonomic diversity of sulfur-cycling microorganisms and their distribution along a lateral hydrothermal gradient. Detailed phylogenies for free-living and syntrophic sulfur-cycling bacterial lineages identified well-supported monophyletic clusters that have implications for the taxonomic classification of these groups. Particularly, we identify clusters within *Candidatus* Desulfofervidus that represent unexplored physiological and genomic diversity. In general, LoopSeq-derived 16S rRNA gene sequences aligned consistently with reference sequences in GenBank; however, chimeras were prevalent in sequences as affiliated with the thermophilic *Candidatus* Desulfofervidus and *Thermodesulfobacterium*, and in smaller numbers within the sulfur-oxidizing family *Beggiatoaceae*. Our analysis of sediments along a well-documented thermal and geochemical gradient show how sulfur-cycling bacteria of different taxonomic groups persist as active catalysts of the sulfur cycle throughout surficial hydrothermal sediments in the Guaymas Basin.

## INTRODUCTION

The Guaymas Basin is a marginal rift basin in the Gulf of California that is characterized by the coexistence of hydrothermal vents and cold seeps. Unlike many other regions with hydrothermal vents, Guaymas Basin is subject to relatively high sediment accumulation rates (Calvert, 1966). Basaltic sills, freshly emplaced into organic-rich sediments, drive the hydrothermal circulation of hydrocarbons, sulfide, and nutrient-rich fluids (Einsele et al. 1980, Lizarralde et al., 2011). Abundant energy and carbon sources sustain highly active and very diverse microbial communities at the sediment surface (Teske et al., 2021). Consequently, Guaymas Basin has been the subject of many sequencing studies throughout the past two decades, starting with the use of near-full-length (∼1500 nucleotides) cloned PCR products (Teske et al., 2002), then progressing to high throughput pyrosequencing of very short (∼80 nucleotides) 16S rRNA gene fragments (Ruff et al., 2015). More recent studies utilize high throughput sequencing methods that produce longer (∼400-500 nucleotides) 16S rRNA gene fragments (Teske et al., 2019, Su et al., 2023). The abundance of sequence data obtained via different methods makes the Guaymas Basin an ideal model system to assess the performance of new next-generation sequencing technologies, such as LoopSeq.

LoopSeq is a high-throughput synthetic long-read sequencing method, which can generate near-full length gene sequences (Callahan et al., 2021). Studies using clone library construction and sequencing methods yield near full-length and high-quality sequences but are greatly limited by throughput and sequencing depth. Clone library-based methods are time consuming and only the most abundant microorganisms are usually detected. In turn, high throughput sequencing methods can yield tens of thousands of sequence reads, allowing to detect extremely rare organisms (Sogin et al., 2006). Yet, these reads are too short for high-resolution phylogenetic analyses and can only be used for relatively basic taxonomic identification. Short reads of 400-500 nucleotides can be used to construct phylogenies, but placement of unknown or poorly resolved lineages remains problematic, due to the limited information provided by short sequences and limited bootstrap support for phylogenetic branching patterns (Teske et al., 2019, Ramírez et al., 2021).

LoopSeq bridges this gap and allows investigators to obtain nearly full-length 16S rRNA gene sequences in high throughput, for costs comparable to short read amplicons. The combination of high throughput and nearly full-length sequences allows for thorough surveys of the diversity of known microbial lineages, as well as high-resolution phylogenetic placement of rare or poorly resolved microbial lineages even for highly diverse ecosystems, such deep-sea hydrothermal sediments. To test the capabilities and reliability of LoopSeq for environmental samples, Guaymas Basin is ideal because i) the ecosystem is generally well understood, ii) the hydrothermal sediments are very diverse, iii) the sediments are rich in organics that can be problematic for sequencing, e.g. hydrocarbons, and iv) extensive 16S rRNA gene sequence databases are available to verify LoopSeq sequences against publicly available short read and full-length sequences.

## METHODS

### Sampling

Guaymas Basin sites were visited and sampled with R/V *Atlantis*, HOV *Alvin*, and AUV *Sentry* during cruise AT37-06 (December 6-29, 2016). *Alvin* dives targeted previously explored sampling areas in the southern axial valley of Guaymas Basin (Teske et al., 2016), or newly identified sites found by AUV *Sentry*. Hydrothermal sediment samples (*Alvin* push cores) used in this study were obtained during *Alvin* Dive 4872 (Dec. 24, 2016) in the Cathedral Hill area (27.00.684 N/ 111.24.266 W) at 2014 m depth. The sample set covers a hydrothermal gradient from hot hydrothermally charged sediments covered with orange and white *Beggiatoaceae* mats to temperate bare sediment, within ∼1 m distance (Fig. 1). Core 4872-01 represents bare sediment that was not covered by a *Beggiatoaceae* mat, whereas Core 4872-06 was covered by a white *Beggiatoaceae* mat, and Core 4872-14 was covered by an orange *Beggiatoaceae* mat. Once the sediment cores were returned to the surface, porewater profiles were collected using rhizons (Rhizosphere Research Products, Wageningen, The Netherlands). Then, these cores were divided into 3 cm sections and methane samples were collected by injecting 2 mL sediment plugs into 30 mL serum vials containing 2 ml of 1 M sodium hydroxide solution each. The serum vials were immediately sealed with thick rubber stoppers, crimped with aluminum seals, and stored at 4 °C. The remaining sediment samples were frozen at −80°C by the shipboard science crew, and later used for DNA extraction shoreside.

**Figure 1.**
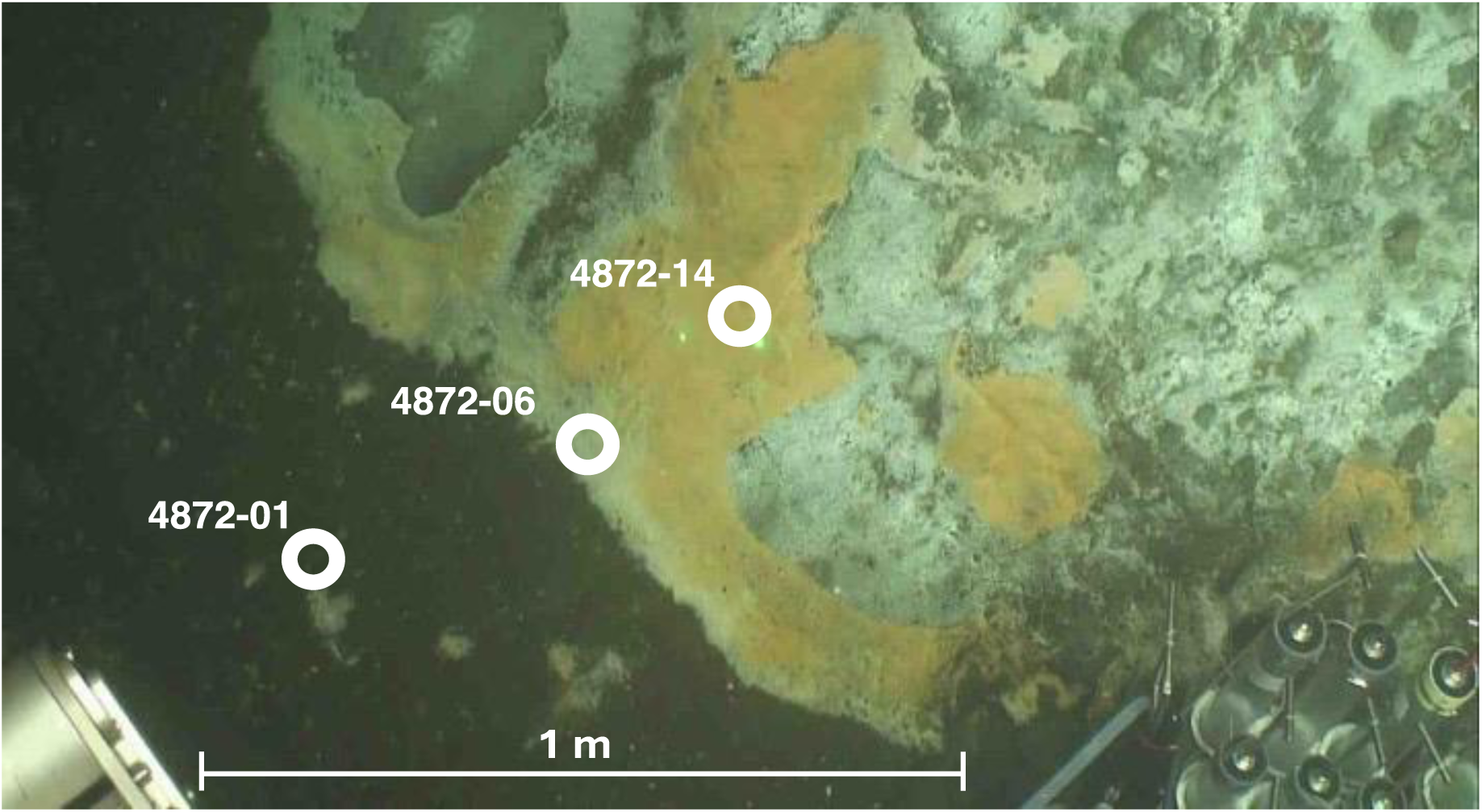
Seafloor image of sampling mat area at Dive 4872. Approximate location of push coring sites are labeled in white. Image courtesy of HOV *Alvin* Dive Team.

### Thermal profiles

Thermal profiles were measured in surficial sediments using *Alvin’s* 50 cm heat flow probe (https://ndsf.whoi.edu/alvin/using-alvin/sampling-equipment/). The 50 cm probe contains thermal sensors every 10 cm, starting 5 cm below the attached plastic disc (the “puck”) that limits probe penetration and rests on the seafloor once the probe was inserted. After 5 to 10 minutes of temperature reading stabilization, temperature readings were recorded.

### Geochemical analyses

For sulfide analysis, 1 mL porewater subsamples were fixed with 0.1 mL of 0.1 M zinc acetate solution to preserve sulfide as zinc sulfide until analysis using the methylene blue method (Cline, 1969). For sulfate measurements, porewater samples were diluted in water (1:50) and analyzed using ion chromatography (Metrohm 930 Compact IC flex with Metrosep A PCC HC/4.0 preconcentration column, and Metrosep A Supp 5 Guard/4.0 chromatography column). Sulfate for the white and orange cores was measured by Wetland Biogeochemistry Analytical Services (WBAS) using ion chromatography on a Dionex ICS-1000 using a RFIC IonPac AS22 4 × 250 mm column and a RFIC IonPac AG22 Guard column 4 × 50 mm. The IC was calibrated using a 5-point standard curve and a minimum R^2^ value of 0.999. The detection limit measured at 0.10 ppm sulfate. A check standard was made from a Dionex 7 anion standard (1000 ppm) to a concentration of 15 ppm. An external Standard from HACH (100 ppm sulfate) was used to make a 20 ppm quality check standard. In general, quality checks were analyzed for 10 % of the samples.

Porewaters from the Guaymas Basin were analyzed at WBAS for nitrate/nitrite (NO_x_) and ammonium (NH_4_^+^) concentrations colorimetrically, using a Flow Solutions IV segmented flow Auto Analyzer from O.I Analytical, College Station, TX. The HCl-acidified samples were neutralized with sodium hydroxide (NaOH) before analysis. NO_x_ was determined using the cadmium reduction method and NH_4_^+^ was determined using the phenate method. Both nutrients were diluted to get their concentrations within the linear range of the auto analyzer. Quality control standards from certified stock standards (Environmental Research Associates), were analyzed every 15-20 samples.

For combined concentration and δ^13^C analysis of methane, 2 mL sediment subsamples were added to 30 mL serum vials containing 2 mL of 1 M NaOH solution, sealed with thick butyl rubber stoppers, crimped with aluminum seals, and stored at 4°C. Since cores were retrieved unpressurized, outgassing may have particularly impacted the measurements of methane concentrations near and above saturation. After the cruise, the methane samples were analyzed by headspace gas chromatography-flame ionization detection (GC-FID) at Florida State University (Magen et al., 2014). Additionally, the gas samples were analyzed for δ^13^CH_4_ by injecting 0.1 to 0.5 mL of sample into a gas chromatograph interfaced to a Finnigan MAT Delta S Isotope Ratio Mass Spectrometer inlet system as previously described (Chanton and Liptay 2000). Values are reported in the per mil (‰) notation relative to Vienna Pee Dee Belemnite (VPDB).

### DNA extraction

DNA was extracted from Guaymas Basin sediment samples using a modified version of the protocol developed by Zhou et al., 1996 (see supplementary material). The extraction method was optimized for deep-sea sediments. The detailed protocol can be found in the supplementary materials.

### LoopSeq sequencing

Sequencing libraries were prepared from extracted genomic DNA with the commercially available LoopSeq kits from Loop Genomics (https://go.elementbiosciences.com/16s-loopseq-user-guide, also available in the supplementary material). Synthetic long reads (SLRs) were constructed from the short-read sequencing reads using the standard Loop Genomics informatics pipeline and processed as previously described (Callahan et al., 2021). The method involves attaching two DNA tags: one Unique Molecular Identifier (UMI) to each unique “parent” DNA molecule and one sample-specific tag (i.e., a sample index) equally to all molecules in the same sample. Barcoded molecules are amplified, multiplexed, and each UMI is distributed intramolecularly to a random position within each parent molecule. Molecules are then fragmented into smaller units at the position of each UMI, creating a library of UMI-tagged fragments with an average length of 400 bp compatible with an Illumina sequencing platform run in PE150 mode (Callahan et al., 2021).

This protocol allows for the assembly of continuous 16S rRNA gene long reads from individual DNA molecules, providing sequencing data like classical long read sequencing methods but with much higher throughput. The terminal primers determine the extent of the 16S rRNA gene sequence yielded by this approach (27F, 5’-AGAGTTTGATCMTGGCTCAG-3’; 1492R, 5’-TACCTTGTTACGACTT-3’). LoopSeq can be performed on existing DNA sequencing instruments, and due to the chemistry and the tagging of individual molecules the method is semi-quantitative providing a major advantage towards other amplification-based methods.

We note that LoopSeq reconstructs a single long read from several short reads. Therefore, we define the resulting amplicon sequence variants (ASVs) as LoopSeq variants (L-ASVs) to differentiate them from “traditional” ASVs that are essentially PCR amplicons.

### DADA2 analysis

The statistical programming language R 4.2.1 (R Core Team, 2022) and the DADA2 (v. 1.20.0) bioinformatics package (Callahan et al., 2016), along with a LoopSeq suitable DAD2 workflow (Callahan et al., 2021) were used to analyze 16S rRNA gene sequences obtained by LoopSeq sequencing. DADA2 creates ASVs (or LoopSeq variants (L-ASVs) in our case) by grouping together amplicon reads of the same sequence.

The function “removePrimers” was used to select for 16S rRNA amplicon sequences with both the forward and reverse primers and allowing a maximum mismatch of two base pairs, as reads lacking either primer do not represent the complete gene. The following parameters were used to filter the remaining sequences: a minimum length of 1000 bp (minLen=1000), a maximum length of 2000 bp (maxLen=2000), a minimum quality score of 3 (minQ=3), a maximum expected error rate of 1 (maxEE=1), and a maximum Ns of 0 (maxN = 0). These parameters may be used to filter reads without selecting for reads with a forward and reverse primer to capture long, but not complete genes. We found that 20.4% of reads that fulfilled the previously stated parameters lacked the forward/reverse primer. Only full-length gene sequences were kept in our analysis. The function “dereqFastq” was used to dereplicate reads. The function “learnErrors” was used with a band size of 32 (BAND_SIZE=32) to learn error rates. The error estimate function was set to the PacBio Error Function. DADA2 was ran with a p-value cutoff of 1^-10^ (OMEGA_A=1e-10), single detection (DETECT_SINGLETONS=TRUE), and a band size of 32 (BAND_SIZE=32). The function “removeBimeraDenovo” (method = consensus) was used to remove chimeras. An overview of raw read processing is depicted in Supplementary Table 2.

The function “assignTaxonomy” was used to assign taxonomy using the Silva reference database (v138) (Quast et al., 2013). A total of 12,494 L-ASVs were identified. Figures depicting taxonomic distribution, alpha diversity, and beta diversity (Fig. 3, Suppl. Figs 2 & 3) were created using the R Phyloseq package (McMurdie & Holmes, 2013). All sequence reads were submitted to the NCBI Sequence Read Archive (BioProject: PRJNA1105367).

### 16S rRNA gene sequence phylogeny

We compared our L-ASV sequences against reference sequences obtained from the NCBI GenBank database. All sequences were aligned using Multiple Sequence Comparison by Log-Expectation (MUSCLE) in MEGA11 (Stecher et al., 2020). From these alignments, MEGA11 was used to construct minimum-evolution phylogenetic trees checked by the aggregate of 1000 bootstrap replicates (Rzhetsky & Nei, 1992, Felsenstein, 1985). Bootstrap values greater than 70% are shown and NCBI GenBank accession numbers are provided on tip labels.

### 16S rRNA gene sequence distance matrices

Distance matrices corresponding to each phylogenetic tree are shown in the Supplementary Data File. The number of base substitutions between sequences are shown as a percentage (i.e., per 100 sites). Analyses were conducted using the Maximum Composite Likelihood model (Tamura et al., 2004). Each analysis involved all 16S rRNA gene sequences displayed in the corresponding phylogenetic tree. All ambiguous positions were removed for each sequence pair (pairwise deletion option). Evolutionary analyses were conducted in MEGA11 (Tamura et al., 2021, Stecher et al., 2020).

## RESULTS AND DISCUSSION

### Sediment characteristics

The color of Guaymas Basin *Beggiatoaceae* mats indicated the relative temperature of the underlying sediment as measured using the *Alvin* heat flow probe; orange mats covered hotter sediments and white mats covered comparatively cooler sediments (McKay et al., 2012). Temperature varied greatly with increasing sediment depth at the three sample sites (Figs. 1 & 2a). Temperatures at 40 centimeters below the seafloor (cmbsf) exceeded 100 °C below the orange mat, reached 75 °C below the white mat, and remained near 20 °C in sediments without a *Beggiatoaceae* mat (Fig. 2a).

**Figure 2.**
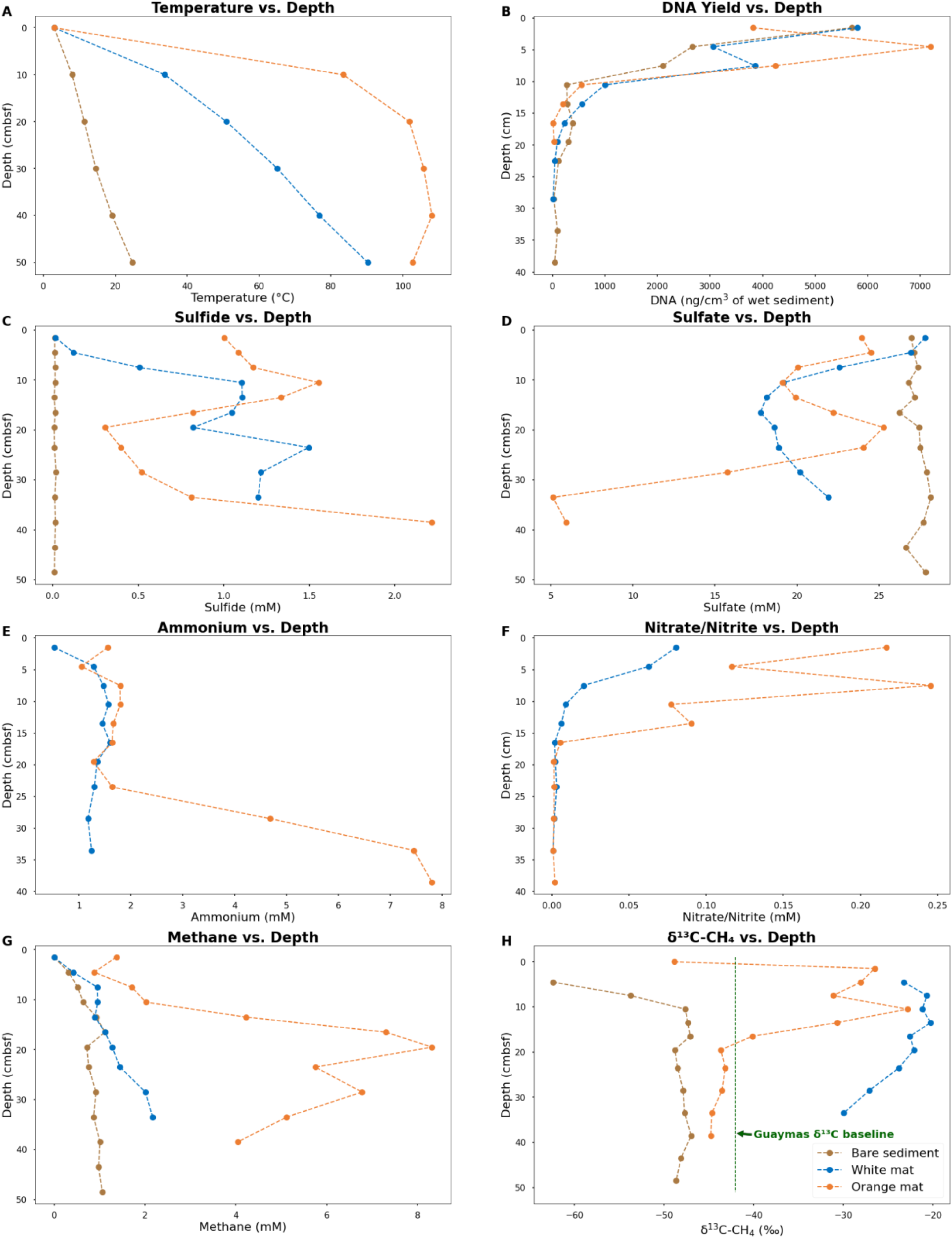
Plots of in-situ temperatures and geochemical parameters for sediment cores. Sulfide, sulfate, Ammonium, and Nitrite/Nitrate concentrations obtained from porewater. Methane concentrations and δ^13^C values obtained from whole sediment. Ammonium and Nitrite/Nitrate concentrations are not available for the bare sediment site (Core 4872-01). Data for the bare sediment site (Core 4872-01) are plotted in brown. Data for the white mat site (Core 4872-06) are plotted in blue. Data for the orange mat site (Core 4872-14) are plotted in orange. Plotted data are available in Supplementary Table 1.

Temperature considerably impacts microbial population density, and therefore the amount of DNA available for extraction. Measurable concentrations of DNA persisted the deepest (40 cmbsf) in Core 1, from bare sediment neighboring the warm mats. In contrast, DNA recovery is limited to surficial sediments (∼ 20 cmbsf) in Core 14 from the hot center of the orange mat (Fig. 2b.). Interestingly, the highest DNA yields are found within the upper ∼6 cm of sediment in all three cores, indicating that the impact of hydrothermal activity extends beyond the *Beggiatoaceae* mats to the bare sediments of Core 1 (Fig. 2b). Previous research in Guaymas Basin consistently reveals a very rapid decline in cell counts and 16S rRNA gene abundances with increasing depth and temperature in hot hydrothermal sediments (Meyer et al., 2013, Møller et al., 2018, Lagostina et al., 2021).

Micromolar concentrations of sulfide are observed throughout the bare sediment core, whereas the white and orange mat cores reach concentrations of sulfide up to 2 millimolar (Fig. 2c). At the surface, sulfide concentrations are near zero for the bare sediment and white mat cores. In contrast, the orange mat core has elevated (∼1 millimolar) sulfide levels at the surface, consistent with high hydrothermal flow in the mat center. Sulfate is abundant in all three cores, with depletion only occurring at ∼32 cm and below in the orange mat core, indicating that hydrothermal circulation contributes to the replenishment of sulfate in surficial sediments (Fig. 2d).

Ammonium is observed throughout our white and orange mat sediment cores (Fig. 2e) Consistent with previous research, ammonium concentrations increase as temperature increases downcore (Ramírez et al., 2021). Ammonium in Guaymas Basin hydrothermal sediments is primarily derived from the thermal degradation of organic matter (Von Damm et al., 1985). Deep in the orange mat core, ammonium concentrations reach ∼8 mM (Fig. 2e), close to the ∼10-15 mM concentrations observed in Guaymas Basin hydrothermal fluids (Von Damm et al., 1985).

Elevated nitrate/nitrite concentrations are observed in surficial mat covered sediments (Fig. 2f). Bottom water concentrations of nitrate in Guaymas Basin are up to 50 μM (Winkel et al., 2014). Elevated nitrate levels (up to nearly 0.25 mM; Fig. 2f) in mat-covered surficial sediments are at least in part due to damaged *Beggiatoaceae* mats leaking nitrate stored in their vacuoles into surficial sediments (Schutte et al., 2018).

High concentrations of methane are observed in all three cores (Fig. 2g). A gradient of peak methane concentrations (∼1 mM in the bare sediment core, ∼2 mM in the white mat core, and ∼8 mM in the orange mat core) illustrates the impact of hydrothermal fluids that are supersaturated in methane. High methane concentrations (multiple millimolar) are a characteristic feature of Guaymas Basin hydrothermal sediments, similar to those found at cold seep sites (Welhan, 1988, McKay et al., 2016, Song et al., 2021).

Hydrothermal methane in Guaymas Basin unmodified by microbial activity has a δ^13^CH_4_ value of approximately −42 ‰ (McKay et al. 2016, Song et al. 2021). The preferential oxidation of ^12^CH_4_ by methane-oxidizing archaea leads to the orange and white mat cores containing methane with a heavier carbon isotopic signature that is less depleted in carbon-13, with values ranging from −30 to −20 ‰ (Fig. 2h). The lighter carbon isotopic signature of methane throughout the bare sediment core and near the surface of the orange and white mat cores, indicates some contribution due to microbial methanogenesis (Fig. 2h). The methanogens in these sediments are most likely methylotrophs, which unlike other types of methanogens are capable of successfully performing methanogenesis in sulfate-replete sediments (Zhuang et al., 2018).

### Inferred Carbon Sources

Guaymas Basin sediments are rich in organic carbon (De la Lanza-Espino & Soto, 1999) and harbor numerous heterotrophic microbial lineages (Pérez Castro et al., 2021). Carbon sources are likely to shift throughout hydrothermal gradients in the sediments. Aromatic hydrocarbons serve as widely available carbon sources for aromatic-degrading bacteria throughout surficial Guaymas Basin sediments (Mara et al., 2022, Gutierrez et al., 2015, Goetz & Jannasch, 1993). In contrast, alkanes and methane occur are more abundant near hydrothermally active “hot spots” (Song et al., 2021, McKay et al., 2016). The abundance of alkanes and methane in hot sediments contributes to the evolution of thermotolerance and thermophily in several groups of alkane and methane degrading bacteria and archaea (Zehnle et al., 2023, Benito Merino et al., 2022). The upper limit of hydrocarbon degradation near approximately 80 °C is supported by the increased abundance of ^13^C-depleted substrates, which reflects the absence of microbial degradation (Song et al., 2021).

### Sequencing Profile

LoopSeq recovered nearly full-length sequences (Suppl. Fig 1) that affiliated with several phyla of heterotrophic bacteria, such as most Proteobacteria Desulfobacterota, Bacteroidota, and Chloroflexi (Fig. 3). These phyla have been detected by previous Guaymas Basin sequencing surveys using clone libraries of near full-length 16S rRNA genes (Teske et al., 2002, Dowell et al., 2016, McKay et al., 2016). We analyzed the alpha diversity of each mat site using five measures (Observed, Chao1, Shannon, Simpson, and Inverse Simpson). Although we expected hydrothermal stress to influence alpha diversity, we found no significant difference in alpha diversity across these three cores (Suppl. Fig. 2). To explore potential differences in beta diversity, we created a non-metric multi-dimensional scaling (NMDS) plot using the Bray-Curtis dissimilarity method. The bare sediment samples largely cluster together, particularly the deeper samples collected below 21 cmbsf (Suppl. Fig. 3. We also observe a cluster of surficial sediment samples (the top 6 cmbsf) from all three coring sites (Suppl. Fig. 3).

**Figure 3.**
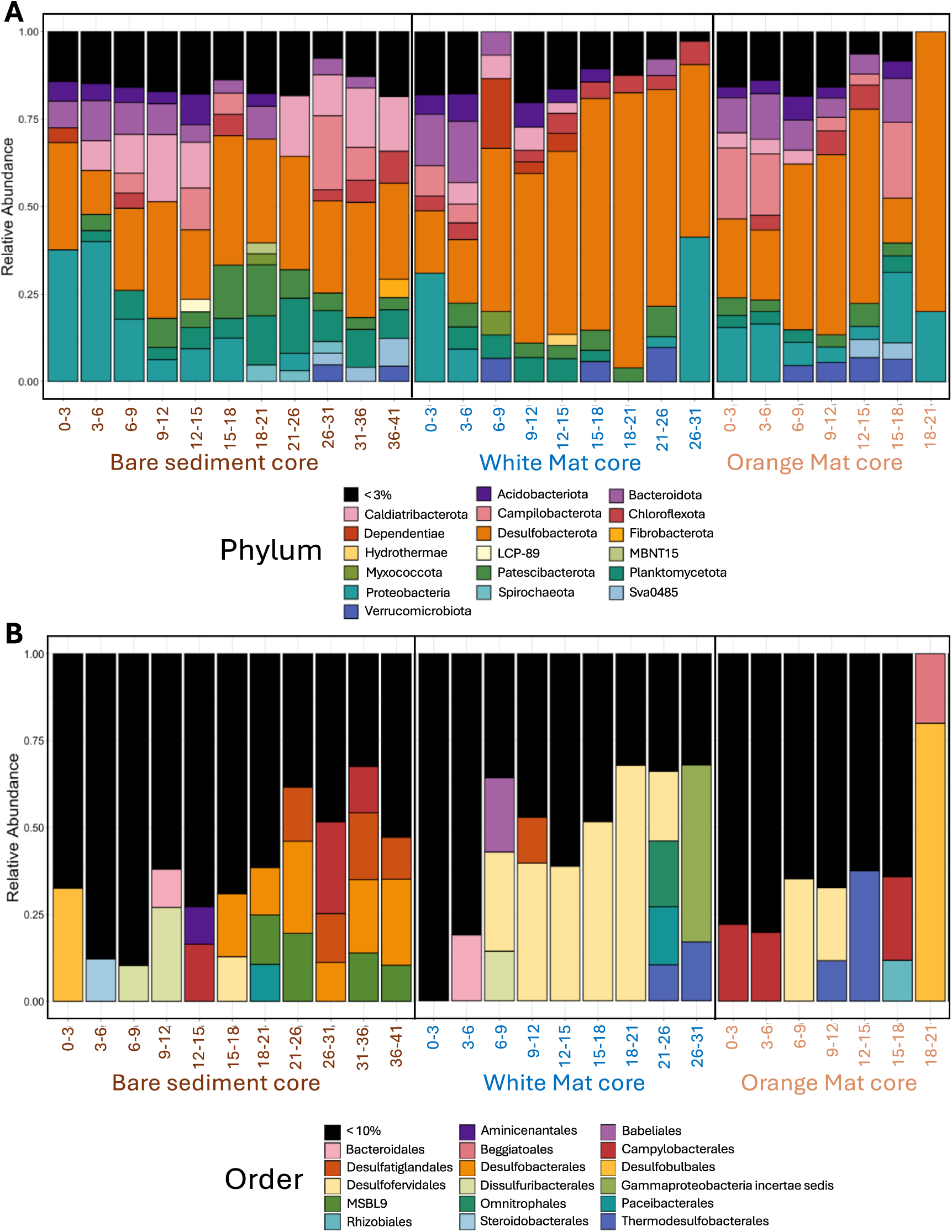
Stacked bar plots of A) phylum-level and B) order-level bacterial diversity. Phyla that comprise more than 3% of total sequence reads and Orders comprising more than 10% of total sequence reads are displayed. These percentage cutoffs were chosen to represent a balance between taxonomic resolution and figure legibility. The X-axis represents depth below seafloor (in centimeters) for the three sediments cores across the mat transect (bare sediment 4872-01, white mat 4872-06, orange mat 4872-14).

Sulfur-cycling lineages that frequently interface with hydrocarbon metabolism are particularly well represented in the sequencing profile and present throughout surficial hydrothermal sediments in the Guaymas Basin (Fig. 3). Sulfur-oxidizing *Beggiatoaceae* assimilate methane-derived carbon (McKay et al., 2012). Numerous sulfate-reducing lineages exist in syntrophic assemblages with methane and alkane oxidizing archaea (Knittel et al., 2003), while other sulfate-reducers perform terminal oxidation of hydrocarbons as free-living organisms (Teske et al., 2019). We performed phylogenetic analyses on these sulfur-cycling lineages to determine their intragroup diversity.

### Beggiatoaceae

The detection of *Beggiatoaceae* is an interesting test of LoopSeq capabilities. Previous efforts to generate full-length *Beggiatoaceae* sequences required collecting large amounts of individually prepared and cleaned filaments (McKay et al., 2012). Due to the size of individual *Beggiatoaceae* cells, they are outnumbered by other much smaller marine bacteria, complicating sequencing efforts and requiring high-throughput methods to overcome the ubiquitous presence of other marine bacterial groups. In our survey, all LoopSeq variants (L-ASV) identified as *Beggiatoaceae* matched reference sequences of the white or orange phylotype, with no additional phylotypes identified (Fig. 4, Suppl. Fig. 4). Light microscopy reveals that past and present sequencing efforts do not capture a complete picture of *Beggiatoaceae* diversity in Guaymas Basin (Salman et al., 2013). There is no evidence for introns within the 16S rRNA genes of Guaymas Basin *Beggiatoaceae,* although they have been observed in non-Guaymas Basin *Beggiatoaceae* (Salman et al., 2011). Recovered L-ASVs were generally found to be associated with surficial sediments underlying white and orange *Beggiatoaceae* mats. Some L-ASVs were found deeper below the surface, which can be attributed to the “Smear effect”, when the coring device pulls some of the long filaments down into the sediment column.

**Figure 4.**
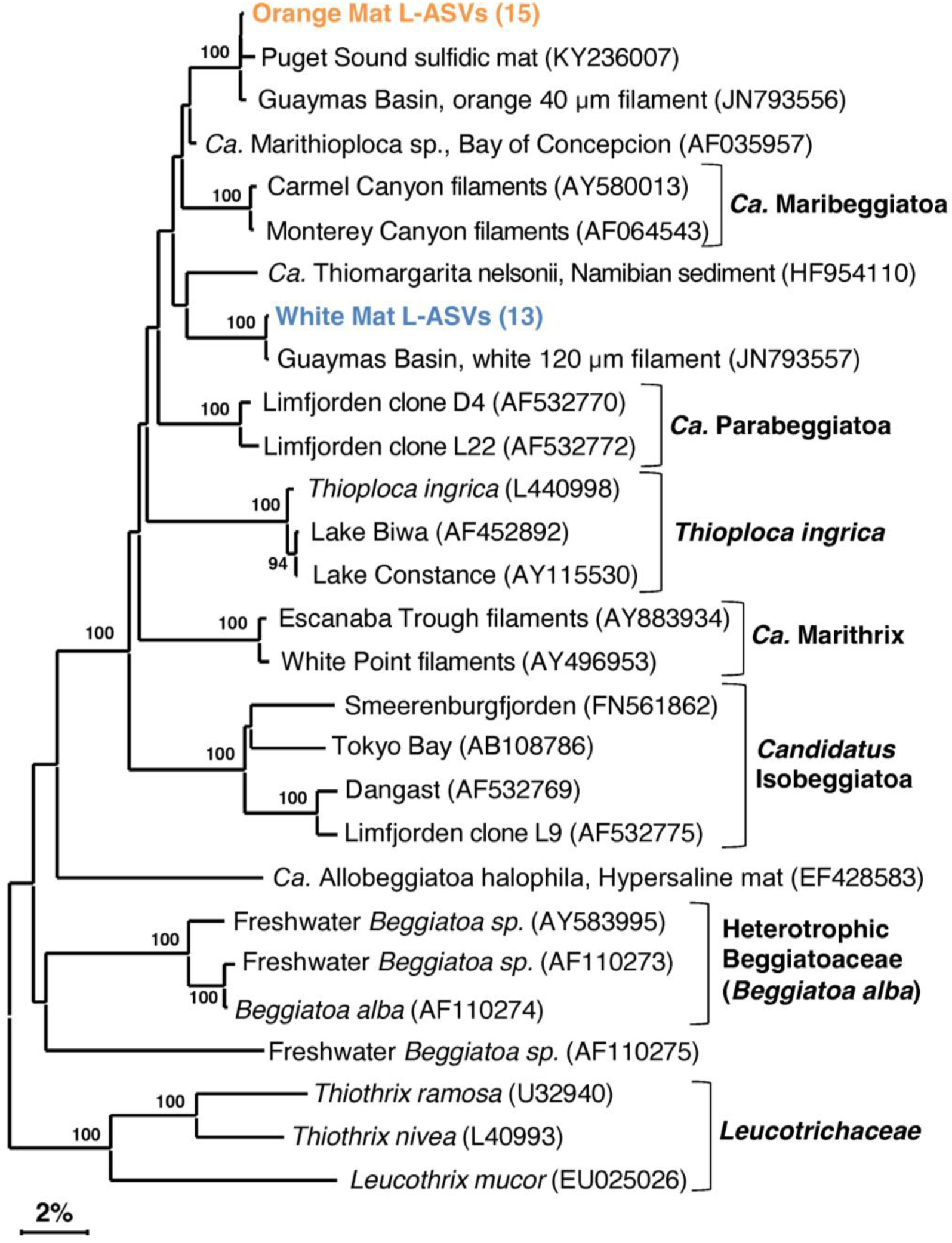
16S rRNA gene sequence based phylogenetic tree of *Beggiatoaceae* and related lineages. Bootstrap values greater than 70% are shown. Two distinct clusters are highlighted, 1) an orange filament cluster composed of L-ASVs from the orange mat site (4872-14) and a reference orange filament sequence obtained from GenBank and 2) a cluster composed of L-ASVs from the white mat site (4872-06) and a reference white filament sequence obtained from GenBank. A version of this tree with expanded L-ASV clusters is available as Supplementary Figure 2.

### *Candidatus* Desulfofervidus

In contrast to *Beggiatoaceae*, *Candidatus* Desulfofervidus sequences are often recovered in Guaymas Basin sequencing surveys (McKay et al., 2016, Dowell et al., 2016). *Ca.* Desulfofervidus is a sulfate-reducing syntrophic partner of methane- and alkane-oxidizing archaea (Laso-Pérez et al., 2016) with a thermal optimum of 50°C (Holler et al., 2011) to 60°C (Kniemeyer et al., 2007, Wegener et al., 2015). Although often observed as a syntroph, *Ca.* Desulfofervidus can also survive as a free-living hydrogenotroph without methane- or alkane-oxidizing partners (Krukenberg et al., 2016). *Ca.* Desulfofervidus is a phylogenetically narrow clade, as it is classified as its own family, order, and class-level lineage within the Phylum *Desulfobacterota* (Waite et al., 2020). However, previous research has not examined the phylogenetic diversity within *Ca.* Desulfofervidus.

Our phylogenies identified several monophyletic well-supported clusters, each with nearly identical 16S rRNA gene sequences. Each cluster contained reference sequences from 50°C (Holler et al., 2011) or 60°C (Kniemeyer et al. 2007, Wegener et al., 2015) enrichments. All reference sequences were from Guaymas Basin, with no close matches from other sites. The close alignment of our sequences with established Guaymas Basin reference sequences attests to their quality. Our phylogeny supports the previous assertion of Krukenberg et al. (2016) that all *Ca.* Desulfofervidus lineages are closely related, since they form a 16S rRNA gene sequence cluster of approximately 98.5 % sequence identity (Suppl. Data File 1, Sheet 2). The five clusters, supported with bootstrap values ≥ 81 % (Fig. 5), imply the presence of unrecognized species diversity within *Ca.* Desulfofervidus.

**Figure 5.**
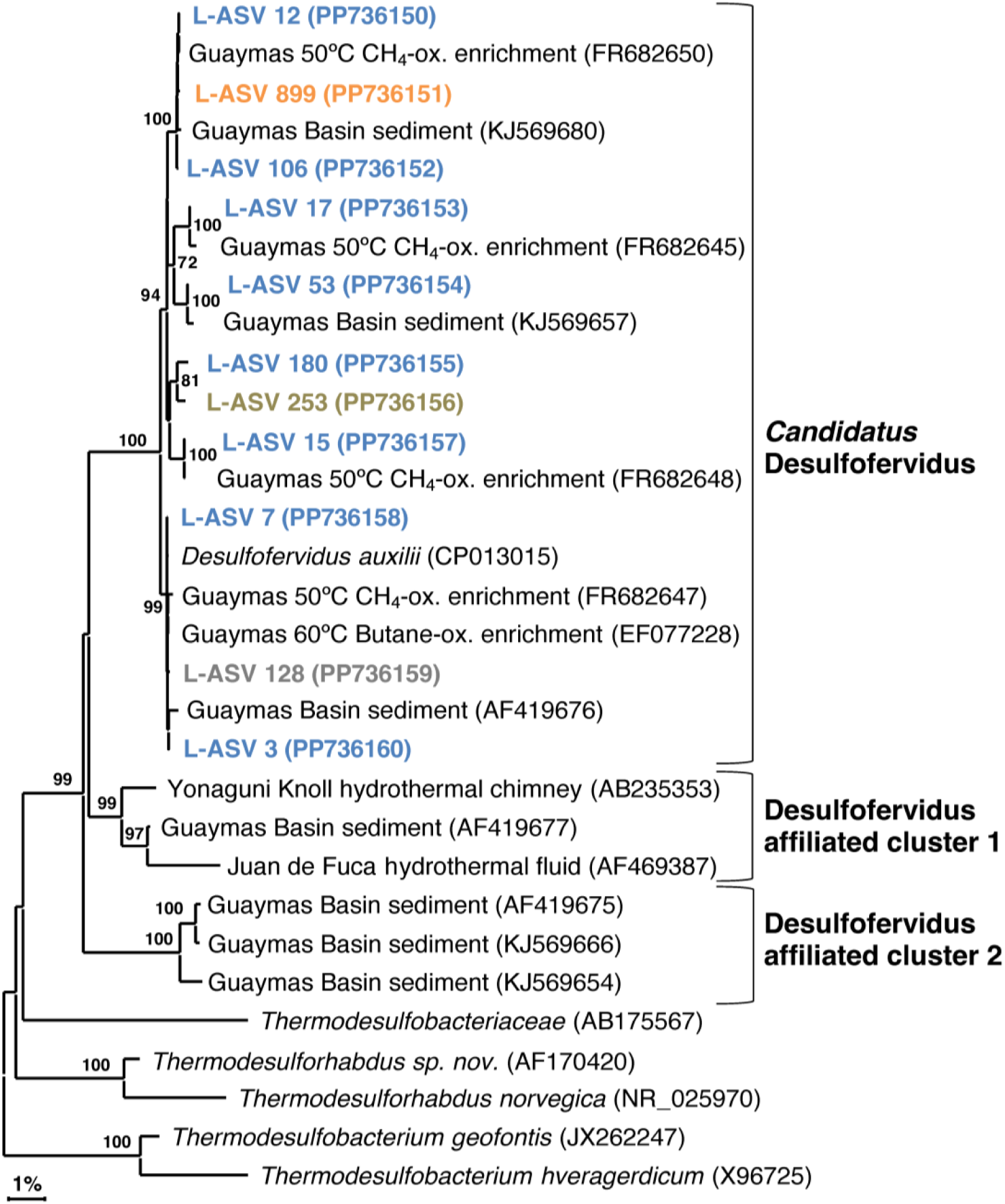
16S rRNA gene sequence based phylogenetic tree of *Candidatus* Desulfofervidus and related lineages. Bootstrap values greater than 70% are shown. NCBI GenBank accession numbers are provided in each tip label. L-ASV tip labels are color coded according to the site where the majority (2/3) of reads were recovered. This phylogenetic tree is based on shorter sequences where the first ∼200 bp were discarded due to chimera issues. **Brown** = bare sediment, 4872-01; **Blue** = white mat site, 4872-06; **Orange** = orange mat, 4872-14; **Grey** = no site had a 2/3+ share of reads.

We found sequences in GenBank that formed two deeply branching clusters with strong bootstrap support that are distinct from the major group of *Ca.* Desulfofervidus (Fig. 5). These clusters were separated from the main group by sequence divergence up to 5.8% (Suppl. Data File 1, Sheet 2). The recovery of these sequences across independent studies at different vent sites suggests unrecognized microbial diversity related to *Ca.* Desulfofervidus. A previous metagenomic survey of Guaymas Basin hydrothermal sediments detected metagenome-assembles genomes (MAGs) branching between *Ca.* Desulfofervidus and *Thermodesulfobacteriaceae* (Dombrowski et al., 2018). Detailed analyses of these MAGs (labeled as B74_G16 and B4_G9) would reveal the genomic content and physiological potential of Desulfofervidus-affiliated lineages.

### SEEP-SRB2

Like *Ca.* Desulfofervidus, SEEP-SRB2 sequences are often recovered in Guaymas Basin sequencing surveys (Dowell et al., 2016). Our survey recovered sequences from slightly below the surface to approximately 15 cmbsf. Unlike *Ca.* Desulfofervidus, SEEP-SRB2 sequences are recovered from diverse non-hydrothermal environments; cold seeps, salt lakes, coastal sediments, and estuaries (Kleindienst et al., 2012, Ruff et al., 2015). Despite its widespread occurrence and likely ecological importance as a syntrophic partner of methane and alkane oxidizers (Krukenberg et al., 2018), the internal phylogenetic structure of this group has not been examined in any detail.

Our SEEP-SRB2 phylogeny is largely comprised of a well-supported major cluster with ∼97.5% sequence identity (Suppl. Data File 1, Sheet 3, equivalent to *Genus* level; Schloss & Handelsman, 2005). The closest cultured relatives of SEEP-SRB2, *Dissulfuribacter thermophilus* and *Dissulfurimicrobium hydrothermale*, are thermophilic sulfur disproportionators isolated from hydrothermal environments (Slobodkin et al., 2013, Slobodkin et al., 2016). These bacteria constitute separate lineages but share a deep branching point with SEEP-SRB2 (Fig. 6). The distinctiveness of these lineages is further supported by sequence distances of up to 11% (Suppl. Data File 1, Sheet 3). Our 16S rRNA gene sequences indicate that SEEP-SRB2 would form a stand-alone group within *Dissulfuribacteraceae* (Waite et al., 2020). Further phylogenetic diversity that would fall between the pure culture examples (*Dissulfuribacter thermophilus* and *Dissulfurimicrobium hydrothermale*) and SEEP-SRB2 remains to be explored. Previously, the creation of a taxonomic family to comprise *Dissulfuribacter thermophilus* and *Dissulfurimicrobium hydrothermale* has been suggested (Slobodkin et al., 2013). Our phylogeny shows that such a group would also include SEEP-SRB2 (Fig. 6).

**Figure 6.**
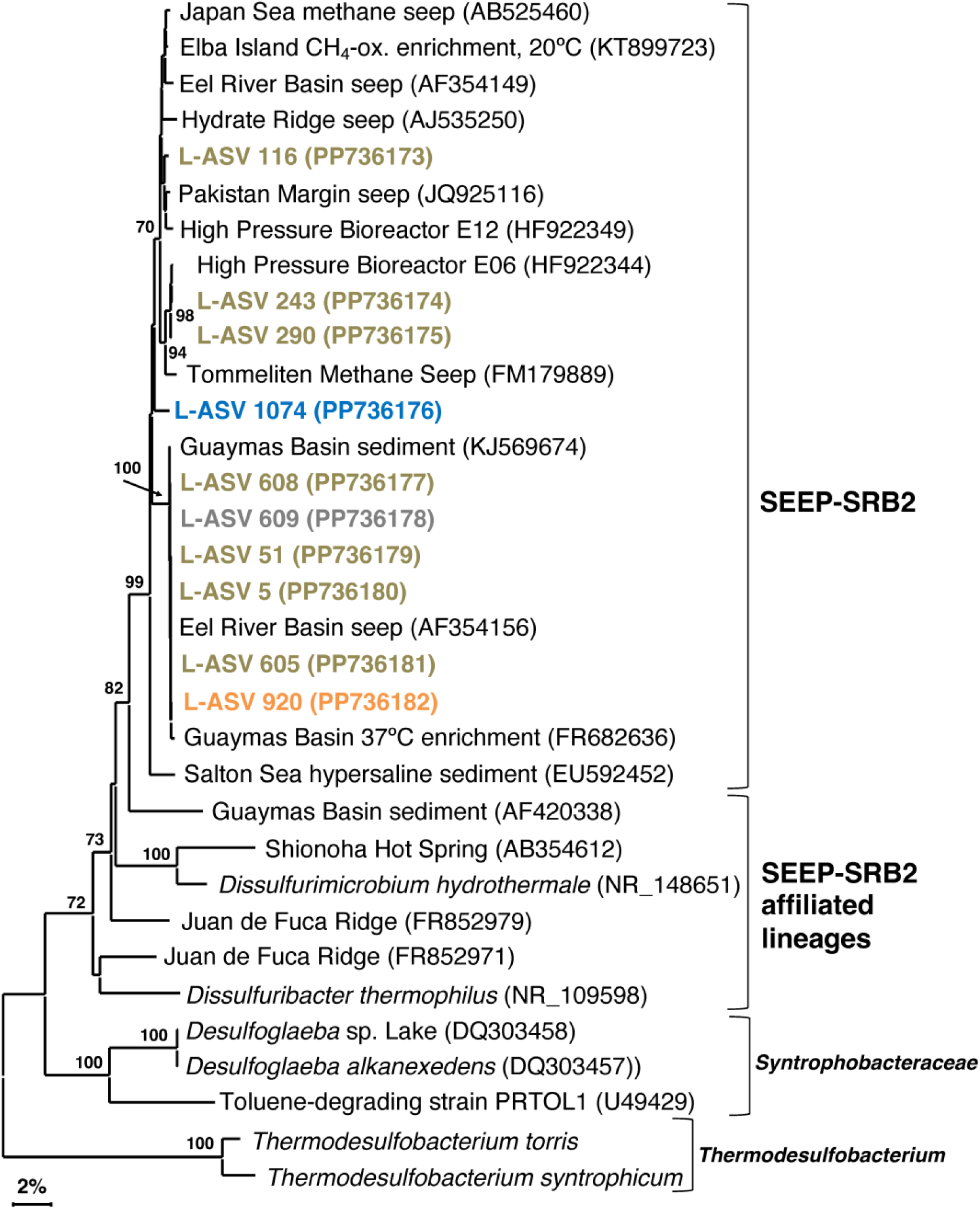
16S rRNA gene sequence based phylogenetic tree of SEEP-SRB2 and related lineages. Bootstrap values greater than 70% are shown. NCBI GenBank accession numbers are provided in each tip label. L-ASV tip labels are color coded according to the site where the majority (2/3) of reads were recovered. This phylogenetic tree is based on shorter sequences where the first ∼200 bp were discarded due to chimera issues. **Brown** = bare sediment, 4872-01; **Blue** = white mat site, 4872-06; **Orange** = orange mat, 4872-14; **Grey** = no site had a 2/3+ share of reads. *Thermodesulfobacterium torris 16S rRNA sequence was obtained from an annotated genome (BioSample ID: SAMN27514933). *Thermodesulfobacterium syntrophicum 16S rRNA sequence was obtained from an annotated genome (BioSample ID: SAMN29995626).

### SEEP-SRB4

In contrast to the syntrophic symbionts SEEP-SRB1 and SEEP-SRB2, SEEP-SRB4 has largely been overlooked since it was proposed 20 years ago (Knittel et al., 2003). FISH surveys indicate that SEEP-SRB4 is not a syntroph, but occurs as individual free-living cells (Knittel et al., 2003). The closest cultured relatives of SEEP-SRB4 are non-thermophilic sulfur-disproportionating genera such as *Desulforhopalus* (Isaksen & Teske, 1996), *Desulfocapsa* (Finster et al., 1998), *Desulfofustis* (Friedrich et al., 1996), and the sulfate-reducing psychrophile *Desulfotalea* (Knoblauch et al., 1999). While SEEP-SRB2 has been enriched from Guaymas Basin sediments at 37°C (Krukenberg et al., 2018), there is currently no evidence that SEEP-SRB4 is moderately thermotolerant. We seek to test this assumption in our survey, checking the distribution of SEEP-SRB4 in surficial hydrothermal sediments. As with SEEP-SRB2, we seek to investigate the previously unrecognized phylogenetic diversity of SEEP-SRB4.

Our SEEP-SRB4 phylogeny depicts one major cluster with strong bootstrap support (100 %) and > 98.9% sequence identity (Suppl. Data File 1, Sheet 4). This cluster contains several well-supported, phylogenetically narrow clusters (Fig. 7). While the phylogeny shows a potential Guaymas Basin-specific subcluster of four sequences, the other subclusters contain globally distributed sequences from diverse cold seep sites and from Guaymas Basin.

**Figure 7.**
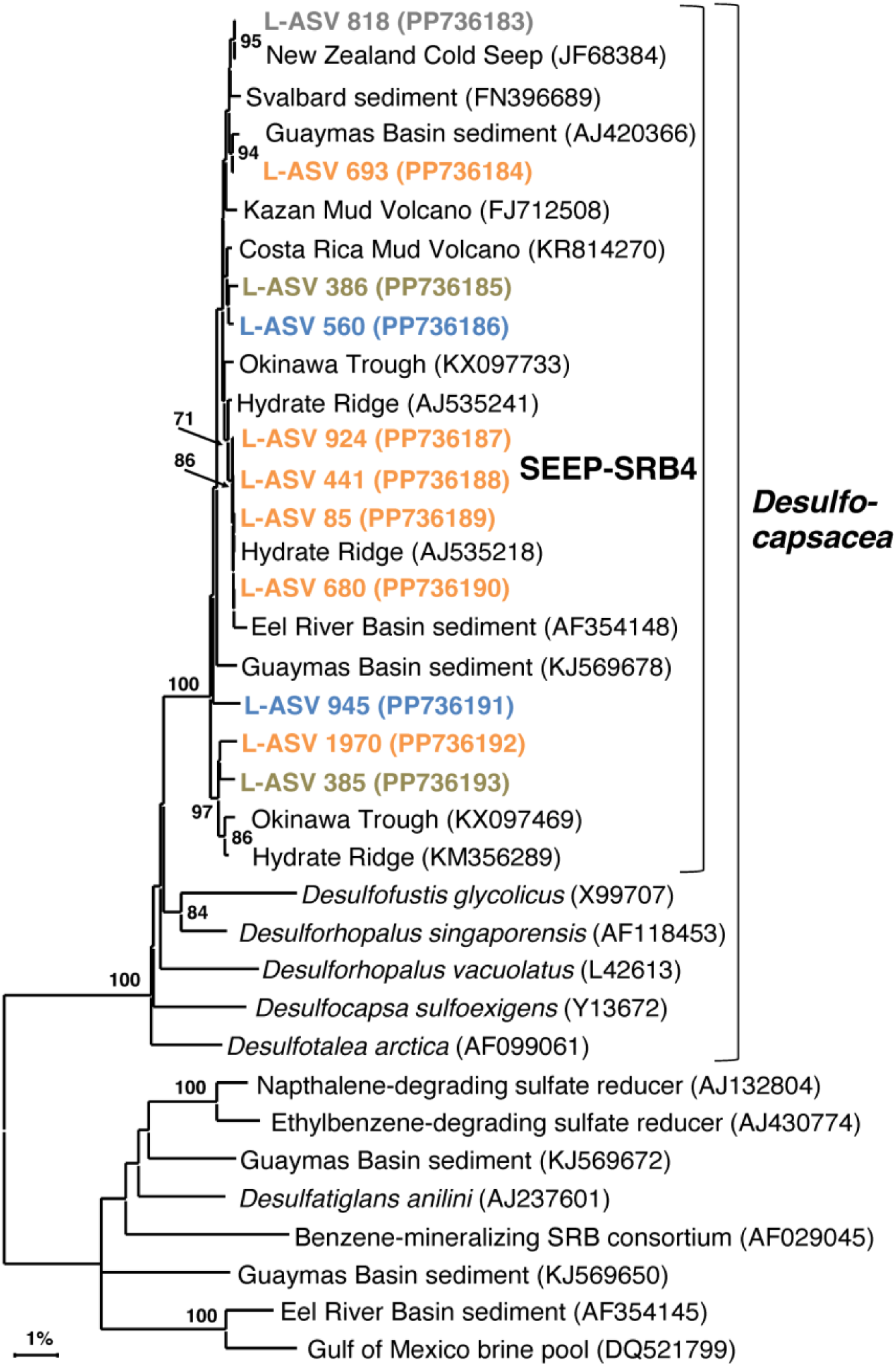
16S rRNA gene sequence based phylogenetic tree of SEEP-SRB4 and related lineages. Bootstrap values greater than 70% are shown. NCBI GenBank accession numbers are provided in each tip label. L-ASV tip labels are color coded according to the site where the majority (2/3) of reads were recovered. This phylogenetic tree is based on shorter sequences where the first ∼200 bp were discarded due to chimera issues. **Brown** = bare sediment, 4872-01; **Blue** = white mat site, 4872-06; **Orange** = orange mat, 4872-14; **Grey** = no site had a 2/3+ share of reads.

The majority of L-ASVs for SEEP-SRB4 are retrieved from the top ∼6 cm of the sediment across the entire lateral hydrothermal gradient (Suppl. Table 3). The preference for surficial sediments may be due to the abundance of elemental sulfur and polysulfides located there (Teske et al., 2016). A similar preference for surficial sediments with coexisting sulfate and sulfide was noted in the first study of SEEP-SRB4 (Knittel et al., 2003). SEEP-SRB4 bacteria seem to be able to colonize surficial hydrothermal sediments as long as in-situ temperatures remain relatively cool, similar to the strategy utilized by Guaymas Basin *Beggiatoaceae* (McKay et al., 2012).

### Desulfatiglans

*Desulfatiglandales* is an order-level lineage within the phylum *Desulfobacterota* that contains polyaromatic degrading organisms, both isolated and uncultured species. The order and family level taxonomy are named after *Desulfatiglans annilini*, the first cultured representative (Suzuki et al., 2014). Sequences related to *Desulfatiglans* are often recovered in Guaymas Basin sequencing surveys (Ramírez et al., 2021, Dowell et al., 2016), including the shallow subsurface (Edgcomb et al., 2022). In contrast to the tightly clustered groups of SEEP-SRB2 and SEEP-SRB4, *Desulfatiglans* exhibits a greater degree of phylogenetic diversity.

Our Guaymas Basin sequences and almost all GenBank sequences were not closely related to cultured isolates that degrade polyaromatic and substituted-aromatic compounds (Fig. 7). Even finding uncultured sequences that were closely related (>98% sequence identity) to some of our Guaymas Basin ASVs proved to be difficult. This could imply an undersampling of the natural diversity of *Desulfatiglans*, or their ongoing diversification and speciation. Previously, *Desulfatiglans* has been classified into two groups distinguished by genomic and physiological differences, Group 1 consisting of sulfate reducers and Group 2 consisting of dehalogenating subsurface bacteria (Jochum et al., 2018). Most of our L-ASVs belong to Group 1 and a few belong to Group 2.

Outside of these two groups, we found two well-supported hydrothermal clusters (Fig. 8). The first cluster contained L-ASV 190 (predominately recovered from the white mat site), sequences from 37 °C and 50 °C enrichments of Guaymas Basin sediment (Kellermann et al., 2012, Wegener et al., 2015) and an environmental sequence from Guaymas Basin hydrothermal sediment (McKay et al., 2016). The second cluster contained L-ASV 207 (predominately recovered from the orange mat site), a sequence from a *Desulfatiglans* MAG that was prevalent in a 70 °C enrichment of Guaymas Basin sediment (Zehnle et al., 2023), and environmental sequences from diverse hydrothermal sediments (Li et al., 2014, Li et al., 2020). We examined these sequences via distance matrix analyses and find that these hydrothermal clusters are generally separated from other sequences by 11-12% sequence divergence (Suppl. Data File 1, Sheet 5).

**Figure 8.**
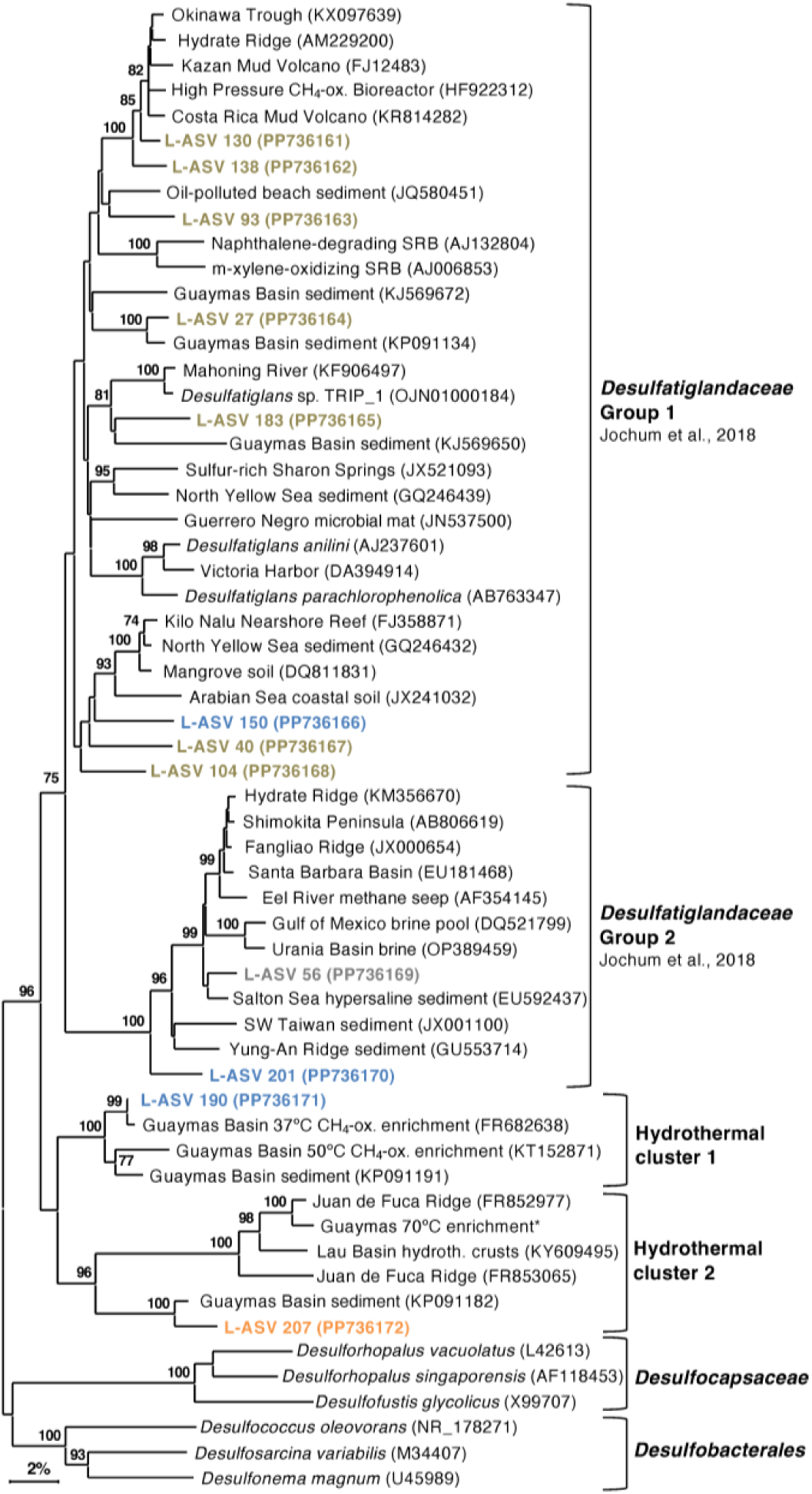
16S rRNA gene sequence based phylogenetic tree of *Desulfatiglans* and related lineages. Bootstrap values greater than 70% are shown. NCBI GenBank accession numbers are provided in each tip label. L-ASV tip labels are color-coded according to the site where the majority (2/3) of reads were recovered. This phylogenetic tree is based on shorter sequences where the first ∼200 bp were discarded due to chimera issues. **Brown** = bare sediment, 4872-01; **Blue** = white mat site, 4872-06; **Orange** = orange mat, 4872-14; **Grey** = no site had a 2/3+ share of reads. The Guaymas 70°C enrichment sequence, marked with an asterisk (*), is from MAG 9 (B70, Benzene enrichment at 70°C) under BioProject PRJNA1013425.

In contrast to other sulfate reducers that occur in warm and hot Guaymas Basin sediments, we observed most of our *Desulfatiglans* L-ASVs in cool sediments, suggesting that they do not depend on methane and light hydrocarbons derived from hydrothermal fluids, and instead utilize polyaromatics (Schnell & Schink, 1991, Zehnle et al., 2023). Polyaromatics from hydrothermal oils are prevalent throughout Guaymas Basin sediments of diverse thermal regimes (Kawka & Simoneit, 1990), and *Desulfatiglans* appears to have evolved to utilize polyaromatics across the thermal spectrum, as suggested by its hydrothermal clusters.

### Chimera Issues

DADA2 detected chimeras in approximately 4% of our reads from a natural hydrothermal community. Previous research found structural errors (chimeras and introgressions) in approximately 2% of reads from a Zymo mock community (Callahan et al., 2021). Specifically, we found that LoopSeq produced chimeras in the majority of *Candidatus* Desulfofervidus sequences we examined. Segments of these sequences were entered into BLASTn, and we discovered that the chimeras consistently occurred in the first ∼200 bp. BLASTn searches of the chimeric segments revealed their relation to other marine sediment bacteria such as *Chloroflexi*. The chimera breaks are located within the highly variable Helix 11 of the 16S rRNA gene (Van de Peer, et al. 1996), and therefore appear to be secondary structure dependent. These issues with *Ca.* Desulfofervidus sequences were observed in samples across the DNA concentration range, from very high (4240 ng/cm^3^) to very low (0.5 ng/cm^3^). Automated chimera identifying programs can have sensitivity issues when used with uncultured lineages that have poorly resolved phylogenies. We found that trimming off the first ∼200 bp remedied the chimera issue, resulting in sequences with high alignment quality and >97% identity to previously archived *Ca.* Desulfofervidus sequences available in GenBank.

LoopSeq recovered eight sequences identified by the DADA2 annotation pipeline as “*Thermodesulfobacterium*”. Initial analysis of theses sequences showed that they were plagued with chimera and artifact issues, a potential consequence of low DNA yields from deep, hot samples. Two sequences were salvaged; one did not have chimera issues (L-ASV 12240), while the other was trimmed to only include the first ∼340 bp (L-ASV 11484). These two sequences came from samples with low DNA concentrations (Sample 4872-06-08, 50 ng/cm^3^ and Sample 4872-06-09, 12 ng/cm^3^). These two sequences formed a Guaymas Basin specific cluster (Suppl. Fig. 5) with the thermophilic ANME-1 syntroph *Thermodesulfobacterium torris* (Benito Merino et al., 2022) and a nearly identical hydrothermal sediment clone (McKay et al., 2016).

### Conclusions

Our sequencing survey has shown that LoopSeq is a promising option for the high throughput long read sequencing of diverse microbial communities in hydrothermal sediments. The huge quantity of sequences yielded by LoopSeq allowed us to further resolve the fine scale phylogenetic structure of sulfur-cycling bacterial groups that are characteristic of Guaymas Basin hydrothermal sediments. The five well-supported clusters revealed in our *Ca.* Desulfofervidus phylogeny may represent unrecognized physiological and genomic diversity that calls for further investigation beyond the initial description of this group (Krukenberg et al., 2016). Investigating SEEP-SRB2, SEEP-SRB4, and *Ca.* Desulfofervidus showed that they are all phylogenetically distinct groups without a spectrum of close relatives among environmental clones as represented in NCBI GenBank (Figs. 5, 6, & 7). SEEP-SRB2, SEEP-SRB4, and *Ca*. Desulfofervidus all contain considerable internal phylogenetic diversity within their respective clades. Sulfur-disproportionating species and genera are among the closest cultured relatives of SEEP-SRB2 and SEEP-SRB4 (Figs. 6 & 7). In our distance analyses we see sequences distances of up to ∼8% between sulfur-disproportionating bacteria and SEEP-SRB2 and up to ∼5% between sulfur-disproportionating bacteria and SEEP-SRB4 (Suppl. Data File 1, Sheets 3 & 4). The near-full-length sequences generated by LoopSeq enabled these distance analyses. Elemental sulfur and sulfur species of intermediate oxidation states are abundant at seep sites (Jørgensen et al., 2019). This provides a favorable environment for sulfur-disproportionating bacteria, and for the diversification of sulfur-oxidizing and sulfate-reducing bacterial lineages.

However, the chimera issues that we detected in sequences identified as *Ca.* Desulfofervidus, *Thermodesulfobacterium*, and *Beggiatoaceae*, underscore the need for phylogenetic analyses and alignment comparisons with high quality reference sequences to confirm the quality and identity of LoopSeq sequences, and the reliability of microbial community analyses. We also note that our DADA2 analysis of these sequences was not sufficient to detect these chimeras. Without manual verification of sequence alignments and scrutiny of isolated distal branches in phylogenetic clusters, problematic sequences can go undetected, and create artificial diversity. Our observation of lineage-specific chimera issues calls for caution when using LoopSeq alone to investigate extreme and unusual bacterial groups. Since we submitted our samples for sequencing (early 2022), Element Biosciences has instituted changes with the handling of low-biomass samples (such as our own) and has adapted the chemistry of LoopSeq to improve DNA yield in low biomass samples (Element Biosciences representative Dr. Caroline Obert, personal communication). Further protocol development of LoopSeq should monitor this issue, and check sequence accuracy among a wide range of microbial target groups.

LoopSeq has successfully enabled a survey of the diversity of sulfur-cycling bacteria across a thermal and substrate gradient up to the current limit of hydrocarbon degradation (Fig. 9). Due to the scope of this study, the diversity of archaea in surficial Guaymas Basin sediments was not investigated with LoopSeq. Further work may use LoopSeq to investigate potentially unknown archaeal diversity in habitats with extreme conditions or along environmental gradients (e.g., Guaymas Basin). Phylogenetic resolution and range of archaeal lineages is limited by sequence coverage. Due to the limitations of popular primers, most available archaeal 16S rRNA gene sequences are 800 bp in length or less (Teske & Sørensen, 2008). High quality near full-length 16S rRNA gene sequence data, such as that provided by LoopSeq, would allow for an improved view of archaeal diversity.

**Figure 9.**
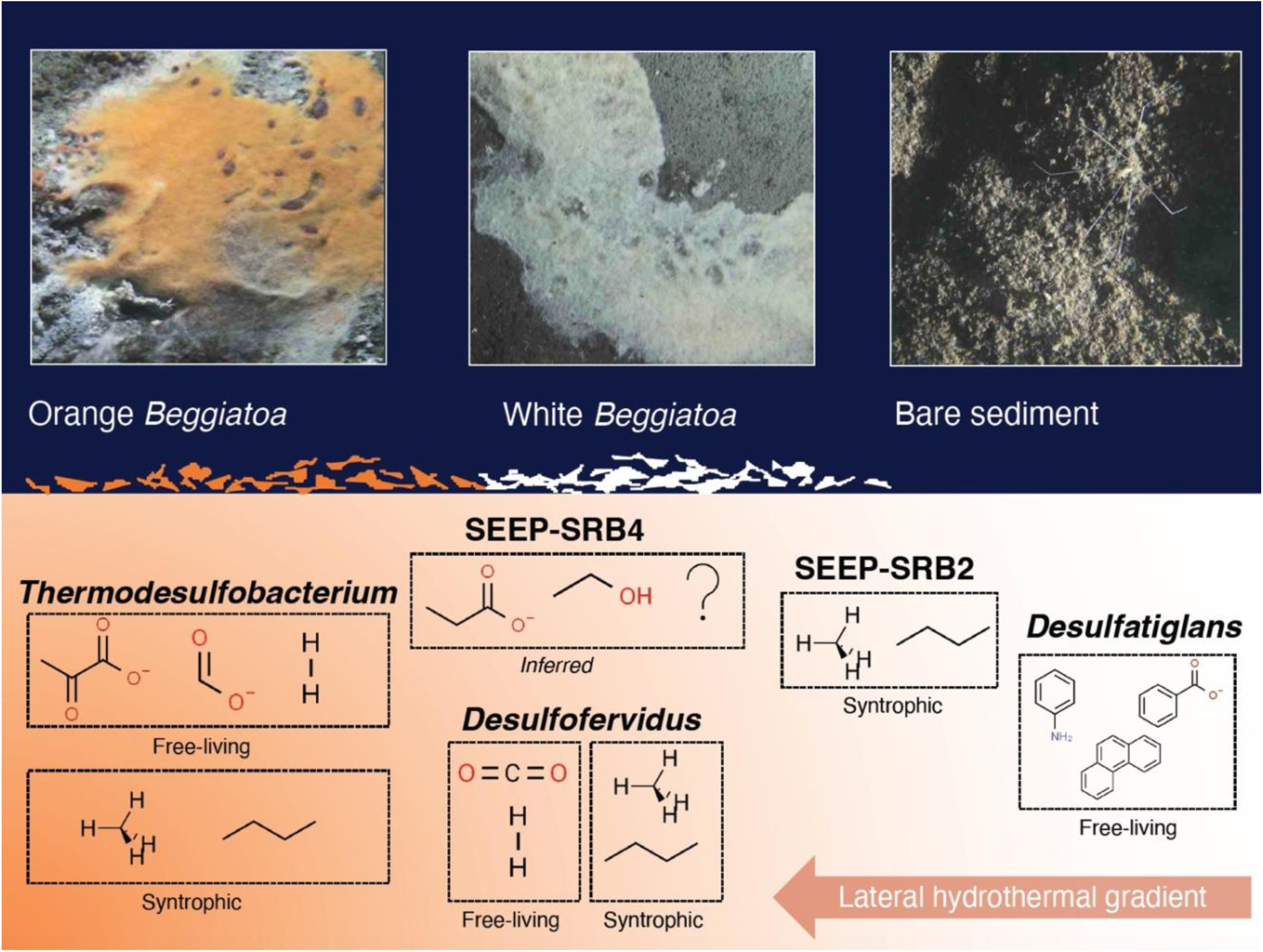
Conceptual figure of sulfur-cycling bacterial groups (with examples of preferred substrates) and their approximate distribution along a lateral hydrothermal gradient in surficial Guaymas Basin sediments. Examples of preferred substrates are shown for both free-living and syntrophic lifestyles, when applicable. Molecular structure images of the preferred substrates were obtained from chemspider.com. *Beggiatoaceae* mat and sediment close up images were captured with the bottom facing camera of HOV *Alvin* during Dive 4872.

## Supporting information

Element Biosciences 16S LoopSeq User Guide

DNA Extraction Protocol

## Data Availability Statement

All sequence reads were submitted to the NCBI Sequence Read Archive (BioProject: PRJNA1105367). L-ASV sequences used in our phylogenetic trees are available in Supplementary Data File 2 or via their NCBI GenBank accession number provided in each phylogenetic tree.

## Author Contributions

JEH analyzed the sequence data, created the figures, isolated the DNA, and wrote the manuscript. JPC performed the methane concentration and stable isotope analyses. MAM wrote the DADA2 script used to analyze the sequence data and prepared samples for submission. SER prepared samples for submission and handled interactions with Element Biosciences over time and advocated on the group’s behalf. AT obtained the samples during Alvin Dive 4872, guided the interpretation of the phylogenetic trees and geochemical data, supported the development of the manuscript, and taught JEH the DNA extraction method. All authors contributed to critical revisions of the manuscript.

## Acknowledgements

We thank the R/V *Atlantis* and HOV *Alvin* crews for their exemplary support at sea, especially *Alvin* pilot Jefferson Grau for expert handling of the craft. We thank Chris Chambers for conducting the sulfide analyses, and Catherine Crowley for clean-up and quantification of the DNA. We thank Element Biosciences for their patience with our complex environmental samples.

