## Supplementary material for "Complex bacterial diversity of Guaymas Basin hydrothermal sediments revealed by synthetic long-read sequencing (LoopSeq)": Element Biosciences 16S LoopSeq User Guide

LoopSeq

### 16S LoopSeq™

#### User Guide

**FOR USE WITH**

16S LoopSeq, catalog # 840-10001
Extension LoopSeq, catalog # 840-10003

**ELEMENT BIOSCIENCES**
**For Research Use Only. Not for Use in diagnostic procedures.**

Document # MA-00027 Rev. D
December 2023

For Research Use Only. Not for use in diagnostic procedures. Information in this document is subject to change without notice. Certain terms, conditions, and/or use restrictions may pertain to your use of products of Element Biosciences, Inc. and its affiliates (collectively, “Element”). Please refer to any applicable terms and conditions.

ELEMENT DISCLAIMS ALL WARRANTIES WITH RESPECT TO THIS DOCUMENT, EXPRESS, STATUTORY, IMPLIED OR OTHERWISE, INCLUDING, BUT NOT LIMITED TO, ANY WARRANTIES OF MERCHANTABILITY, SATISFACTORY QUALITY, NON-INFRINGEMENT OR FITNESS FOR A PARTICULAR PURPOSE. IN NO EVENT SHALL ELEMENT BE LIABLE, WHETHER IN CONTRACT, TORT, WARRANTY, PURSUANT TO ANY STATUTE, OR ON ANY OTHER BASIS FOR SPECIAL, CONSEQUENTIAL, INCIDENTAL, EXEMPLARY OR INDIRECT DAMAGES IN CONNECTION WITH THIS DOCUMENT, WHETHER OR NOT FORESEEABLE AND WHETHER OR NOT ELEMENT BIOSCIENCES IS ADVISED OF THE POSSIBILITY OF SUCH DAMAGES.

© 2023 Element Biosciences, Inc. All rights reserved. Element Biosciences, Loop Genomics, and associated logos are trademarks of Element Biosciences, Inc. Products described herein may be manufactured under or covered by one or more pending or issued US or foreign patents. Visit elementbiosciences.com/legal/patents for more information about Element’s patents and trademarks. Other names mentioned herein may be trademarks of their respective companies.

CHAPTER 1

### Library Prep Overview

### Introduction

The 16S LoopSeq Workflow pairs the LoopSeq 16S Kit and LoopSeq Core Construction Kit to prepare single-index libraries for sequencing on an Illumina system. The workflow supports preparation of up to 96 samples.

The workflow starts with enrichment, which captures targets of interest. PCR then adds unique molecular identifiers (UMIs) and a LoopSeq index, barcoding the samples. Single-sample calibration (SSC) dilutes the barcoded samples, which are amplified and pooled for streamlined library prep. Library prep starts with the random distribution of UMIs throughout the samples and generation of blunt-ended DNA fragments. Adapter ligation adds primers for multiplex sequencing. Bead-based cleanups maintain purity.

**Figure 1:** Structure of a LoopSeq library

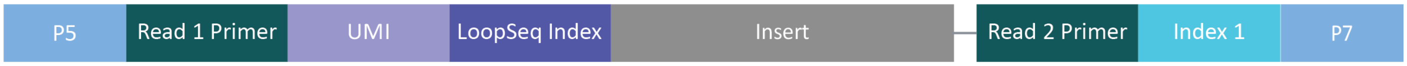

#### Sequence Tags

16S LoopSeq tags each sample with three types of sequences:

- A UMI identifies each molecule in a sample.
- A LoopSeq index identifies each sample.
- An index sequence identifies the pool, which contains an equal volume of each sample in the prep.

#### Product Configuration

The purchase of 16S LoopSeq includes a 16S kit and a core construction kit. Element also offers the core construction kit as an extension.

| Product | Kits | Requirement |
|---|---|---|
| 16S LoopSeq | LoopSeq 16S Kit | Required |
| | LoopSeq Core Construction Kit | Required |
| Extension LoopSeq | LoopSeq Core Construction Kit | Optional |

#### Safety Data Sheets

When using the LoopSeq 16S Kit, LoopSeq Core Construction Kit, and other reagents, always wear personal protective equipment (PPE): a lab coat, powder-free disposable gloves, and protective goggles. Review the safety data sheets (SDS) for chemical properties. The SDS inform safety, disposal, and hazards for your region and are available at elementbiosciences.com/resources.

### Workflow Summary

Two protocols comprise the 16S LoopSeq Workflow: a barcode protocol uses the LoopSeq 16S Kit to structure molecules and a library prep protocol uses the LoopSeq Core Construction Kit to generate libraries.

The barcode protocol takes ~9 hours, including ~2.5 hours of hands-on time. Library prep takes ~7 hours, including ~2.5 hours of hands-on time.

**Figure 2:** Protocol steps and reagents

| | Procedure | Kit Reagents | |
|---|---|---|---|
| 1 | **Enrich samples** | Sample Enrichment Mix | Start with the LoopSeq 16S Kit |
| 2 | **Amplify enriched samples** | 16S SSC Barcoding Plate 96-Plex | |
| 3 | **Clean up samples** | | |
| Safe Stopping Point | | | |
| 4 | **Calibrate barcodes** | | |
| 5 | **Amplify and pool samples** | Amplification Mix B | |
| 6 | **Clean up sample pool** | | |
| Safe Stopping Point | | | Switch to the LoopSeq Core Construction Kit |
| 7 | **Distribute barcodes** | Distribution Enzyme, Distribution Mix | |
| 8 | **Activate and neutralize barcodes** | Activation Enzyme, Activation Mix, Neutralization Enzyme | |
| 9 | **Clean up activated samples** | | |
| Safe Stopping Point | | | |
| 10 | **Fragment samples and repair ends** | Fragmentation Enzyme, Fragmentation Mix | |
| 11 | **Ligate adapters** | Ligation Enzyme, Ligation Mix | |
| 12 | **Clean up library** | | |
| Safe Stopping Point | | | |
| 13 | **Amplify library** | Index Master Mix, Index Primer P1–P4 | |
| 14 | **Clean up indexed library** | | |

Barcode Protocol (steps 1–6) Library Prep Protocol (steps 7–14)

### Kit Contents and Storage

Each kit is packaged in one box and shipped on dry ice. When you receive your kits, promptly store the contents at the proper temperature. Reference reagent labels for fill volumes.

#### LoopSeq 16S Kit

| Reagent | Quantity | Cap Color | Storage Temperature |
|---|---|---|---|
| 16S SSC Barcoding Plate 96-Plex | 1 | Not applicable | -30°C to -10°C |
| Amplification Mix B | 2 | Orange | -30°C to -10°C |
| Sample Enrichment Mix | 1 | Red | -30°C to -10°C |

#### LoopSeq Core Construction Kit

| Reagent | Quantity | Cap Color | Storage Temperature |
|---|---|---|---|
| Activation Enzyme | 1 | Red | -30°C to -10°C |
| Activation Mix | 1 | Red | -30°C to -10°C |
| Distribution Enzyme | 1 | Red | -30°C to -10°C |
| Distribution Mix | 1 | Red | -30°C to -10°C |
| Fragmentation Enzyme | 1 | Orange | -30°C to -10°C |
| Fragmentation Mix | 1 | Orange | -30°C to -10°C |
| Index Master Mix | 1 | Orange | -30°C to -10°C |
| Index Primer P1 | 1 | Orange | -30°C to -10°C |
| Index Primer P2 | 1 | Orange | -30°C to -10°C |
| Index Primer P3 | 1 | Orange | -30°C to -10°C |
| Index Primer P4 | 1 | Orange | -30°C to -10°C |
| Ligation Enzyme | 1 | Orange | -30°C to -10°C |
| Ligation Mix | 1 | Orange | -30°C to -10°C |
| Neutralization Enzyme | 1 | Red | -30°C to -10°C |

### User-Supplied Materials

In addition to the LoopSeq kits, the workflow requires the third-party consumables and equipment listed in the following tables. The consumables table includes forward and reverse primer sequences, which are modifications of the typical 27F and 1429R primer sequences.

#### User-Supplied Consumables

| Supplier | Consumable | Catalog # |
|---|---|---|
| General lab supplier | 96-well PCR plates | Not applicable |
| | Absolute ethanol | Not applicable |
| | Filtered pipette tips | Not applicable |
| | Microcentrifuge tubes, 1.5–5 ml | Not applicable |
| | PCR tubes, strip or individual | Not applicable |
| | [Optional] Amber microcentrifuge tubes, 1.5 ml | Not applicable |
| | [Optional] 27F primer<br>5′–AGAGTTTGATCMTGGCTCAG–3′ | Not applicable |
| | [Optional] 1429R primer<br>5′–TACCTTGTTACGACTT–3′ | Not applicable |
| Agilent | Agilent High Sensitivity DNA Kit | Part # 5067-4626[1] |
| Beckman Coulter | Either bead type:<br>• AMPure XP Reagent, 60 ml<br>• SPRIselect, 5 ml or 60 ml | The corresponding catalog #:<br>• A63881<br>• B23317 or B23318 |
| Bio-Rad | Microseal 'B' Film, adhesive | Catalog # MSB1001[1] |
| New England Biolabs (NEB) | Q5 High-Fidelity DNA Polymerase | Catalog # M0491S or M0491L |
| Qiagen | Buffer EB, 250 ml | Catalog # 19086[2] |
| Roche | KAPA Library Quantification Kit - Complete kit (Universal) | Catalog # 07960140001[1] |

| Supplier | Consumable | Catalog # |
|---|---|---|
| Thermo Fisher Scientific | 96-well 0.8 ml deepwell storage plates | Catalog # AB0765[1] |
| | Either quantification kit:<br>• Qubit 1X dsDNA HS Kit<br>• Qubit dsDNA HS Assay Kit | The corresponding catalog #:<br>• Q33230 or Q33231<br>• Q32851 or Q32854 |
| | [Optional] SYBR Green I Nucleic Acid Gel Stain | Catalog # S7563, S7585, or S7567 |
| | [Optional] E-Gel CloneWell II Agarose Gels with SYBR Safe, 0.8% | Catalog # G661818 |
| VWR | Nuclease-free water, 500 ml | Catalog # 97062-794[1] |
| Zymo | [Optional] HostZERO Microbial DNA Kit | Catalog # D4310[1] |

[1] Consumables that you have tested and demonstrate equivalent performance are acceptable.

[2] User-prepared 10 mM Tris Buffer, pH 8.5 is an acceptable substitute.

### User-Supplied Equipment

| Supplier | Equipment | Catalog # |
|---|---|---|
| General lab supplier | Centrifuge, multipurpose | Not applicable |
| | Ice bucket | Not applicable |
| | Pipettes, single- and multi-channel | Not applicable |
| | Vortex mixer | Not applicable |
| Agilent | 2100 Bioanalyzer Instrument | Catalog # G2939BA[1] |
| Bio-Rad | Any quantitative PCR (qPCR) system:<br>• CFX Opus 96 Real-Time PCR Instrument<br>• CFX Opus 384 Real-Time PCR System<br>• CFX Connect Real-Time System | The corresponding catalog #:<br>• 12011319[1]<br>• 12011452[1]<br>• 1855201[1] |
| | Either thermal cycler:<br>• C1000 Touch Thermal Cycler<br>• T100 Thermal Cycler | The corresponding catalog #:<br>• Catalog # 1851197[1]<br>• Catalog # 1861096[1] |
| Permagen | PCR Strip Magnetic Separator, 0.2 ml, 8- or 12-Strip | SKU # MSR812[1,2] |
| QInstruments | BioShake XP | Order # 1808-0505[1] |

| Supplier | Equipment | Catalog # |
| --- | --- | --- |
| Thermo Fisher Scientific | MagJET Separation Rack, 12 x 1.5 ml tube | Catalog # MR02[2] |
| | Magnetic Stand-96 | Catalog # AM10027[2] |
| | Either fluorometer:<br>• Qubit 3 Fluorometer<br>• Qubit 4 Fluorometer | The corresponding catalog #:<br>• Q33216<br>• Q33238 |
| | [Optional] E-Gel Power Snap Electrophoresis Device | Catalog # G8100 |
| V&P Scientific, Inc. | [Optional] MagPin M | SKU # VP 407AM-N1[1,3] |

[1] Equipment that you have tested and demonstrates equivalent performance is acceptable.

[2] The strip magnet supports strip tubes, the rack supports larger tubes, and the stand supports plates.

[3] Element recommends the MagPin M for plate-based cleanup procedures.

CHAPTER 2

### Experiment Planning

### Input Requirements

Start with 0.1–1 ng purified genomic DNA (gDNA) in 2 µl. Use Buffer EB to dilute the input gDNA.

The workflow captures and amplifies only full-length 16S ribosomal DNA (rDNA) molecules, so depletion is not necessary. However, if you typically use the Zymo HostZERO Microbial DNA Kit or equivalent for human microbiome studies, you can continue use.

#### Extraction Method

Extraction methods that perform well for V3–V4 and other short 16S regions might not yield the required long-read fragment. For degraded samples, use a Qiagen extraction kit or other gentle extraction method. Replace any elution buffer with Buffer EB.

**NOTE**
When preparing partially or highly degraded samples, the number of reported synthetic long reads might be low.

For samples of uncertain quality that require harsher preparations, such as gram-positive bacteria or soil samples, use a gel to check the purified samples for degradation. The expected result is a significant mass of ≥ 2 kb fragments, which supports amplification of full-length 16S molecules.

#### Sample Quality

The input gDNA quality affects how many 16S molecules you can successfully capture, sequence, and assemble into synthetic long reads. Inconsistent quality among samples or low purity can introduce process variations that compromise quality.

Process variations include the following examples:

- Different extraction methods or kits for DNA isolation
- Sample extractions from different physical sites, such as soil versus water
- Samples from different host organisms, such as human gut versus rumen
- Sample storage conditions

#### Quantification Method

Element recommends a Qubit fluorometer method for quantifying the input gDNA and accurately determining concentration. Quantification methods based on other DNA-binding fluorescent dye might be suitable alternatives.

NanoDrop is not recommended. Salt concentration and the presence of free nucleotides, RNA, and other contaminants that absorb at similar wavelengths to double-stranded DNA (dsDNA) can impact the reported concentration.

### Enrichment Optimization

Total DNA concentration, bacterial DNA concentration, DNA integrity, and the presence of PCR inhibitors in microbiome samples can vary significantly. Accordingly, enrichment conditions can vary across samples. To optimize the enrichment reactions, Element recommends validating the PCR conditions and performing a quality control (QC) check of the results.

#### PCR Conditions

Before starting enrichment, validate the optimal number of PCR cycles and inhibitor interference for each sample using Q5 High-Fidelity DNA Polymerase and enrichment primers 27F (forward) and 1429R (reverse). For sequences, see *User-Supplied Consumables* on page 1.

Too few cycles cause low-abundance samples to drop out. Too many cycles cause overcycling, which generates nonspecific amplification products that interfere with sequencing. Never exceed 30 cycles. If you suspect that PCR inhibitors are interfering with amplification, test 1:20 and 1:100 dilutions of the original extraction.

#### Quality Control

Size the enrichment PCR products with a Bioanalyzer 2100 Instrument or equivalent to perform a QC check of the enrichment reaction. The expected result is one peak at the correct size of 1400–2600 bp.

Additionally, you can use qPCR and SYBR Green I Nucleic Acid Gel Stain as an alternative method of determining the optimum number of PCR cycles. For each test enrichment reaction, add 0.1 µl SYBR Green I Nucleic Acid Gel Stain and reduce water by the same amount. Immediately before amplification plateaus, stop the PCR reaction.

### Index Sequences

The index primers provided in the LoopSeq Core Construction Kit include the following Index 1 (i7) sequences. Each sequence is 8 bp.

| Index Primer | Index 1 Sequence |
|---|---|
| Index Primer P1 | ATTACTCG |
| Index Primer P2 | TCCGGAGA |
| Index Primer P3 | CGCTCATT |
| Index Primer P4 | GAGATTCC |

### Protocol Parameters

Throughout protocols, observe the following parameters to avoid cross-contamination and otherwise facilitate successful prep:

- Follow the steps in the order indicated using the specified volumes and durations.
- Proceed immediately from one step to the next unless you have reached a safe stopping point.
- Always use filtered pipette tips. When adding or transferring reagents and samples, change pipette tips between each reagent and each sample.
- Return LoopSeq reagents to -30°C to -10°C storage after use.
- Combine absolute ethanol and nuclease-free water to prepare ***fresh*** 80% ethanol for each cleanup you are performing within the day. Prepare a sufficient volume for each reaction and allow 10–20% overage.
- Appropriately prepare a library removed from storage:
  - » For a library stored at 2°C to 8°C, pulse vortex and briefly centrifuge.
  - » For a library stored at -25°C to -15°C, thaw on ice. Make sure the library is fully thawed, pulse vortex, and briefly centrifuge.
- If you are using the MagPin M, supplement the cleanup instructions in this guide with the techniques described in *Care and Use of the VP 407AM-N1 96 Pin Magnetic Bead Extractor (Technical Note 310A)*. The technical note is available at vp-sci.com/wp-content/uploads/310A-VP-407AM-N1-Magnetic-Bead-Extractor.pdf.

CHAPTER 3

### Barcode Protocol

### Enrich Samples

The enrich samples procedure captures targets of interest and dilutes samples to the appropriate concentration for amplification.

1. Gather the following consumables:
   - 96-well PCR plate
   - Deepwell plate
   - Microseal ‘B’
   - Buffer EB
   - 0.1–1 ng purified gDNA
   - [Optional] Agilent High Sensitivity DNA Kit
   - [Optional] Amber microcentrifuge tube
   - [Optional] E-Gel CloneWell II Agarose Gels with SYBR Safe, 0.8%
   - [Optional] Nuclease-free water
   - [Optional] SYBR Green 1 Nucleic Acid Gel Stain
2. Remove Sample Enrichment Mix from -30°C to -10°C storage.
3. Fully thaw Sample Enrichment Mix at room temperature.
4. Briefly centrifuge Sample Enrichment Mix.
5. Add 8 µl Sample Enrichment Mix to each well of a new PCR plate.
6. If you are using real-time PCR to monitor amplification, do as follows.
   a. In a new amber microcentrifuge tube, dilute SYBR Green 1 Nucleic Acid Gel Stain 1:200 in nuclease-free water.
   b. Add 0.1 µl dilution to each reaction.
7. Add 2 µl 0.1–1 ng purified gDNA to each well.
   - Change tips between each sample.
   - Note each sample position (column and row).
8. Mix using either method:
   - Seal the plate and shake at 1800 rpm for 2 minutes.
   - Set a pipette to 7 µl and thoroughly pipette each sample. Seal the plate.
9. Briefly centrifuge the plate.
10. Place the plate in the thermal cycler.
11. Run the following ~101-minute program.

| Temperature | Time | Number of Cycles | Ramp Speed |
|---|---|---|---|
| Volume set to 10 µl | | | |
| Lid set to 100°C | | | |
| 95°C | 3 minutes | 1 | 2°C to 3°C/second |

| Temperature | Time | Number of Cycles | Ramp Speed |
|---|---|---|---|
| 95°C | 30 seconds | 30 | 2°C to 3°C/second |
| 52°C | 45 seconds | | |
| 72°C | 2 minutes | | |
| 4°C | Hold | Not applicable | 2°C to 3°C/second |

12. Remove the plate from the thermal cycler.
13. Add 198 µl Buffer EB to each well of a new deepwell plate.
14. Transfer 2 µl enriched sample from each well of the PCR plate to each well of the deepwell plate to prepare 100x diluted samples.
15. [Optional] Seal the original plate and store the remaining 8 µl sample at -25°C to -15°C for ≤ 2 months.
16. Seal the plate and briefly vortex to mix or shake at 1800 rpm for 2 minutes.
17. Briefly centrifuge the plate and immediately proceed to the next step.
18. [Optional] Evaluate the smear analysis percentage of the enriched sample:
    a. Combine 2 µl enriched sample and 38 µl Buffer EB in a new microcentrifuge tube.
    b. Size the diluted sample using a 2100 Bioanalyzer Instrument and Agilent High Sensitivity DNA Kit.
    c. Set the smear analysis region to 1400–2600 bp.
    d. If the region is < 80% amplified product, perform additional purification using E-Gel CloneWell II Agarose Gels with SYBR Safe, 0.8%.

### Amplify Enriched Samples

The amplify enriched samples procedure adds UMIs and a LoopSeq index so each sample includes per-molecule barcodes and a per-sample barcode.

1. Gather the following consumables:
   - 96-well PCR plate
   - Microseal 'B'
   - [Optional] Agilent High Sensitivity DNA Kit
   - [Optional] Amber microcentrifuge tube
   - [Optional] Nuclease-free water
   - [Optional] SYBR Green 1 Nucleic Acid Gel Stain
2. Remove the 16S SSC Barcoding Plate 96-Plex (barcode plate) from -30°C to -10°C storage.
3. Fully thaw the barcode plate on ice. Keep on ice.
4. Prepare the barcode plate:
   a. Check the plate for defects. Do not use a plate with a loose seal, cracks, or chips.
   b. Make sure the barcode oligos are fully thawed.
   c. Centrifuge the plate at 1500 rpm for 30 seconds.
   d. Clean the seal with an alcohol wipe.
5. Peel the seal covering the barcode plate to uncover the desired wells or pierce the desired wells.
   - Avoid splashing the liquid in the wells.
   - If piercing, change tips between each well.
   - Do not reuse wells, which are single-use.
6. Transfer 7.5 µl barcode oligo from each well of the barcode plate to each well of a new PCR plate.
   - Change tips between each well.
   - Pipette carefully to avoid well-to-well oligo transfer.
7. If you are using real-time PCR to monitor amplification, do as follows.
   a. In a new amber microcentrifuge tube, dilute SYBR Green 1 Nucleic Acid Gel Stain 1:200 in nuclease-free water.
   b. Add 0.1 µl dilution to each reaction.
8. Add 2.5 µl 100x diluted sample to each well of the PCR plate.
9. [Optional] Return the remaining 197.5 µl 100x diluted sample to -25°C to -15°C storage.
10. Seal the PCR plate and shake at 1800 rpm for 2 minutes.
11. Briefly centrifuge the PCR plate.
12. Place the PCR plate in the thermal cycler.
13. Run the following ~36-minute program.

| Temperature | Time | Number of Cycles | Ramp Speed |
|---|---|---|---|
| Volume set to 10 µl | | | |
| Lid set to 100°C | | | |
| 95°C | 3 minutes | 1 | 2°C to 3°C/second |
| 95°C | 30 seconds | 10 | 2°C to 3°C/second |
| 65°C | 45 seconds | | |
| 72°C | 2 minutes | | |
| 4°C | Hold | Not applicable | 2°C to 3°C/second |

14. Remove the plate from the thermal cycler.
15. Briefly centrifuge the plate and immediately proceed.
16. [Optional] Evaluate the smear analysis percentage of the enriched sample:
    a. Combine 2 µl enriched sample and 18 µl Buffer EB in a new microcentrifuge tube.
    b. Size the diluted sample using a 2100 Bioanalyzer Instrument and Agilent High Sensitivity DNA Kit.
    c. Set the smear analysis region to 1400–2600 bp.

    —The region must contain > 65% amplified product.—

### Clean Up Samples

The cleanup samples procedure removes < 300 bp fragments, remaining adapter strands, adapter dimers, and other PCR byproducts to purify the samples.

1. Gather the following consumables:
    - 96-well PCR plate or PCR tubes
    - Deepwell plate
    - Microseal 'B'
    - Buffer EB
    - Freshly prepared 80% ethanol
    - Sample purification beads
2. Add 40 µl Buffer EB to each reaction.
3. Thoroughly vortex sample purification beads to resuspend. Make sure beads are not aggregated at the bottom of the bottle.
4. Add 30 µl beads (0.6x) to each reaction.
    - Aspirate and dispense beads slowly.
    - Fully dispense beads from the pipette tip.
5. Seal the plate and shake at 1800 rpm for 2 minutes.
6. Incubate beads and sample at room temperature for 3 minutes.
7. Place the plate on the magnet and wait until the beads settle and the supernatant clears (~5 minutes). ***Keep on the magnet.***
8. Unseal the plate.
9. Remove and discard the entire volume of supernatant (~80 µl).
    - Do not disturb the bead pellets.
    - Pipette carefully to avoid aspirating beads.
10. Wash the content of each well:
    a. Without resuspending the beads, add 150 µl 80% ethanol to each reaction and incubate for 30 seconds.
    b. Remove and discard ethanol.
    c. Without resuspending the beads, add another 150 µl 80% ethanol to each reaction and incubate for 30 seconds.
    d. Remove and discard ethanol.
    e. Using a 10 µl or 20 µl pipette, remove residual ethanol.
11. Remove the plate from the magnet.
12. Add 52 µl Buffer EB to each reaction.

    —To prevent overdrying, the cleanup procedures do not air-dry the beads.—
13. Seal the plate and shake at 1800 rpm for 2 minutes.
14. Incubate the reactions at room temperature for 3 minutes.
15. Place the plate on the magnet and wait until the beads settle and the supernatant clears (~2 minutes).
16. Unseal the plate.
17. Transfer 50 µl supernatant to a new PCR plate or tubes.

**SAFE STOPPING POINT**
Immediately proceed or store the samples: seal the plate and store short-term at 2°C to 8°C for ≤ 24 hours. For long-term storage, store at -25°C to -15°C for ≤ 2 weeks.

### Calibrate Barcodes

The calibrate barcodes procedure serially dilutes each sample to a complexity of 7500 UMIs per microliter. A complexity calculator guides dilution.

1. Gather the following consumables:
   - Deepwell plates (4)
   - PCR plate
   - Microseal 'B'
   - Buffer EB
   - Qubit dsDNA HS Assay Kit
2. Add 8 µl Buffer EB to each well of a new PCR plate.
3. Transfer 2 µl enriched sample from each well or tube to each well to prepare 5x diluted samples.
4. Seal the plate and shake at 2200 rpm for 2 minutes.
5. Quantify ≥ 2 µl of each diluted sample using a Qubit dsDNA HS Assay Kit and Qubit fluorometer. Run at least duplicate reactions to verify the concentration.
6. Download the *LoopSeq Complexity Calculator* as an Excel file.
7. Open the calculator and select the **96-Plex 16S** worksheet tab.
8. Populate the calculator:
   a. In cell B1, enter the starting dilution factor or leave the default of 40.
   b. In cell B2, enter the starting sample input in µl or leave the default of 5.
   c. In column D, enter the average concentration of each sample determined in step 5.

   —The calculator autopopulates columns G, H, and J.—
9. In each well of a new deepwell plate, combine the following components to prepare dilution 1:
   - The starting sample input recorded in cell B2.
   - The Buffer EB volume indicated in column J.
10. Seal the plate and shake at 1200 rpm for 1 minute.
11. Prepare serial dilutions 2–4 per the volumes indicated in the dilution columns of the calculator. Use a new deepwell plate for each dilution.
12. Immediately proceed with dilution 4.

### Amplify and Pool Samples

The amplify and pool samples procedure amplifies the barcoded molecules and combines all samples in one tube.

1. Gather the following consumables:
   - 1.5 ml microcentrifuge tubes (2)
   - 96-well PCR plate or PCR tubes (one tube per sample)
   - Microseal 'B'
   - Buffer EB
   - [Optional] Amber microcentrifuge tube
   - [Optional] Nuclease-free water
   - [Optional] SYBR Green 1 Nucleic Acid Gel Stain
2. Remove Amplification Mix B from -30°C to -10°C storage.
3. Fully thaw Amplification Mix B at room temperature.
4. Briefly centrifuge Amplification Mix B.
5. Add 18 µl Amplification Mix B to each well of a new PCR plate.
6. If you are using real-time PCR to monitor amplification, do as follows.
   a. In a new amber microcentrifuge tube, dilute SYBR Green 1 Nucleic Acid Gel Stain 1:200 in nuclease-free water.
   b. Add 0.2 µl dilution to each reaction.
7. Add 2 µl dilution 4 to each well.
8. Seal the plate and gently vortex.
9. Briefly centrifuge the plate.
10. Place the plate in the thermal cycler.
11. Run the following ~75-minute program.

| Temperature | Time | Number of Cycles | Ramp Speed |
|---|---|---|---|
| Volume set to 20 µl | | | |
| Lid set to 100°C | | | |
| 95°C | 3 minutes | 1 | 2°C to 3°C/second |
| 95°C | 30 seconds | 22 | 2°C to 3°C/second |
| 60°C | 45 seconds | | |
| 72°C | 2 minutes | | |
| 4°C | Hold | Not applicable | 2°C to 3°C/second |

12. Remove the plate from the thermal cycler.
13. Pool the reactions in a new microcentrifuge tube.
    - If you are processing < 5 reactions, combine the volume of each reaction that achieves a total volume of 50 µl. If necessary, use Buffer EB to reach 50 µl.
    - If you are processing ≥ 5 reactions, combine 10 µl of each reaction.

    —The minimum cleanup volume is 50 µl.—

14. If you combined ≥ 50 reactions, transfer 500 µl sample pool to a new microcentrifuge tube.
15. [Optional] Seal the plate or cap the tube and store the remaining individual samples and sample pool at -25°C to -15°C for ≤ 2 weeks.

### Clean Up Sample Pool

The cleanup sample pool procedure removes < 300 bp fragments to purify the combined samples.

1. Gather the following consumables:
   » Microcentrifuge tube
   » Buffer EB
   » Freshly prepared 80% ethanol
   » Sample purification beads
   » [Optional] Agilent High Sensitivity DNA Kit
2. Thoroughly vortex sample purification beads to resuspend. Make sure beads are not aggregated at the bottom of the bottle.
3. Add 0.6x volume of beads to the reaction.
   » Aspirate and dispense beads slowly.
   » Fully dispense beads from the pipette tip.

   —For example, add 300 µl beads to a 500 µl reaction.—
4. Set a pipette to 70% of the reaction volume and pipette tube content to fully mix beads and sample.
5. Cap the tube and incubate beads and sample at room temperature for 5 minutes.
6. Place the tube on the magnet and wait until the beads settle and the supernatant clears (~5 minutes). ***Keep on the magnet.***
7. Uncap the tube.
8. Remove and discard the entire volume of supernatant.
   » Do not disturb the bead pellet.
   » Pipette carefully to avoid aspirating beads.
9. Wash the content of the tube:
   a. Without resuspending the beads, add ≥ 500 µl 80% ethanol and incubate for 30 seconds.
   b. Remove and discard ethanol.
   c. Without resuspending the beads, add ≥ 500 µl 80% ethanol and incubate for 30 seconds.
   d. Remove and discard ethanol.
10. Cap the tube and briefly centrifuge.
11. Return the tube to the magnet.
12. Using a 10 µl or 20 µl pipette, remove residual ethanol.
13. Remove the tube from the magnet.
14. Add 42 µl Buffer EB to the reaction.
15. Cap the tube and vortex for 2 minutes to resuspend.
16. Incubate the reaction at room temperature for 3 minutes.
17. Place the tube on the magnet and wait until the beads settle and the supernatant clears (~5 minutes).
18. Uncap the tube.
19. Transfer 40 µl supernatant to a new microcentrifuge tube.

20. In a new microcentrifuge tube, combine the following components.

| Component | Volume (µl) |
|---|---|
| Enriched sample | 2 |
| Buffer EB | 18 |
| Total | 20 |

21. Set a pipette to 14 µl and pipette the diluted sample 10 times to mix.
22. Quantify ≥ 2 µl of each diluted sample using a Qubit dsDNA HS Assay Kit and Qubit fluorometer. Run at least duplicate reactions to verify the concentration.
    - For < 3 ng/µl samples, repeat *Amplify and Pool Samples* on page 23 with replicate amplification wells.
    - For 3–24 ng/µl samples, proceed to *Distribute Barcodes* on page 28.
    - For > 24 ng/µl samples, normalize to 15 ng/µl using Buffer EB and proceed to *Distribute Barcodes* on page 28.
23. [Optional] Evaluate the success of the amplification procedure:
    a. Size the diluted sample using a 2100 Bioanalyzer Instrument and Agilent High Sensitivity DNA Kit.
    b. Set the smear analysis region to 1400–2600 bp.

    —If amplification is successful, the trace displays a single peak in the region.—

**SAFE STOPPING POINT**
Switch to the LoopSeq Core Construction Kit and immediately proceed to *Library Prep Protocol* on page 27. Alternatively, cap the tube and store at 2°C to 8°C for ≤ 72 hours.

CHAPTER 4

### Library Prep Protocol

### Distribute Barcodes

The distribute barcodes procedure fragments samples and randomly distributes the UMIs across molecule lengths.

1. Retrieve a new PCR plate or PCR tube.
2. Remove the following reagents from -30°C to -10°C storage:
   » Distribution Enzyme
   » Distribution Mix
3. Prepare the reagents:
   a. Thaw the reagents on ice.
   b. Vortex the tubes to mix reagents.
   c. Centrifuge the tubes. Keep on ice.
4. In the new PCR plate or tube, combine the following components.

| Component | Volume (µl) |
|---|---|
| Sample pool | 30 |
| Distribution Mix | 10 |
| Total | 40 |

5. Set a pipette to 28 µl and pipette the diluted sample 10 times to mix.
6. [Optional] Return the remaining 10 µl sample pool to 2°C to 8°C storage.
7. Add 4 µl Distribution Enzyme to the reaction.
8. Mix using either method:
   » Seal the plate or cap the tube and thoroughly vortex.
   » Set a pipette to 31 µl and thoroughly pipette plate or tube content. Seal the plate or cap the tube.
9. Briefly centrifuge the plate or tube.
10. Place the plate or tube in the thermal cycler.
11. Run the following ~20-minute program.

| Step | Temperature | Time |
|---|---|---|
| Volume set to 44 µl | | |
| Lid set to 100°C | | |
| 1 | 20°C | 15 minutes |
| 2 | 75°C | 5 minutes |
| 3 | 4°C | Hold |

12. Remove the plate or tube from the thermal cycler.
13. Briefly centrifuge the plate or tube and immediately proceed.

### Activate and Neutralize Barcodes

The activate and neutralize barcodes procedure activates the distributed barcodes and neutralizes the barcodes that did not distribute.

1. Remove the following reagents from -30°C to -10°C storage:
   » Activation Enzyme
   » Activation Mix
   » Neutralization Enzyme
2. Prepare the reagents:
   a. Thaw the reagents on ice.
   b. Vortex the tubes to mix reagents.
   c. Centrifuge the tubes. Keep on ice.
3. Add 53.5 µl Activation Mix to the reaction.
4. Add 2.5 µl Activation Enzyme to the reaction.
5. Mix using either method:
   » Seal the plate or cap the tube and thoroughly vortex.
   » Set a pipette to 70 µl and thoroughly pipette plate or tube content. Seal the plate or cap the tube.
6. Briefly centrifuge the plate or tube.
7. Place the plate or tube in the thermal cycler.
8. Run the following ~2-hour program.

| Step | Temperature | Time |
|---|---|---|
| Volume set to 100 µl | | |
| Heated lid turned off | | |
| 1 | 20°C | 2 hours |
| 2 | 4°C | Hold |

9. Remove the plate or tube from the thermal cycler.
10. Add 6 µl Neutralization Enzyme to the plate or tube.
11. Mix using either method:
    » Seal the plate or cap the tube and thoroughly vortex.
    » Set a pipette to 70 µl and thoroughly pipette plate or tube content. Seal the plate or cap the tube.
12. Briefly centrifuge the plate or tube.
13. Place the plate or tube in the thermal cycler.
14. Run the following ~15-minute program.

| Step | Temperature | Time |
|---|---|---|
| Volume set to 106 µl or 100 µl | | |
| Lid set to 100°C | | |

| Step | Temperature | Time |
|---|---|---|
| 1 | 37°C | 15 minutes |
| 2 | 4°C | Hold |

15. Remove the plate or tube from the thermal cycler.
16. Start the following thermal cycler program to prepare for *Fragment Samples and Repair Ends* on page 32.

| Step | Temperature | Time |
|---|---|---|
| Volume set to 50 µl | | |
| Lid set to 100°C | | |
| 1 | 4°C | Hold |
| 2 | 32°C | 5 minutes |
| 3 | 65°C | 30 minutes |
| 4 | 4°C | Hold |

17. Briefly centrifuge the plate or tube and immediately proceed to the next step.

### Clean Up Activated Samples

The cleanup activated samples procedure removes inactive barcodes and < 200 bp fragments to purify the sample pool.

1. Gather the following consumables:
   - PCR tube
   - Buffer EB
   - Freshly prepared 80% ethanol
   - Sample purification beads
2. Transfer the reaction to a new PCR tube.
3. Thoroughly vortex sample purification beads to resuspend. Make sure beads are not aggregated at the bottom of the bottle.
4. Add 84.8 µl beads (0.8x) to the sample pool.
   - Aspirate and dispense beads slowly.
   - Fully dispense beads from the pipette tip.
5. Set a pipette to 126 µl and pipette tube content to fully mix beads and sample.
6. Cap the tube and incubate beads and sample at room temperature for 5 minutes.
7. Place the tube on the magnet and wait until the beads settle and the supernatant clears (~3 minutes). ***Keep on the magnet.***
8. Uncap the tube.
9. Remove and discard the entire volume of supernatant (~180 µl).
   - Do not disturb the bead pellet.
   - Pipette carefully to avoid aspirating beads.
10. Wash the content of the tube:
    a. Without resuspending the beads, add 200 µl 80% ethanol to the tube and incubate for 30 seconds.
    b. Remove and discard ethanol.
    c. Without resuspending the beads, add another 200 µl 80% ethanol to the tube and incubate for 30 seconds.
    d. Remove and discard ethanol.
11. Cap the tube and briefly centrifuge.
12. Return the tube to the magnet.
13. Using a 10 µl or 20 µl pipette, remove residual ethanol.
14. Remove the tube from the magnet.
15. Add 37 µl Buffer EB to the reaction.
16. Cap the tube and vortex for 2 minutes to resuspend.
17. Incubate the reaction at room temperature for 3 minutes.
18. Place the tube on the magnet and wait until the beads settle and the supernatant clears (~3 minutes).
19. Uncap the tube.
20. Transfer 35 µl supernatant to a new PCR tube.

**SAFE STOPPING POINT**
Immediately proceed or cap the tube and store at 2°C to 8°C for ≤ 72 hours.

### Fragment Samples and Repair Ends

The fragment sample and repair ends procedure digests the samples into fragments and blunts the damaged ends. An A-tailing reaction adds an adenine (A) base to the blunt ends to prepare for adapter ligation.

1. Remove the following reagents from -30°C to -10°C storage:
   - » Fragmentation Enzyme
   - » Fragmentation Mix
2. Prepare the reagents:
   a. Thaw the reagents on ice.
   b. Vortex the tubes to mix reagents.
   c. Centrifuge the tubes. Keep on ice.
3. Place the sample pool on ice. ***Keep on ice.***

   **CAUTION**
   Assembling the reaction on ice and keeping it cold prevents over-fragmentation.
4. Add 5 µl Fragmentation Mix to the pool.
5. Add 10 µl Fragmentation Enzyme to the pool.
6. Without vortexing, mix using either method:
   - » Set a pipette to 35 µl, pipette tube content, and cap.
   - » Cap the tube and flick.
7. Briefly centrifuge the tube.
8. Immediately place the tube in the prechilled thermal cycler and resume the ~35-minute program.
9. Remove the tube from the thermal cycler.
10. Briefly centrifuge the tube and immediately proceed to the next step.

### Ligate Adapters

The ligate adapters procedure adds primer sites for attaching surface primers and generates the library.

1. Remove the following reagents from -30°C to -10°C storage:
   » Ligation Enzyme
   » Ligation Mix
2. Prepare the reagents:
   a. Thaw the reagents on ice.
   b. Vortex the tubes to mix reagents.
   c. Centrifuge the tubes. Keep on ice.
3. Add 40 µl Ligation Mix to the reaction.
4. Add 10 µl Ligation Enzyme to the reaction.
5. Mix using either method:
   » Set a pipette to 70 µl, pipette tube content, and cap.
   » Cap the tube and thoroughly vortex.
6. Briefly centrifuge the tube.
7. Place the tube in the thermal cycler.
8. Run the following ~15-minute program.

| Step | Temperature | Time |
|---|---|---|
| Volume set to 100 µl | | |
| Heated lid turned off | | |
| 1 | 20°C | 15 minutes |
| 2 | 4°C | Hold |

9. Remove the tube from the thermal cycler.
10. Briefly centrifuge the tube and immediately proceed to the next step.

### Clean Up Library

The cleanup library procedure removes < 300 bp fragments, remaining adapter strands, adapter dimer, and other PCR byproducts to purify the library.

1. Gather the following consumables:
   - Microcentrifuge tube
   - PCR tube
   - Buffer EB
   - Freshly prepared 80% ethanol
   - Sample purification beads
2. Transfer the 100 µl reaction to a new microcentrifuge tube.
3. Thoroughly vortex sample purification beads to resuspend. Make sure beads are not aggregated at the bottom of the bottle.
4. Add 60 µl beads (0.6x) to the reaction.
   - Aspirate and dispense beads slowly.
   - Fully dispense beads from the pipette tip.
5. Set a pipette to 112 µl and pipette tube content to fully mix beads and library.
6. Cap the tube and incubate beads and library at room temperature for 5 minutes.
7. Place the tube on the magnet and wait until the beads settle and the supernatant clears (~3 minutes). ***Keep on the magnet.***
8. Uncap the tube.
9. Remove and discard the entire volume of supernatant (~160 µl).
   - Do not disturb the bead pellet.
   - Pipette carefully to avoid aspirating beads.
10. Wash the content of the tube:
    a. Without resuspending the beads, add 200 µl 80% ethanol to the tube and incubate for 30 seconds.
    b. Remove and discard ethanol.
    c. Without resuspending the beads, add another 200 µl 80% ethanol to the tube and incubate for 30 seconds.
    d. Remove and discard ethanol.
11. Cap the tube and briefly centrifuge.
12. Return the tube to the magnet.
13. Using a 10 µl or 20 µl pipette, remove residual ethanol.
14. Remove the tube from the magnet.
15. Add 22 µl Buffer EB to each reaction.
16. Cap the tube and vortex for 2 minutes to resuspend.
17. Incubate the reaction at room temperature for 3 minutes.
18. Place the tube on the magnet and wait until the beads settle and the supernatant clears (~3 minutes).
19. Uncap the tube.
20. Transfer 20 µl supernatant to a new PCR tube.

**SAFE STOPPING POINT**
Immediately proceed or cap the tube and store at 2°C to 8°C for ≤ 72 hours.

### Amplify Library

The amplify library procedure attaches an Index 1 sequence to the 3′ end of each library in the pool.

1. If you are using real-time PCR to monitor amplification, gather the following consumables:
   » Amber microcentrifuge tube
   » Nuclease-free water
   » SYBR Green 1 Nucleic Acid Gel Stain
2. Remove the following reagents from -30°C to -10°C storage:
   » Index Master Mix
   » Index Primer P1, P2, P3, or P4 (choose one)
3. Prepare the reagents:
   a. Thaw the reagents on ice.
   b. Vortex the tubes to mix reagents.
   c. Centrifuge the tubes. Keep on ice.
4. Add 25 µl Index Master Mix to the reaction.
5. If you are using real-time PCR to monitor amplification, do as follows.
   a. In a new amber microcentrifuge tube, dilute SYBR Green 1 Nucleic Acid Gel Stain 1:200 in nuclease-free water.
   b. Add 0.5 µl dilution to each reaction.
6. Add 5 µl Index Primer P1, P2, P3, or P4 to the reaction.
7. Mix using either method:
   » Set a pipette to 35 µl, pipette tube content, and cap.
   » Cap the tube and thoroughly vortex.
8. Briefly centrifuge the tube.
9. Place the tube in the thermal cycler.
10. Run the following ~35-minute program.

| Temperature | Time | Number of Cycles |
|---|---|---|
| Volume set to 50 µl | | |
| Lid set to 100°C | | |
| 95°C | 3 minutes | 1 |
| 95°C | 30 seconds | 10 |
| 65°C | 45 seconds | |
| 72°C | 30 seconds | |
| 4°C | Hold | Not applicable |

11. Remove the tube from the thermal cycler.
12. Briefly centrifuge the tube and immediately proceed to the next step.

### Clean Up Indexed Library

The cleanup indexed library procedure removes remaining oligos and < 300 bp fragments to purify the indexed library.

1. Gather the following consumables:
   » 1.5 ml microcentrifuge tubes (2)
   » Buffer EB
   » Freshly prepared 80% ethanol
   » Sample purification beads
2. Transfer the 50 µl reaction to a new microcentrifuge tube.
3. Thoroughly vortex sample purification beads to resuspend. Make sure beads are not aggregated at the bottom of the bottle.
4. Add 30 µl beads (0.6x) to the reaction.
   » Aspirate and dispense beads slowly.
   » Fully dispense beads from the pipette tip.
5. Set a pipette to 56 µl and pipette tube content to fully mix beads and library.
6. Cap the tube and incubate beads and library at room temperature for 5 minutes.
7. Place the tube on the magnet and wait until the beads settle and the supernatant clears (~3 minutes). ***Keep on the magnet.***
8. Uncap the tube.
9. Remove and discard the entire volume of supernatant (~80 µl).
   » Do not disturb the bead pellet.
   » Pipette carefully to avoid aspirating beads.
10. Wash the content of the tube:
    a. Without resuspending the beads, add 200 µl 80% ethanol to the tube and incubate for 30 seconds.
    b. Remove and discard ethanol.
    c. Without resuspending the beads, add another 200 µl 80% ethanol to the tube and incubate for 30 seconds.
    d. Remove and discard ethanol.
11. Cap the tube and briefly centrifuge.
12. Return the tube to the magnet.
13. Using a 10 µl or 20 µl pipette, remove residual ethanol.
14. Remove the tube from the magnet.
15. Add 22 µl Buffer EB to the reaction.
16. Cap the tube and vortex for 2 minutes to resuspend.
17. Incubate the reaction at room temperature for 3 minutes.
18. Place the tube on the magnet and wait until the beads settle and the supernatant clears (~3 minutes).
19. Uncap the tube.
20. Transfer 20 µl supernatant to a new microcentrifuge tube.

**SAFE STOPPING POINT**
If you are not immediately quantifying and sequencing, cap the tube and store short-term at 2°C to 8°C for ≤ 1 week. For long-term storage, store at -25°C to -15°C for ≤ 1 month.

### Size and Quantify Library

The size and quantify library procedure performs a QC check of the library.

1. Dilute the library 1:10 in Buffer EB and size the diluted library using a 2100 Bioanalyzer Instrument and Agilent High Sensitivity DNA Kit.

   **Figure 3:** Example trace of a correct library

   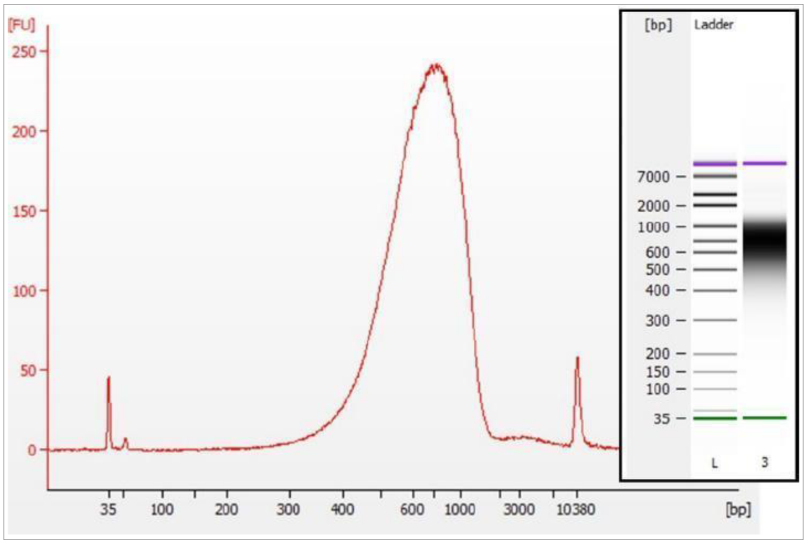

2. Quantify the library using either of the following methods to determine concentration:
   - » Dilute the library 1:10 in Buffer EB. Quantify ≥2 µl diluted library using a Qubit fluorometer with the Qubit dsDNA HS Assay Kit.
   - » Quantify the library using a qPCR system with the KAPA Library Quantification Kit - Complete kit (Universal) or equivalent. Follow manufacturer instructions.
3. Proceed to sequencing, following the Illumina instructions for your system.

### Technical Support

Visit the User Documentation page on the Element Biosciences website for additional guides and the most recent version of this guide. For technical assistance, contact Element Technical Support.

**Website:** www.elementbiosciences.com

**Telephone:** +1 866.ELEMBIO (+1 866.353.6246)

### Document History

| Document # | Date | Description of Change |
|---|---|---|
| Document # MA-00027 Rev. D | December 2023 | • Added evaluation and real-time PCR steps.<br>• Added user-supplied consumables and equipment for evaluation steps and real-time PCR.<br>• Updated the evaluation steps for sizing and quantifying the library.<br>• Updated the consumable lists for several procedures.<br>• Updated the thermal cycler volume for activating and neutralizing barcodes.<br>• Updated the bead volume for activated sample cleanup.<br>• Updated the estimated duration of the barcode protocol.<br>• Moved the fragmentation thermal cycler program to barcode activation and neutralization. |
| Document # MA-00027 Rev. C | June 2023 | • Removed redundant input gDNA concentration.<br>• Corrected the volume of remaining 100x diluted sample.<br>• Corrected the barcode plate and enrichment mix names.<br>• Updated the instructions on preparing the barcode plate and calibrating barcodes.<br>• Updated links to the complexity calculator, MagPin M technical note, and user guides.<br>• Updated pipette settings to 70% of a total volume. |
| Document # MA-00027 Rev. B | December 2022 | • Added MagPin M (V&P Scientific, Inc., SKU # VP 407AM-N1). |
| Document # MA-00027 Rev. A | November 2022 | • Initial release |

Element Biosciences
10055 Barnes Canyon Road, Suite 100
San Diego, CA 92121
elementbiosciences.com


**ELEMENT BIOSCIENCES**
**For Research Use Only. Not for use in diagnostic procedures.**
