## Supplementary material for "Complex bacterial diversity of Guaymas Basin hydrothermal sediments revealed by synthetic long-read sequencing (LoopSeq)": DNA Extraction Protocol

DNA extraction from Guaymas Basin sediment samples was adapted from a previously published extraction method that combines physical, chemical, and enzymatic steps and was optimized for sediment samples (Zhou, Bruns, & Tiedje 1996). We scaled it down from 3 g sediment to 0.5 g sediment and adjusted all components accordingly.

1.35 ml of DNA extraction buffer (see below) was added to 0.5 g frozen sediment and stirred with a sterile spatula.

**DNA extraction buffer	for 300 ml total**

100 mM Tris-HCl, pH 8.0 (adjusted to pH 8.0 with 1 M HCl)	30 ml of 1M stock

100 mM EDTA, pH 8.0	60 ml of 0.5M stock

100 mM sodium phosphate buffer, pH 8.0	60 ml of 0.5M Stock

1.5 M NaCl	90 ml of 5M Stock

1% CTAB	3 g to 300 ml

Physical extraction:

The slurry was frozen in at -20°C and thawed thoroughly in a water bath at 65°C. Freezing and thawing was repeated three times.

Biological extraction:

After the slurry cooled down to 35°C, 5 μl of Proteinase K (20 mg ml-1) was added and samples incubated for 30 min at 37°C under mild horizontal shaking. This step can be extended ad lib, e.g., to an overnight incubation with enzyme. (For difficult samples, e.g., containing many gram-positive bacteria it is advised to include an incubation with Lysozyme before the Proteinase step; we note that this additional step was not used with our Guaymas Basin samples).

Chemical extraction:

0.3 ml of 10% sodium-dodecylsulfate (SDS) was added and incubated (2 hours, 65°C) with gentle end over end inversion every 25 min. The slurry was centrifuged (3220 *g* for 10 min at RT) and the supernatant collected.

To increase yield, an additional extraction step can be used, but was not performed here. In this step (described for 3 g sediment), the pellet was again treated with 2.7 ml DNA extraction buffer and 0.6 ml 10% SDS, incubated for 10 min at RT and centrifuged (3220 *g* for 10 min at RT). Supernatants were combined (~11 ml), and samples can be split as needed to the final organic extraction; an equal volume of chloroform/isoamylalcohol (24:1) was added and the mixture was shaken gently, but thoroughly. After centrifugation (3220 *g* for 10 min at RT) the aqueous phase was collected and 0.6 volumes of isopropanol (abs.) were added to precipitate the DNA (overnight, 4°C).

Pellet washing

The pellet was washed with cold ethanol (80% v/v), centrifuged (20,000 *g* for 10 min at 4°C), dried (15 -30 min at RT) and gently resuspended (1 hour at 4°C, no pipetting) in a suitable volume (50 µl – 250 µl) of 0.5× TE or PCR-water. If CTAB crystals form in the precipitate, the suspension was warmed up in a water bath (1 min at 70°C) to dissolve precipitated CTAB, centrifuged (20,000 *g* for 25 min at RT) and the supernatant discarded.

Pellet clean-up:

The DNA suspension was immediately purified using the Wizard DNA clean-up system, which is based on the adsorption and desorption of DNA to a matrix. The DNA was eluted twice with 25 μl 0.5× TE (65 °C) and the yield checked by gel electrophoresis and subsequent staining in an ethidium bromide bath (30 min at RT). Gel electrophoresis was always carried out with agarose gels (1% agarose in 1× TAE) in Biorad electrophoresis chambers containing 1x TAE at an electrical current of 8-10 V cm-1. Pockets were loaded with 2 μl of DNA extract (or low DNA mass ladder) mixed with 1 μl of 6× loading buffer. Photometric measurements of DNA concentrations were performed in triplicates with Nanodrop or Qubit fluorometer.

**Reference**: Zhou J, Bruns MA, Tiedje JM. 1996. DNA recovery from soils of diverse composition. Applied and Environmental Microbiology. 62:316-22.

**Images displaying the color gradient of chromophoric biomass in Guaymas Basin sediments after phenol-chloroform centrifugation and before the removal of the organic phase. Note the removal of chromophoric biomass with depth and hydrothermal influence.**

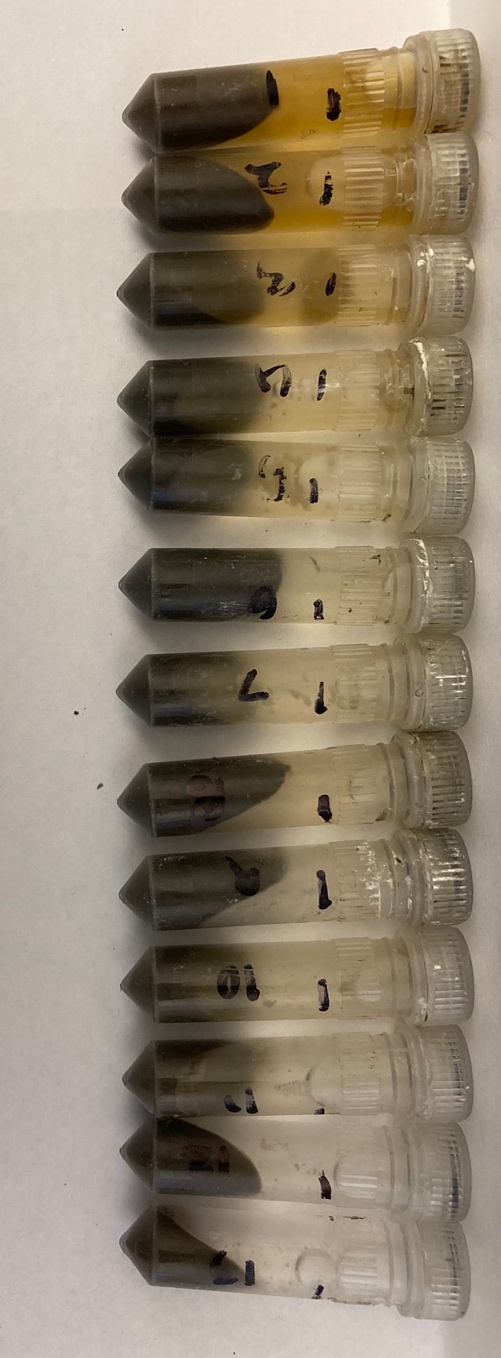

Bare sediment site samples (4872-01). Samples arranged by depth left to right (leftmost sample = top layer of sediments, rightmost sample = bottom layer of sediments).

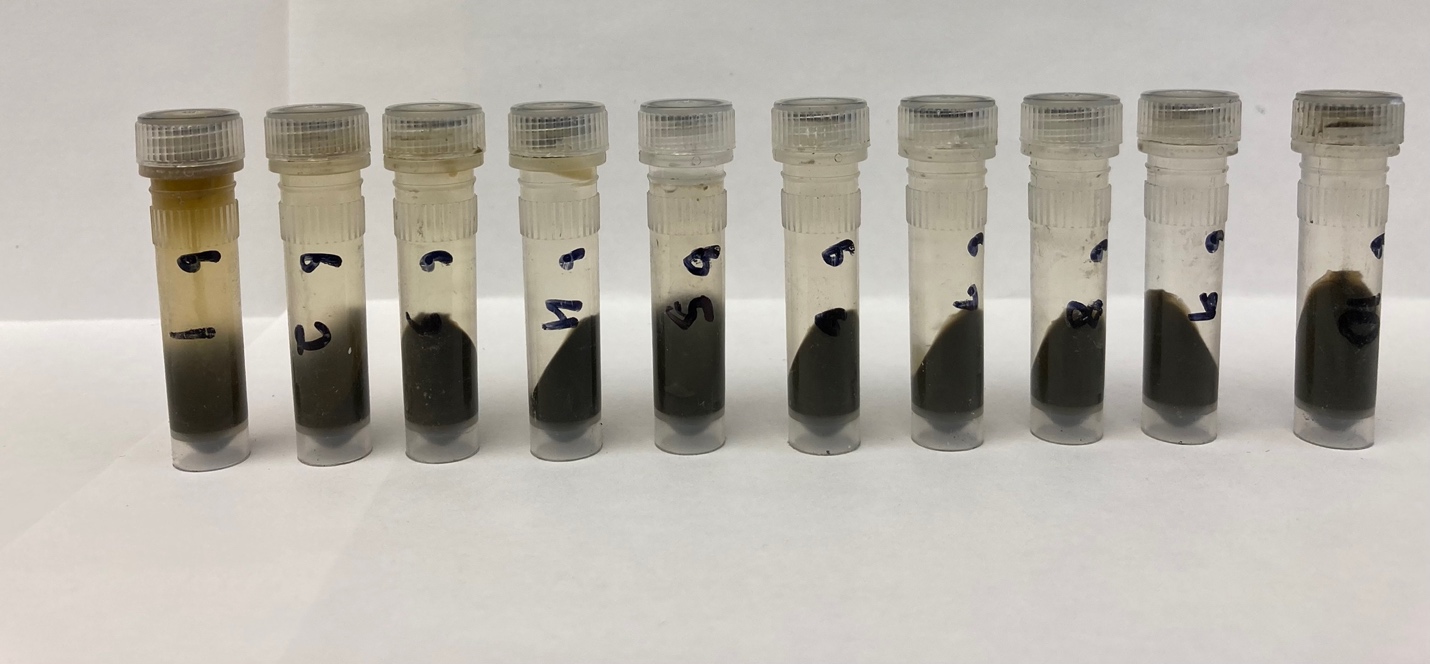

White mat sediment site samples (4872-06). Samples arranged by depth left to right (leftmost sample = top layer of sediments, rightmost sample = bottom layer of sediments).

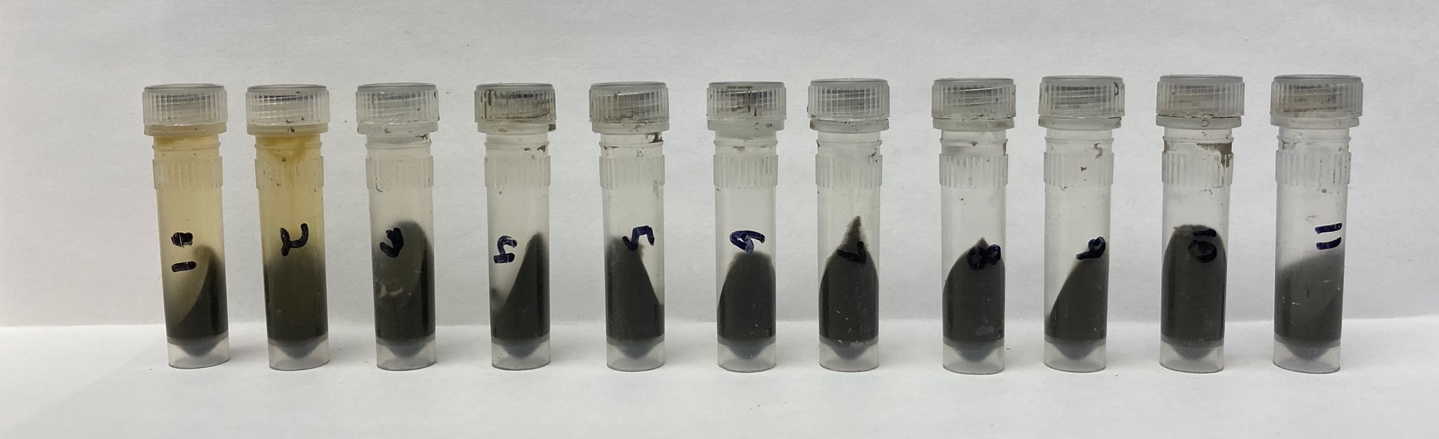

Orange mat sediment site samples (4872-14). Samples arranged by depth left to right (leftmost sample = top layer of sediments, rightmost sample = bottom layer of sediments).
