## Supplementary Data File 2 for "Complex bacterial diversity of Guaymas Basin hydrothermal sediments revealed by synthetic long-read sequencing (LoopSeq)"

>GuaymasBasin_Dive4872_L-ASV_8212

ATTGAACGCTGGCGGCATGCTTAATACATGCAAGTCGAACGGTAACAGGTCCTTCGGGATGCTGACGAGTGGCGGACGGGTGAGTAATGCATAGGAATCTGCCCAGTAGTAGGGGACAACCTGAGGAAACTCAGGCTAATACCGCATAAGTCCTACGGGAGAAAGGGGGCCTCTTCTTGAAAGCTCTCGCTATTGGATGAGCCTATGTCGGATTAGCTTGTTGGTGGGGTAATGGCCTACCAAGGCTGCGATCCGTAGCTGGTCTGAGAGGACGATCAGCCACACTGGGACTGAGACACGGCCCAGACTCCTACGGGAGGCAGCAGTGGGGAATATTGGACAATGGGCGAAAGCCTGATCCAGCAATGCCGCGTGTGTGAAGAAGGCTTGCGGGTTGTAAAGCACTTTCAGTTGGGAAGAAAAGTATTAGGTTAATACTCTAATATTTTGACGTTACCAACAGAAGAAGCACCGGCTAACTCTGTGCCAGCAGCCGCGGTAATACAGAGGGTGCGAGCGTTAATCGGAATTACTGGGCGTAAAGCGTGCGTAGGCGGCTAAGTCAGTCGGATGTGAAAGCCCAAGGCTTAACCTTGGAACTGCATTCGATACTGCTTGACTAGAGTACAGTAGAGGGAAGCGGAATTCTTAGTGTAGCGGTGAAATGCGTAGATATTAGGAAGAACACCAGTTGCGAAGGCGGCTTCCTGGACTGATACTGACGCTGAGGTACGAAAGCGTGGGGAGCAAACAGGATTAGATACCCTGGTAGTCCACGCCCTAAACGATGAGAACTAGATGTTGGGGGAAATTGATCCCTTAGTATCGCAGCTAACGCCATAAGTTCTCCGCCTGGGGAGTACGACCGCAAGGTTAAAACTCAAATGAATTGACGGGGGCCCGCACAAGCGGTGGAGCATGTGGTTTAATTCGATGCAACGCGAAGAACCTTACCTGGCCTTGACATCCTCGGGACCTCGCAGAGATGTGAGGGTGCCTTCGGGAACCGAGAGACAGGTGCTGCATGGCTGTCGTCAGCTCGTGTCGTGAGATGTTGGGTTAAGTCCCGCAACGAGCGCAACCCTTGTCCCTAGTTGCCAGCGATTCGGTCGGGAACTCTAGAGAGACTGCCGGTGATAAACCGGAGGAAGGTGGGGATGACGTCAAGTCATCATGGCCCTTACGGCCAGGGCTACACACGTGCTACAATGGGTAGTACAGAGGGTCGCAAACCCGCGAGGGGGAGCTAATCTCACAAAACTACTCGTAGTCCGGATTGGAGTCTGCAACTCGACTCCATGAAGTCGGAATCGCTAGTAATCGCGAATCAGCATGTCGCGGTGAATACGTTCCCGGGCCTTGTACACACCGCCCGTCACACCATGGGAGTGGGCTGTACCAGAAGTAGGTAGTCTAACCGCAAGGGGGACGCTTACCACGGTATGGTTCATGACTGGGGTG

>GuaymasBasin_Dive4872_L-ASV_8231

ATTGAACGCTGGCGGCATGCTTAATACATGCAAGTCGAACGGTAACAGGTCCTTCGGGATGCTGACGAGTGGCGGACGGGTGAGTAATGCATAGGAATCTGCCCAGTAGTAGGGGACAACCTGAGGAAACTCAGGCTAATACCGCATAAGTCCTACGGGAGAAAGGGGGCCTCTTCTTGAAAGCTCTCGCTATTGGATGAGCCTATGTCGGATTAGCTTGTTGGTGGGGTAATGGCCTACCAAGGCTGCGATCCGTAGCTGGTCTGAGAGGACGATCAGCCACACTGGGACTGAGACACGGCCCAGACTCCTACGGGAGGCAGCAGTGGGGAATATTGGACAATGGGCGAAAGCCTGATCCAGCAATGCCGCGTGTGTGAAGAAGGCTTGCGGGTTGTAAAGCACTTTCAGTTGGGAAGAAAAGTATTAGGTTAATACTCTAATATTTTGACGTTACCAACAGAAGAAGCACCGGCTAACTCTGTGCCAGCAGCCGCGGTAATACAGAGGGTGCGAGCGTTAATCGGAATTACTGGGCGTAAAGCGTGCGTAGGCGGCTAAGTCAGTCGGATGTGAAAGCCCAAGGCTTAACCTTGGAACTGCATTCGATACTGCTTGACTAGAGTACAGTAGAGGGAAGCGGAATTCTTAGTGTAGCGGTGAAATGCGTAGATATTAGGAAGAACACCAGTTGCGAAGGCGGCTTCCTGGACTGATACTGACGCTGAGGTACGAAAGCGTGGGGAGCAAACAGGATTAGATACCCTGGTAGTCCACGCCCTAAACGATGAGAACTAGATGTTGGGGGAAATTGATCCCTTAGTATCGCAGCTAACGCCATAAGTTCTCCGCCTGGGGAGTACGACCGCAAGGTTAAAACTCAAATGAATTGACGGGGGCCCGCACAAGCGGTGGAGCATGTGGTTTAATTCGATGCAACGCGAAGAACCTTACCTGGCCTTGACATCCTCGGGACCTCGCAGAGATGTGAGGGTGCCTTCGGGAACCGAGAGACAGGTGCTGCATGGCTGTCGTCAGCTCGTGTCGTGAGATGTTGGGTTAAGTCCCGCAACGAGCGCAACCCTTGTCCCTAGTTGCCAGCGATTCGGTCGGGAACTCTAGAGAGACTGCCGGTGACAAACCGGAGGAAGGTGGGGATGACGTCAAGTCATCATGGCCCTTACGGCCAGGGCTACACACGTGCTACAATGGGTAGTACAGAGGGTCACAAACCCGCGAGGGGGAGCTAATCTCACAAAACTACTCGTAGTCCGGATTGGAGTCTGCAACTCGACTCCATGAAGTCGGAATCGCTAGTAATCGCGAATCAGCATGTCGCGGTGAATACGTTCCCGGGCCTTGTACACACCGCCCGTCACACCATGGGAGTGGGCTGTACCAGAAGTAGGTAGTCTAACCGCAAGGGGGACGCTTACCACGGTATGGTTCATGACTGGGGTG

>GuaymasBasin_Dive4872_L-ASV_8564

ATTGAACGCTGGCGGCATGCTTAATACATGCAAGTCGAACGGTAACAGGTCCTTCGGGATGCTGACGAGTGGCGGACGGGTGAGTAATGCATAGGAATCTGCCCAGTAGTAGGGGACAACCTGAGGAAACTCAGGCTAATACCGCATAAGTCCTACGGGAGAAAGGGGGCCTCTTCTTGAAAGCTCTCGCTATTGGATGAGCCTATGTCGGATTAGCTTGTTGGTGGGGTAATGGCCTACCAAGGCTGCGATCCGTAGCTGGTCTGAGAGGACGATCAGCCACACTGGGACTGAGACACGGACCAGACTCCTACGGGAGGCAGCAGTGAGGAATATTGGACAATGGGCGAAAGCCTGATCCAGCAATGCCGCGTGTGTGAAGAAGGCTTGCGGGTTGTAAAGCACTTTCAGTTGGGAAGAAAAGTATTAGGTTAATACTCTAATATTTTGACGTTACCAACAGAAGAAGCACCGGCTAACTCTGTGCCAGCAGCCGCGGTAATACAGAGGGTGCGAGCGTTAATCGGAATTACTGGGCGTAAAGCGTGCGTAGGCGGCTAAGTCAGTCGGATGTGAAAGCCCAAGGCTTAACCTTGGAACTGCATTCGATACTGCTTGACTAGAGTACAGTAGAGGGAAGCGGAATTCTTAGTGTAGCGGTGAAATGCGTAGATATTAGGAAGAACACCAGTTGCGAAGGCGGCTTCCTGGACTGATACTGACGCTGAGGTACGAAAGCGTGGGGAGCAAACAGGATTAGATACCCTGGTAGTCCACGCCCTAAACGATGAGAACTAGATGTTGGGGGAAATTGATCCCTTAGTATCGCAGCTAACGCCATAAGTTCTCCGCCTGGGGAGTACGACCGCAAGGTTAAAACTCAAATGAATTGACGGGGGCCCGCACAAGCGGTGGAGCATGTGGTTTAATTCGATGCAACGCGAAGAACCTTACCTGGCCTTGACATCCTCGGGACCTCGCAGAGATGTGAGGGTGCCTTCGGGAACCGAGAGACAGGTGCTGCATGGCTGTCGTCAGCTCGTGTCGTGAGATGTTGGGTTAAGTCCCGCAACGAGCGCAACCCTTGTCCCTAGTTGCCAGCGATTCGGTCGGGAACTCTAGAGAGACTGCCGGTGACAAACCGGAGGAAGGTGGGGATGACGTCAAGTCATCATGGCCCTTACGGCCAGGGCTACACACGTGCTACAATGGGTAGTACAGAGGGTCGCAAACCCGCGAGGGGGAGCTAATCTCACAAAACTACTCGTAGTCCGGATTGGAGTCTGCAACTCGACTCCATGAAGTCGGAATCGCTAGTAATCGCGAATCAGCATGTCGCGGTGAATACGTTCCCGGGCCTTGTACACACCGCCCGTCACACCATGGGAGTGGGCTGTACCAGAAGTAGGTAGTCTAACCGCAAGGGGGACGCTTACCACGGTATGGTTCATGACTGGGGTG

>GuaymasBasin_Dive4872_L-ASV_8488

ATTGAACGCTGGCGGCATGCTTAATACATGCAAGTCGAACGGTAACAGGTCCTTCGGGATGCTGACGAGTGGCGGACGGGTGAGTAATGCATAGGAATCTGCCCAGTAGTAGGGGACAACCTGAGGAAACTCAGGCTAATACCGCATAAGTCCTACGGGAGAAAGGGGGCCTCTTCTTGAAAGCTCTCGCTATTGGATGAGCCTATGTCGGATTAGCTTGTTGGTGGGGTAATGGCCTACCAAGGCTGCGATCCGTAGCTGGTCTGAGAGGATGATCAGCCACACTGGGACTGAGACACGGCCCAGACTCCTACGGGAGGCAGCAGTGGGGAATATTGGACAATGGGCGAAAGCCTGATCCAGCAATGCCGCGTGTGTGAAGAAGGCTTGCGGGTTGTAAAGCACTTTCAGTTGGGAAGAAAAGTATTAGGTTAATACTCTAATATTTTGACGTTACCAACAGAAGAAGCACCGGCTAACTCTGTGCCAGCAGCCGCGGTAATACGGAGGGTGCGAGCGTTAATCGGAATTACTGGGCGTAAAGCGTGCGTAGGCGGCTAAGTCAGTCGGATGTGAAAGCCCAAGGCTTAACCTTGGAACTGCATTCGATACTGCTTGACTAGAGTACAGTAGAGGGAAGCGGAATTCTTAGTGTAGCGGTGAAATGCGTAGATATTAGGAAGAACACCAGTTGCGAAGGCGGCTTCCTGGACTGATACTGACGCTGAGGTACGAAAGCGTGGGGAGCAAACAGGATTAGATACCCTGGTAGTCCACGCCCTAAACGATGAGAACTAGATGTTGGGGGAAATTGATCCCTTAGTATCGCAGCTAACGCCATAAGTTCTCCGCCTGGGGAGTACGACCGCAAGGTTAAAACTCAAATGAATTGACGGGGGCCCGCACAAGCGGTGGAGCATGTGGTTTAATTCGATGCAACGCGAAGAACCTTACCTGGCCTTGACATCCTCGGGACCTCGCAGAGATGTGAGGGTGCCTTCGGGAACCGAGAGACAGGTGCTGCATGGCTGTCGTCAGCTCGTGTCGTGAGATGTTGGGTTAAGTCCCGCAACGAGCGCAACCCTTGTCCCTAGTTGCCAGCGATTCGGTCGGGAACTCTAGAGAGACTGCCGGTGACAAACCGGAGGAAGGTGGGGATGACGTCAAGTCATCATGGCCCTTACGGCCAGGGCTACACACGTGCTACAATGGGTAGTACAGAGGGTCGCAAACCCGCGAGGGGGAGCTAATCTCACAAAACTACTCGTAGTCCGGATTGGAGTCTGCAACTCGACTCCATGAAGTCGGAATCGCTAGTAATCGCGAATCAGCATGTCGCGGTGAATACGTTCCCGGGCCTTGTACACACCGCCCGTCACACCATGGGAGTGGGCTGTACCAGAAGTAGGTAGTCTAACCGCAAGGGGGACGCTTACCACGGTATGGTTCATGACTGGGGTG

>GuaymasBasin_Dive4872_L-ASV_8842

ATTGAACGCTGGCGGCATGCTTAATACATGCAAGTCGAACGGTAACAGGTCCTTCGGGATGCTGACGAGTGGCGGACGGGTGAGTAATGCATAGGAATCTGCCCAGTAGTAGGGGACAACCTGAGGAAACTCAGGCTAATACCGCATAAGTCCTACGGGAGAAAGGGGGCCTCTTCTTGAAAGCTCTCGCTATTGGATGAGCCTATGTCGGATTAGCTTGTTGGTGGGGTAATGGCCTACCAAGGCTGCGATCCGTAGCTGGTCTGAGAGGACGATCAGCCACACTGGGACTGAGACACGGCCCAGACTCCTACGGGAGGCAGCAGTGGGGAATATTGGACAATGGGCGAAAGCCTGATCCAGCAATGCCGCGTGTGTGAAGAAGGCTTGCGGGTTGTAAAGCACTTTCAGTTGGGAAGAAAAGTATTAGGTTAATACTCTAATATTTTGACGTTACCAACAGAAGAAGCACCGGCTAACTCTGTGCCAGCAGCCGCGGTAATACGGAGGGTGCGAGCGTTAATCGGAATTACTGGGCGTAAAGCGTGCGTAGGCGGCTAAGTCAGTCGGATGTGAAAGCCCAAGGCTTAACCTTGGAACTGCATTCGATACTGCTTGACTAGAGTACAGTAGAGGGAAGCGGAATTCTTAGTGTAGCGGTGAAATGCGTAGATATTAGGAAGAACACCAGTTGCGAAGGCGGCTTCCTGGACTGATACTGACGCTGAGGTACGAAAGCGTGGGGAGCAAACAGGATTAGATACCCTGGTAGTCCACGCCCTAAACGATGAGAACTAGATGTTGGGGGAAATTGATCCCTTAGTATCGCAGCTAACGCCATAAGTTCTCCGCCTGGGGAGTACGACCGCAAGGTTAAAACTCAAATGAATTGACGGGGGCCCGCACAAGCGGTGGAGCATGTGGTTTAATTCGATGCAACGCGAAGAACCTTACCTGGCCTTGACATCCTCGGGACCTCGCAGAGATGTGAGGGTGCCTTCGGGAACCGAGAGACAGGTGCTGCATGGCTGTCGTCAGCTCGTGTCGTGAGATGTTGGGTTAAGTCCCGCAACGAGCGCAACCCTTGTCCCTAGTTGCCAGCGATTCGGTCGGGAACTCTAGAGAGACTGCCGGTGACAAACCGGAGGAAGGTGGGGATGACGTCAAGTCATCATGGCCCTTACGGCCAGGGCTACACACGTGCTACAATGGGTAGTACAGAGGGTCGCAAACCCGCGAGGGGGAGCTAATCTCACAAAACTACTCGTAGTCCGGATTGGAGTCTGCAACTCGACTCCATGAAGTCGGAATCGCTAGTAATCGCGAATCAGCATGTCGCGGTGAATACGTTCCCGGGCCTTGTACACACCGCCCGTCACACCATGGGAGTGGGCTGTACCAGAAGTAGGTAGTCTAACCGCAAGGGGGACGCTTACCACGGTATGGTTCATGACTGGGGTG

>GuaymasBasin_Dive4872_L-ASV_8848

ATTGAACGCTGGCGGCATGCTTAATACATGCAAGTCGAACGGTAACAGGTCCTTCGGGATGCTGACGAGTGGCGGACGGGTGAGTAATGCATAGGAATCTGCCCAGTAGTAGGGGACAACCTGAGGAAACTCAGGCTAATACCGCATAAGTCCTACGGGAGAAAGGGGGCCTCTTCTTGAAAGCTCTCGCTATTGGATGAGCCTATGTCGGATTAGCTTGTTGGTGGGGTAATGGCCTACCAAGGCTGCGATCCGTAGCTGGTCTGAGAGGACGATCAGCCACACTGGGACTGAGACACGGCCCAGACTCCTACGGGAGGCAGCAGTGGGGAATATTGGACAATGGGCGAAAGCCTGATCCAGCAATGCCGCGTGTGTGAAGAAGGCTTGCGGGTTGTAAAGCACTTTCAGTTGGGAAGAAAAGTATTAGGTTAATACTCTAATATTTTGACGTTACCAACAGAAGAAGCACCGGCTAACTCTGTGCCAGCAGCCGCGGTAATACAGAGGGTGCGAGCGTTAATCGGAATTACTGGGCGTAAAGCGTGCGTAGGCGGCTAAGTCAGTCGGATGTGAAAGCCCAAGGCTTAACCTTGGAACTGCATTCGATACTGCTTGACTAGAGTACAGTAGAGGGAAGCGGAATTCTTAGTGTAGCGGTGAAATGCGTAGATATTAGGAAGAACACCAGTTGCGAAGGCGGCTTCCTGGACTGATACTGACGCTGAGGTACGAAAGCGTGGGGAGCAAACAGGATTAGATACCCTGGTAGTCCACGCCGTAAACGATGAGAACTAGATGTTGGGGGAAATTGATCCCTTAGTATCGCAGCTAACGCCATAAGTTCTCCGCCTGGGGAGTACGACCGCAAGGTTAAAACTCAAATGAATTGACGGGGGCCCGCACAAGCGGTGGAGCATGTGGTTTAATTCGATGCAACGCGAAGAACCTTACCTGGCCTTGACATCCTCGGGACCTCGCAGAGATGTGAGGGTGCCTTCGGGAACCGAGAGACAGGTGCTGCATGGCTGTCGTCAGCTCGTGTCGTGAGATGTTGGGTTAAGTCCCGCAACGAGCGCAACCCTTGTCCCTAGTTGCCAGCGATTCGGTCGGGAACTCTAGAGAGACTGCCGGTGACAAACCGGAGGAAGGTGGGGATGACGTCAAGTCATCATGGCCCTTACGGCCAGGGCTACACACGTGCTACAATGGGTAGTACAGAGGGTCGCAAACCCGCGAGGGGGAGCTAATCTCACAAAACTACTCGTAGTCCGGATTGGAGTCTGCAACTCGACTCCATGAAGTCGGAATCGCTAGTAATCGCGAATCAGCATGTCGCGGTGAATACGTTCCCGGGCCTTGTACACACCGCCCGTCACACCATGGGAGTGGGCTGTACCAGAAGTAGGTAGTCTAACCGCAAGGGGGACGCTTACCACGGTATGGTTCATGACTGGGGTG

>GuaymasBasin_Dive4872_L-ASV_8881

ATTGAACGCTGGCGGCATGCTTAATACATGCAAGTCGAACGGTAACAGGTCCTTCGGGATGCTGACGAGTGGCGGACGGGTGAGTAATGCATAGGAATCTGCCCAGTAGTAGGGGACAACCTGAGGAAACTCAGGCTAATACCGCATAAGTCCTACGGGAGAAAGGGGGCCTCTTCTTGAAAGCTCTCGCTATTGGATGAGCCTATGTCGGATTAGCTTGTTGGTGGGGTAATGGCCTACCAAGGCTGCGATCCGTAGCTGGTCTGAGAGGATGATCAGCCACACTGGGACTGAGACACGGCCCAGACTCCTACGGGAGGCAGCAGTAGGGAATATTGGACAATGGGCGAAAGCCTGATCCAGCAATGCCGCGTGTGTGAAGAAGGCTTGCGGGTTGTAAAGCACTTTCAGTTGGGAAGAAAAGTATTAGGTTAATACTCTAATATTTTGACGTTACCAACAGAAGAAGCACCGGCTAACTCTGTGCCAGCAGCCGCGGTAATACAGAGGGTGCGAGCGTTAATCGGAATTACTGGGCGTAAAGCGTGCGTAGGCGGCTAAGTCAGTCGGATGTGAAAGCCCAAGGCTTAACCTTGGAACTGCATTCGATACTGCTTGACTAGAGTACAGTAGAGGGAAGCGGAATTCTTAGTGTAGCGGTGAAATGCGTAGATATTAGGAAGAACACCAGTTGCGAAGGCGGCTTCCTGGACTGATACTGACGCTGAGGTACGAAAGCGTGGGGAGCAAACAGGATTAGATACCCTGGTAGTCCACGCCCTAAACGATGAGAACTAGATGTTGGGGGAAATTGATCCCTTAGTATCGCAGCTAACGCCATAAGTTCTCCGCCTGGGGAGTACGACCGCAAGGTTAAAACTCAAATGAATTGACGGGGGCCCGCACAAGCGGTGGAGCATGTGGTTTAATTCGATGCAACGCGAAGAACCTTACCTGGCCTTGACATCCTCGGGACCTCGCAGAGATGTGAGGGTGCCTTCGGGAACCGAGAGACAGGTGCTGCATGGCTGTCGTCAGCTCGTGTCGTGAGATGTTGGGTTAAGTCCCGCAACGAGCGCAACCCTTGTCCCTAGTTGCCAGCGATTCGGTCGGGAACTCTAGAGAGACTGCCGGTGACAAACCGGAGGAAGGTGGGGATGACGTCAAGTCATCATGGCCCTTACGGCCAGGGCTACACACGTGCTACAATGGGTAGTACAGAGGGTCGCAAACCCGCGAGGGGGAGCTAATCTCACAAAACTACTCGTAGTCCGGATTGGAGTCTGCAACTCGACTCCATGAAGTCGGAATCGCTAGTAATCGCGAATCAGCATGTCGCGGTGAATACGTTCCCGGGCCTTGTACACACCGCCCGTCACACCATGGGAGTGGGCTGTACCAGAAGTAGGTAGTCTAACCGCAAGGGGGACGCTTACCACGGTATGGTTCATGACTGGGGTG

>GuaymasBasin_Dive4872_L-ASV_2125

ATTGAACGCTGGCGGCATGCTTAATACATGCAAGTCGAACGGTAACAGGTCCTTCGGGATGCTGACGAGTGGCGGACGGGTGAGTAATGCATAGGAATCTGCCCAGTAGTAGGGGACAACCTGAGGAAACTCAGGCTAATACCGCATAAGTCCTACGGGAGAAAGGGGGCCTCTTCTTGAAAGCTCTCGCTATTGGATGAGCCTATGTCGGATTAGCTTGTTGGTGGGGTAATGGCCTACCAAGGCTGCGATCCGTAGCTGGTCTGAGAGGACGATCAGCCACACTGGGACTGAGACACGGCCCAGACTCCTACGGGAGGCAGCAGTAGGGAATATTGGACAATGGGCGAAAGCCTGATCCAGCAATGCCGCGTGTGTGAAGAAGGCTTGCGGGTTGTAAAGCACTTTCAGTTGGGAAGAAAAGTATTAGGTTAATACTCTAATATTTTGACGTTACCAACAGAAGAAGCACCGGCTAACTCTGTGCCAGCAGCCGCGGTAATACAGAGGGTGCGAGCGTTAATCGGAATTACTGGGCGTAAAGCGTGCGTAGGCGGCTAAGTCAGTCGGATGTGAAAGCCCAAGGCTTAACCTTGGAACTGCATTCGATACTGCTTGACTAGAGTACAGTAGAGGGAAGCGGAATTCTTAGTGTAGCGGTGAAATGCGTAGATATTAGGAAGAACACCAGTTGCGAAGGCGGCTTCCTGGACTGATACTGACGCTGAGGTACGAAAGCGTGGGGAGCAAACAGGATTAGATACCCTGGTAGTCCACGCCCTAAACGATGAGAACTAGATGTTGGGGGAAATTGATCCCTTAGTATCGCAGCTAACGCCATAAGTTCTCCGCCTGGGGAGTACGACCGCAAGGTTAAAACTCAAATGAATTGACGGGGGCCCGCACAAGCGGTGGAGCATGTGGTTTAATTCGATGCAACGCGAAGAACCTTACCTGGCCTTGACATCCTCGGGACCTCGCAGAGATGTGAGGGTGCCTTCGGGAACCGAGAGACAGGTGCTGCATGGCTGTCGTCAGCTCGTGTCGTGAGATGTTGGGTTAAGTCCCGCAACGAGCGCAACCCTTGTCCTTAGTTGCCAGCGATTCGGTCGGGAACTCTAGAGAGACTGCCGGTGACAAACCGGAGGAAGGTGGGGATGACGTCAAGTCATCATGGCCCTTACGGCCAGGGCTACACACGTGCTACAATGGGTAGTACAGAGGGTCGCAAACCCGCGAGGGGGAGCTAATCTCACAAAACTACTCGTAGTCCGGATTGGAGTCTGCAACTCGACTCCATGAAGTCGGAATCGCTAGTAATCGCGAATCAGCATGTCGCGGTGAATACGTTCCCGGGCCTTGTACACACCGCCCGTCACACCATGGGAGTGGGCTGTACCAGAAGTAGGTAGTCTAACCGCAAGGGGGACGCTTACCACGGTATGGTTCATGACTGGGGTG

>GuaymasBasin_Dive4872_L-ASV_121

ATTGAACGCTGGCGGCATGCTTAATACATGCAAGTCGAACGGTAACAGGTCCTTCGGGATGCTGACGAGTGGCGGACGGGTGAGTAATGCATAGGAATCTGCCCAGTAGTAGGGGACAACCTGAGGAAACTCAGGCTAATACCGCATAAGTCCTACGGGAGAAAGGGGGCCTCTTCTTGAAAGCTCTCGCTATTGGATGAGCCTATGTCGGATTAGCTTGTTGGTGGGGTAATGGCCTACCAAGGCTGCGATCCGTAGCTGGTCTGAGAGGACGATCAGCCACACTGGGACTGAGACACGGCCCAGACTCCTACGGGAGGCAGCAGTAGGGAATATTGGACAATGGGCGAAAGCCTGATCCAGCAATGCCGCGTGTGTGAAGAAGGCTTGCGGGTTGTAAAGCACTTTCAGTTGGGAAGAAAAGTATTAGGTTAATACTCTAATATTTTGACGTTACCAACAGAAGAAGCACCGGCTAACTCTGTGCCAGCAGCCGCGGTAATACAGAGGGTGCGAGCGTTAATCGGAATTACTGGGCGTAAAGCGTGCGTAGGCGGCTAAGTCAGTCGGATGTGAAAGCCCAAGGCTTAACCTTGGAACTGCATTCGATACTGCTTGACTAGAGTACAGTAGAGGGAAGCGGAATTCTTAGTGTAGCGGTGAAATGCGTAGATATTAGGAAGAACACCAGTTGCGAAGGCGGCTTCCTGGACTGATACTGACGCTGAGGTACGAAAGCGTGGGGAGCAAACAGGATTAGATACCCTGGTAGTCCACGCCCTAAACGATGAGAACTAGATGTTGGGGGAAATTGATCCCTTAGTATCGCAGCTAACGCCATAAGTTCTCCGCCTGGGGAGTACGACCGCAAGGTTAAAACTCAAATGAATTGACGGGGGCCCGCACAAGCGGTGGAGCATGTGGTTTAATTCGATGCAACGCGAAGAACCTTACCTGGCCTTGACATCCTCGGGACCTCGCAGAGATGTGAGGGTGCCTTCGGGAACCGAGAGACAGGTGCTGCATGGCTGTCGTCAGCTCGTGTCGTGAGATGTTGGGTTAAGTCCCGCAACGAGCGCAACCCTTGTCCCTAGTTGCCAGCGATTCGGTCGGGAACTCTAGAGAGACTGCCGGTGACAAACCGGAGGAAGGTGGGGATGACGTCAAGTCATCATGGCCCTTACGGCCAGGGCTACACACGTGCTACAATGGGTAGTACAGAGGGTCGCAAACCCGCGAGGGGGAGCTAATCTCACAAAACTACTCGTAGTCCGGATTGGAGTCTGCAACTCGACTCCATGAAGTCGGAATCGCTAGTAATCGCGAATCAGCATGTCGCGGTGAATACGTTCCCGGGCCTTGTACACACCGCCCGTCACACCATGGGAGTGGGCTGTACCAGAAGTAGGTAGTCTAACCGCAAGGGGGACGCTTACCACGGTATGGTTCATGACTGGGGTG

>GuaymasBasin_Dive4872_L-ASV_8243

ATTGAACGCTGGCGGCATGCTTAATACATGCAAGTCGAACGGTAACAGGTCCTTCGGGATGCTGACGAGTGGCGGACGGGTGAGTAATGCATAGGAATCTGCCCAGTAGTAGGGGACAACCTGAGGAAACTCAGGCTAATACCGCATAAGTCCTACGGGAGAAAGGGGGCCTCTTCTTGAAAGCTCTCGCTATTGGATGAGCCTATGTCGGATTAGCTTGTTGGTGGGGTAATGGCCTACCAAGGCTGCGATCCGTAGCTGGTCTGAGAGGACGATCAGCAACACTGGGACTGAGACACGGCCCAGACTCCTACGGGAGGCAGCAGTAGGGAATATTGGACAATGGGCGAAAGCCTGATCCAGCAATGCCGCGTGTGTGAAGAAGGCTTGCGGGTTGTAAAGCACTTTCAGTTGGGAAGAAAAGTATTAGGTTAATACTCTAATATTTTGACGTTACCAACAGAAGAAGCACCGGCTAACTCTGTGCCAGCAGCCGCGGTAATACAGAGGGTGCGAGCGTTAATCGGAATTACTGGGCGTAAAGCGTGCGTAGGCGGCTAAGTCAGTCGGATGTGAAAGCCCAAGGCTTAACCTTGGAACTGCATTCGATACTGCTTGACTAGAGTACAGTAGAGGGAAGCGGAATTCTTAGTGTAGCGGTGAAATGCGTAGATATTAGGAAGAACACCAGTTGCGAAGGCGGCTTCCTGGACTGATACTGACGCTGAGGTACGAAAGCGTGGGGAGCAAACAGGATTAGATACCCTGGTAGTCCACGCCCTAAACGATGAGAACTAGATGTTGGGGGAAATTGATCCCTTAGTATCGCAGCTAACGCCATAAGTTCTCCGCCTGGGGAGTACGACCGCAAGGTTAAAACTCAAATGAATTGACGGGGGCCCGCACAAGCGGTGGAGCATGTGGTTTAATTCGATGCAACGCGAAGAACCTTACCTGGCCTTGACATCCTCGGGACCTCGCAGAGATGTGAGGGTGCCTTCGGGAACCGAGAGACAGGTGCTGCATGGCTGTCGTCAGCTCGTGTCGTGAGATGTTGGGTTAAGTCCCGCAACGAGCGCAACCCTTGTCCCTAGTTGCCAGCGATTCGGTCGGGAACTCTAGAGAGACTGCCGGTGACAAACCGGAGGAAGGTGGGGATGACGTCAAGTCATCATGGCCCTTACGGCCAGGGCTACACACGTGCTACAATGGGTAGTACAGAGGGTCGCAAACCCGCGAGGGGGAGCTAATCTCACAAAACTACTCGTAGTCCGGATTGGAGTCTGCAACTCGACTCCATGAAGTCGGAATCGTTAGTAATCGCGAATCAGCATGTCGCGGTGAATACGTTCCCGGGCCTTGTACACACCGCCCGTCACACCATGGGAGTGGGCTGTACCAGAAGTAGGTAGTCTAACCGCAAGGGGGACGCTTACCACGGTATGGTTCATGACTGGGGTG

>GuaymasBasin_Dive4872_L-ASV_8882

ATTGAACGCTGGCGGCATGCTTAATACATGCAAGTCGAACGGTAACAGGTCCTTCGGGATGCTGACGAGTGGCGGACGGGTGAGTAATGCATAGGAATCTGCCCAGTAGTAGGGGACAACCTGAGGAAACTCAGGCTAATACCGCATAAGTCCTACGGGAGAAAGGGGGCCTCTTCTTGAAAGCTCTCGCTATTGGATGAGCCTATGTCGGATTAGCTTGTTGGTGGGGTAATGGCCTACCAAGGCTGCGATCCGTAGCTGGTCTGAGAGGACGATCAGCCACACTGGGACTGAGACACGGCCCAGACTCCTACGGGAGGCAGCAGTAGGGAATATTGGACAATGGGCGAAAGCCTGATCCAGCAATGCCGCGTGTGTGAAGAAGGCTTGCGGGTTGTAAAGCACTTTCAGTTGGGAAGAAAAGTATTAGGTTAATACTCTAATATTTTGACGTTACCAACAGAAGAAGCACCGGCTAACTCTGTGCCAGCAGCCGCGGTAATATAGAGGGTGCGAGCGTTAATCGGAATTACTGGGCGTAAAGCGTGCGTAGGCGGCTAAGTCAGTCGGATGTGAAAGCCCAAGGCTTAACCTTGGAACTGCATTCGATACTGCTTGACTAGAGTACAGTAGAGGGAAGCGGAATTCTTAGTGTAGCGGTGAAATGCGTAGATATTAGGAAGAACACCAGTTGCGAAGGCGGCTTCCTGGACTGATACTGACGCTGAGGTACGAAAGCGTGGGGAGCAAACAGGATTAGATACCCTGGTAGTCCACGCCCTAAACGATGAGAACTAGATGTTGGGGGAAATTGATCCCTTAGTATCGCAGCTAACGCCATAAGTTCTCCGCCTGGGGAGTACGACCGCAAGGTTAAAACTCAAATGAATTGACGGGGGCCCGCACAAGCGGTGGAGCATGTGGTTTAATTCGATGCAACGCGAAGAACCTTACCTGGCCTTGACATCCTCGGGACCTCGCAGAGATGTGAGGGTGCCTTCGGGAACCGAGAGACAGGTGCTGCATGGCTGTCGTCAGCTCGTGTCGTGAGATGTTGGGTTAAGTCCCGCAACGAGCGCAACCCTTGTCCCTAGTTGCCAGCGATTCGGTCGGGAACTCTAGAGAGACTGCCGGTGACAAACCGGAGGAAGGTGGGGATGACGTCAAGTCATCATGGCCCTTACGGCCAGGGCTACACACGTGCTACAATGGGTAGTACAGAGGGTCGCAAACCCGCGAGGGGGAGCTAATCTCACAAAACTACTCGTAGTCCGGATTGGAGTCTGCAACTCGACTCCATGAAGTCGGAATCGCTAGTAATCGCGAATCAGCATGTCGCGGTGAATACGTTCCCGGGCCTTGTACACACCGCCCGTCACACCATGGGAGTGGGCTGTACCAGAAGTAGGTAGTCTAACCGCAAGGGGGACGCTTACCACGGTATGGTTCATGACTGGGGTG

>GuaymasBasin_Dive4872_L-ASV_2086

ATTGAACGCTGGCGGCATGCTTAATACATGCAAGTCGAACGGTAACAGGTCCTTCGGGATGCTGACGAGTGGCGGACGGGTGAGTAATGCATAGGAATCTGCCCAGTAGTAGGGGACAACCTGAGGAAACTCAGGCTAATACCGCATAAGTCCTACGGGAGAAAGGGGGCCTCTTCTTGAAAGCTCTCGCTATTGGATGAGCCTATGTCGGATTAGCTTGTTGGTGGGGTAATGGCCTACCAAGGCTGCGATCCGTAGCTGGTCTGAGAGGACGATCAGCCACACTGGGACTGAGACACGGCCCAGACTCCTACGGGAGGCAGCAGTAGGGAATATTGGACAATGGGCGAAAGCCTGATCCAGCAATGCCGCGTGTGTGAAGAAGGCTTGCGGGTTGTAAAGCACTTTCAGTTGGGAAGAAAAGTATTAGGTTAATACTCTAATATTTTGACGTTACCAACAGAAGAAGCACCGGCTAACTCTGTGCCAGCAGCCGCGGTAATACAGAGGGTGCGAGCGTTAATCGGAATTACTGGGCGTAAAGCGTGCGTAGGCGGCTAAGTCAGTCGGATGTGAAAGCCCAAGGCTTAACCTTGGAACTGCATTCGATACTGCTTGACTAGAGTACAGTAGAGGGAAGCGGAATTCTTAGTGTAGCGGTGAAATGCGTAGATATTAGGAAGAACACCAGTTGCGAAGGCGGCTTCCTGGACTGATACTGACGCTGAGGTACGAAAGCGTGGGGAGCAAACAGGATTAGATACCCTGGTAGTCCACGCCCTAAACGATGAGAACTAGATGTTGGGGGAAATTGATCCCTTAGTATCGCAGCTAACGCCATAAGTTCTCCGCCTGGGGAGTACGGCCGCAAGGTTAAAACTCAAAGGAATTGACGGGGGCCCGCACAAGCGGTGGAGCATGTGGTTTAATTCGATGCAACGCGAAGAACCTTACCTGGCCTTGACATCCTCGGGACCTCGCAGAGATGTGAGGGTGCCTTCGGGAACCGAGAGACAGGTGCTGCATGGCTGTCGTCAGCTCGTGTCGTGAGATGTTGGGTTAAGTCCCGCAACGAGCGCAACCCTTGTCCCTAGTTGCCAGCGATTCGGTCGGGAACTCTAGAGAGACTGCCGGTGACAAACCGGAGGAAGGTGGGGATGACGTCAAGTCATCATGGCCCTTACGGCCAGGGCTACACACGTGCTACAATGGGTAGTACAGAGGGTCGCAAACCCGCGAGGGGGAGCTAATCTCACAAAACTACTCGTAGTCCGGATTGGAGTCTGCAACTCGACTCCATGAAGTCGGAATCGCTAGTAATCGCGAATCAGCATGTCGCGGTGAATACGTTCCCGGGCCTTGTACACACCGCCCGTCACACCATGGGAGTGGGCTGTACCAGAAGTAGGTAGTCTAACCGCAAGGGGGACGCTTACCACGGTATGGTTCATGACTGGGGTG

>GuaymasBasin_Dive4872_L-ASV_8481

ATTGAACGCTGGCGGCATGCTTAATACATGCAAGTCGAACGGTAACAGGTCCTTCGGGATGCTGACGAGTGGCGGACGGGTGAGTAATGCATAGGAATCTGCCCAGTAGTAGGGGACAACCTGAGGAAACTCAGGCTAATACCGCATAAGTCCTACGGGAGAAAGGGGGCCTCTTCTTGAAAGCTCTCGCTATTGGATGAGCCTATGTCGGATTAGCTTGTTGGTGGGGTAATGGCCTACCAAGGCTGCGATCCGTAGCTGGTCTGAGAGGACGATCAGCCACACTGGGACTGAGACACGGCCCAGACTCCTACGGGAGGCAGCAGTGGGGAATTTTGCACAATGGGCGAAAGCCTGATCCAGCAATGCCGCGTGTGTGAAGAAGGCTTGCGGGTTGTAAAGCACTTTCAGTTGGGAAGAAAAGTATTAGGTTAATACTCTAATATTTTGACGTTACCAACAGAAGAAGCACCGGCTAACTCTGTGCCAGCAGCCGCGGTAATACAGAGGGTGCGAGCGTTAATCGGAATTACTGGGCGTAAAGCGTGCGTAGGCGGCTAAGTCAGTCGGATGTGAAAGCCCAAGGCTTAACCTTGGAACTGCATTCGATACTGCTTGACTAGAGTACAGTAGAGGGAAGCGGAATTCTTAGTGTAGCGGTGAAATGCGTAGATATTAGGAAGAACACCAGTTGCGAAGGCGGCTTCCTGGACTGATACTGACGCTGAGGTACGAAAGCGTGGGGAGCAAACAGGATTAGATACCCTGGTAGTCCACGCCCTAAACGATGAGAACTAGATGTTGGGGGAAATTGATCCCTTAGTATCGCAGCTAACGCCATAAGTTCTCCGCCTGGGGAGTACGACCGCAAGGTTAAAACTCAAATGAATTGACGGGGGCCCGCACAAGCGGTGGAGCATGTGGTTTAATTCGATGCAACGCGAAGAACCTTACCTGGCCTTGACATCCTCGGGACCTCGCAGAGATGTGAGGGTGCCTTCGGGAACCGAGAGACAGGTGCTGCATGGCTGTCGTCAGCTCGTGTCGTGAGATGTTGGGTTAAGTCCCGCAACGAGCGCAACCCTTGTCCCTAGTTGCCAGCGATTCGGTCGGGAACTCTAGAGAGACTGCCGGTGACAAACCGGAGGAAGGTGGGGATGACGTCAAGTCATCATGGCCCTTACGGCCAGGGCTACACACGTGCTACAATGGGTAGTACAGAGGGTCGCAAACCCGCGAGGGGGAGCTAATCTCACAAAACTACTCGTAGTCCGGATTGGAGTCTGCAACTCGACTCCATGAAGTCGGAATCGCTAGTAATCGCGAATCAGCATGTCGCGGTGAATACGTTCCCGGGCCTTGTACACACCGCCCGTCACACCATGGGAGTGGGCTGTACCAGAAGTAGGTAGTCTAACCGCAAGGGGGACGCTTACCACGGTATGGTTCATGACTGGGGTG

>GuaymasBasin_Dive4872_L-ASV_206

ATTGAACGCTGGCGGCATGCTTAATACATGCAAGTCGAACGGTAACAGGTCCTTCGGGATGCTGACGAGTGGCGGACGGGTGAGTAATGCATAGGAATCTGCCCAGTAGTAGGGGACAACCTGAGGAAACTCAGGCTAATACCGCATAAGTCCTACGGGAGAAAGGGGGCCTCTTCTTGAAAGCTCTCGCTATTGGATGAGCCTATGTCGGATTAGCTTGTTGGTGGGGTAATGGCCTACCAAGGCTGCGATCCGTAGCTGGTCTGAGAGGACGATCAGCCACACTGGGACTGAGACACGGCCCAGACTCCTACGGGAGGCAGCAGTAGGGAATATTGGACAATGGGCGAAAGCCTGATCCAGCAATGCCGCGTGTGTGAAGAAGGCTTGCGGGTTGTAAAGCACTTTCAGTTGGGAAGAAAAGTATTAGGTTAATACTCTAATATTTTGACGTTACCAACAGAAGAAGCACCGGCTAACTCTGTGCCAGCAGCCGCGGTAATACAGAGGGTGCGAGCGTTAATCGGAATTACTGGGCGTAAAGCGTGCGTAGGCGGCTAAGTCAGTCGGATGTGAAAGCCCAAGGCTTAACCTTGGAACTGCATTCGATACTGCTTGACTAGAGTACAGTAGAGGGAAGCGGAATTCTTAGTGTAGCGGTGAAATGCGTAGATATTAGGAAGAACACCAGTTGCGAAGGCGGCTTCCTGGACTGATACTGACGCTGAGGTACGAAAGCGTGGGGAGCAAACAGGATTAGATACCCTGGTAGTCCACGCCCTAAACGATGAGAACTAGATGTTGGGGGAAATTGATCCCTTAGTATCGCAGCTAACGCCATAAGTTCTCCGCCTGGGGAGTACGACCGCAAGGTTAAAACTCAAAGGAATTGACGGGGGCCCGCACAAGCGGTGGAGCATGTGGTTTAATTCGATGCAACGCGAAGAACCTTACCTGGCCTTGACATCCTCGGGACCTCGCAGAGATGTGAGGGTGCCTTCGGGAACCGAGAGACAGGTGCTGCATGGCTGTCGTCAGCTCGTGTCGTGAGATGTTGGGTTAAGTCCCGCAACGAGCGCAACCCTTGTCCCTAGTTGCCAGCGATTCGGTCGGGAACTCTAGAGAGACTGCCGGTGACAAACCGGAGGAAGGTGGGGATGACGTCAAGTCATCATGGCCCTTACGGCCAGGGCTACACACGTGCTACAATGGGTAGTACAGAGGGTCGCAAACCCGCGAGGGGGAGCTAATCTCACAAAACTACTCGTAGTCCGGATTGGAGTCTGCAACTCGACTCCATGAAGTCGGAATCGCTAGTAATCGCGAATCAGCATGTCGCGGTGAATACGTTCCCGGGCCTTGTACACACCGCCCGTCACACCATGGGAGTGGGCTGTACCAGAAGTAGGTAGTCTAACCGCAAGGGGGACGCTTACCACGGTATGGTTCATGACTGGGGTG

>GuaymasBasin_Dive4872_L-ASV_8511

ATTGAACGCTGGCGGCATGCTTAATACATGCAAGTCGAACGGTAACAGGTCCTTCGGGATGCTGACGAGTGGCGGACGGGTGAGTAATGCATAGGAATCTGCCCAGTAGTAGGGGACAACCTGAGGAAACTCAGGCTAATACCGCATAAGTCCTACGGGAGAAAGGGGGCCTCTTCTTGAAAGCTCTCGCTATTGGATGAGCCTATGTCGGATTAGCTTGTTGGTGGGGTAATGGCCTACCAAGGCTGCGATCCGTAGCTGGTCTGAGAGGACGATCAGCCACACTGGGACTGAGACACGGCCCAGACTCCTACGGGAGGCAGCAGTAGGGAATATTGGACAATGGGCGAAAGCCTGATCCAGCAATGCCGCGTGTGTGAAGAAGGCTTGCGGGTTGTAAAGCACTTTCAGTTGGGAAGAAAAGTATTAGGTTAATACTCTAATATTTTGACGTTACCAACAGAAGAAGCACCGGCTAACTCCGTGCCAGCAGCCGCGGTAATACGGAGGGTGCAAGCGTTAATCGGAATTACTGGGCGTAAAGCGTGCGTAGGCGGCTAAGTCAGTCGGATGTGAAAGCCCAAGGCTTAACCTTGGAACTGCATTCGATACTGCTTGACTAGAGTACAGTAGAGGGAAGCGGAATTCTTAGTGTAGCGGTGAAATGCGTAGATATTAGGAAGAACACCAGTTGCGAAGGCGGCTTCCTGGACTGATACTGACGCTGAGGTACGAAAGCGTGGGGAGCAAACAGGATTAGATACCCTGGTAGTCCACGCCCTAAACGATGAGAACTAGATGTTGGGGGAAATTGATCCCTTAGTATCGCAGCTAACGCCATAAGTTCTCCGCCTGGGGAGTACGACCGCAAGGTTAAAACTCAAAGGAATTGACGGGGGCCCGCACAAGCGGTGGAGCATGTGGTTTAATTCGATGCAACGCGAAGAACCTTACCTGGGCTTGACATCCTCGGGACCTCGCAGAGATGTGAGGGTGCCTTCGGGAACCGAGAGACAGGTGCTGCATGGCTGTCGTCAGCTCGTGTCGTGAGATGTTGGGTTAAGTCCCGCAACGAGCGCAACCCTTGTCCCTAGTTGCCAGCGATTCGGTCGGGAACTCTAGAGAGACTGCCGGTGACAAACCGGAGGAAGGTGGGGATGACGTCAAGTCATCATGGCCCTTACGGCCAGGGCTACACACGTGCTACAATGGGTAGTACAGAGGGTCGCAAACCCGCGAGGGGGAGCTAATCTCACAAAACTACTCGTAGTCCGGATTGGAGTCTGCAACTCGACTCCATGAAGTCGGAATCGCTAGTAATCGCGAATCAGCATGTCGCGGTGAATACGTTCCCGGGCCTTGTACACACCGCCCGTCACACCATGGGAGTGGGCTGTACCAGAAGTAGGTAGTCTAACCGCAAGGGGGACGCTTACCACGGTATGGTTCATGACTGGGGTG

>GuaymasBasin_Dive4872_L-ASV_726

ATTGAACGCTGGCGGCATGCTTAATACATGCAAGTCGAACGGTAACATGCCCTTCGGGGTGATGACGAGTGGCGGACGGGTGAGTAATGCATAGGAATCTACCCAGTAGACGGGGAACAACTTGGGGAAACTCAAGCTAATACCGCGTAAGCCTTACGGGGGATAGCGGGGGACTTTTTAGGAAGCCTCGCGATATTGGATGAGCCTATGCCGGATTAGCTAGTTGGTGGGGTAAAAGCTTACCAAGGCTACGATCCGTAGCTGGTCTGAGAGGACGATCAGCCACACTGGGACTGAGACACGGCCCAGACTCCTACGGGAGGCAGCAGTAGGGAATATTGGACAATGGGCGAAAGCTTGATCCAGCAATGCCGCGTGTGTGAAGAAGGCCTGCGGGTTGTAAAGCACTTTCAGTTGGGAAGAAAAGCTTTAGGTTAATACCCTGAAGTATTGACGTTACCAACAGAAGAAGCACCGGCTAACTCTGTGCCAGCAGCCGCGGTAATACAGAGGGTGCGAGCGTTAATCGGAATTACTGGGCGTAAAGCGTACGTAGGCGGTTGAGTCAGTCGGATGTGAAAGCCCAAGGCTTAACCTTGGAACTGCATTCGATACTACTCGGCTAGAGTACAGGAGAGGGAAGCGGAATTCCTAGTGTAGCGGTGAAATGCGTAGAGATTAGGAGGAACACCAGTGGCGAAGGCGGCTTCTTGGACTGATACTGACGCTGAGGTACGAAAGCGTGGGGAGCAAACAGGATTAGATACCCTGGTAGTCCACGCCCTAAACGATGAGAACTAGATGTTGGGGGAATTTAATCCCTTAGTATCGCAGCTAACGCGCTAAGTTCTCCGCCTGGGGAGTACGGCCGCAAGGTTAAAACTCAAAGGAATTGACGGGGGCCCGCACAAGCGGTGGAGCATGTGGTTTAATTCGATGCAACGCGAAGAACCTTACCTGGCCTTGACATCCTTGGAACCTCGCAGAGATGTGAGGGTGCCTTCGGGAACCGAGAGACAGGTGCTGCATGGCTGTCGTCAGCTCGTGTCGTGAGATGTTGGGTTAAGTCCCGCAACGAGCGCAACCCTTATCCCTAGTTGCCAGCGATTCGGTCGGGAACTCTAGAGAGACTGCCGGTGACAAACCGGAGGAAGGTGGGGACGACGTCAAGTCATCATGGCCCTTACGGCCAGGGCTACACACGTGCTACAATGGGGTAGTACAAAGGGTTGCGAACCCGCGAGGGGGTGCTAATCTCACAAAACTACTCGTAGTCCGGATTGGAGTCTGCAACTCGACTCCATGAAGTTGGAATCGCTAGTAATCGCGGATCAGCATGTCGCGGTGAATACGTTCCCGGGCCTTGTACACACCGCCCGTCACACCATGGGAGTGGGCTGTACCAGAAGTAGGTAGTCTAACCGCAAGGGGGACGCTTACCACGGTATGGTTCATGACTGGGGTG

>GuaymasBasin_Dive4872_L-ASV_10139

ATTGAACGCTGGCGGCATGCTTAATACATGCAAGTCGAACGGTAACATGCCCTTCGGGGTGATGACGAGTGGCGGACGGGTGAGTAATGCATAGGAATCTACCCAGTAGACGGGGAACAACTTGGGGAAACTCAAGCTAATACCGCGTAAGCCTTACGGGGGATAGCGGGGGACTTTTTAGGAAGCCTCGCGATATTGGATGAGCCTATGCCGGATTAGCTAGTTGGTGGGGTAAAAGCTTACCAAGGCTACGATCCGTAGCTGGTCTGAGAGGACGATCAGCCACACTGGGACTGAGACACGGCCCAGACTCCTACGGGAGGCAGCAGTAGGGAATATTGGACAATGGGCGAAAGCTTGATCCAGCAATGCCGCGTGTGTGAAGAAGGCCTGCGGGTTGTAAAGCACTTTCAGTTGGGAAGAAAAGCTTTAGGTTAATACCCTGAAGTATTGACGTTACCAACAGAAGAAGCACCGGCTAACTCTGTGCCAGCAGCCGCGGTAATACAGAGGGTGCGAGCGTTAATCGGAATTACTGGGCGTAAAGCGTACGTAGGCGGTTGAGTCAGTCGGATGTGAAAGCCCAAGGCTTAACCTTGGAACTGCATTCGATACTACTCGGCTAGAGTACAGGAGAGGGAAGCGGAATTCCTAGTGTAGCGGTGAAATGCGTAGAGATTAGGAGGAACACCAGTGGCGAAGGCGGCTTCTTGGACTGATACTGACGCTGAGGTACGAAAGCGTGGGGAGCAAACAGGATTAGATACCCTGGTAGTCCACGCCCTAAACGATGAGAACTAGATGTTGGGGGAATTTAATCCCTTAGTATCGCAGCTAACGCGCTAAGTTCTCCGCCTGGGGAGTACGGCCGCAAGGTTAAAACTCAAAGGAATTGACGGGGGCCCGCACAAGCGGTGGAGCATGTGGTTTAATTCGATGCAACGCGAAGAACCTTACCTGGCCTTGACATCCTTGGAACCTCGCAGAGATGTGAGGGTGCCTTCGGGAACCGAGAGACAGGTGGTGCATGGCTGTCGTCAGCTCGTGTCGTGAGATGTTGGGTTAAGTCCCGCAACGAGCGCAACCCTTATCCCTAGTTGCCAGCGATTCGGTCGGGAACTCTAGAGAGACTGCCGGTGACAAACCGGAGGAAGGTGGGGACGACGTCAAGTCATCATGGCCCTTACGGCCAGGGCTACACACGTGCTACAATGGGGTAGTACAAAGGGTTGCGAACCCGCGAGGGGGTGCTAATCTCACAAAACTACTCGTAGTCCGGATTGGAGTCTGCAACTCGACTCCATGAAGTTGGAATCGCTAGTAATCGCGGATCAGCATGTCGCGGTGAATACGTTCCCGGGCCTTGTACACACCGCCCGTCACACCATGGGAGTGGGCTGTACCAGAAGTAGGTAGTCTAACCGCAAGGGGGACGCTTACCACGGTATGGTTCATGACTGGGGTG

>GuaymasBasin_Dive4872_L-ASV_10029

ATTGAACGCTGGCGGCATGCTTAATACATGCAAGTCGAACGGTAACATGCCCTTCGGGGTGATGACGAGTGGCGGACGGGTGAGTAATGCATAGGAATCTACCCAGTAGACGGGGAACAACTTGGGGAAACTCAAGCTAATACCGCGTAAGCCTTACGGGGGATAGCGGGGGACTTTTTAGGAAGCCTCGCGATATTGGATGAGCCTATGCCGGATTAGCTAGTTGGTGGGGTAAAAGCTTACCAAGGCTACGATCCGTAGCTGGTCTGAGAGGACGATCAGCCACACTGGGACTGAGACACGGCCCAGACTCCTACGGGAGGCAGCAGTAGGGAATATTGGACAATGGGCGAAAGCTTGATCCAGCAATGCCGCGTGGGTGAAGAAGGCCTGCGGGTTGTAAAGCACTTTCAGTTGGGAAGAAAAGCTTTAGGTTAATACCCTGAAGTATTGACGTTACCAACAGAAGAAGCACCGGCTAACTCTGTGCCAGCAGCCGCGGTAATACAGAGGGTGCGAGCGTTAATCGGAATTACTGGGCGTAAAGCGTACGTAGGCGGTTGAGTCAGTCGGATGTGAAAGCCCAAGGCTTAACCTTGGAACTGCATTCGATACTACTCGGCTAGAGTACAGGAGAGGGAAGCGGAATTCCTAGTGTAGCGGTGAAATGCGTAGAGATTAGGAGGAACACCAGTGGCGAAGGCGGCTTCTTGGACTGATACTGACGCTGAGGCACGAAAGCGTGGGGAGCAAACAGGATTAGATACCCTGGTAGTCCACGCCGTAAACGATGAGAACTAGATGTTGGGGGAATTTAATCCCTTAGTATCGCAGCTAACGCGCTAAGTTCTCCGCCTGGGGAGTACGGCCGCAAGGTTAAAACTCAAAGGAATTGACGGGGGCCCGCACAAGCGGTGGAGCATGTGGTTTAATTCGATGCAACGCGAAGAACCTTACCTGGCCTTGACATCCTTGGAACCTCGCAGAGATGTGAGGGTGCCTTCGGGAACCGAGAGACAGGTGCTGCATGGCTGTCGTCAGCTCGTGTCGTGAGATGTTGGGTTAAGTCCCGCAACGAGCGCAACCCTTATCCCTAGTTGCCAGCGATTCGGTCGGGAACTCTAGAGAGACTGCCGGTGACAAACCGGAGGAAGGTGGGGACGACGTCAAGTCATCATGGCCCTTACGGCCAGGGCTACACACGTGCTACAATGGGGTAGTACAAAGGGTTGCGAACCCGCGAGGGGGTGCTAATCTCACAAAACTACTCGTAGTCCGGATTGGAGTCTGCAACTCGACTCCATGAAGTTGGAATCGCTAGTAATCGCGGATCAGCATGTCGCGGTGAATACGTTCCCGGGCCTTGTACACACCGCCCGTCACACCATGGGAGTGGGCTGTACCAGAAGTAGGTAGTCTAACCGCAAGGGGGACGCTTACCACGGTATGGTTCATGACTGGGGTG

>GuaymasBasin_Dive4872_L-ASV_9211

ATTGAACGCTGGCGGCATGCTTAATACATGCAAGTCGAACGGTAACATGCCCTTCGGGGTGATGACGAGTGGCGGACGGGTGAGTAATGCATAGGAATCTACCCAGTAGACGGGGAACAACTTGGGGAAACTCAAGCTAATACCGCGTAAGCCTTACGGGGGATAGCGGGGGACTTTTTAGGAAGCCTCGCGATATTGGATGAGCCTATGCCGGATTAGCTAGTTGGTGGGGTAAAAGCTTACCAAGGCTACGATCCGTAGCTGGTCTGAGAGGACGATCAGCCACACTGGGACTGAGACACGGCCCAGACTCCTACGGGAGGCAGCAGTAGGGAATATTGGACAATGGGCGAAAGCTTGATCCAGCAATGCCGCGTGTGTGAAGAAGGCCTGCGGGTTGTAAAGCACTTTCAGTTGGGAAGAAAAGCTTTAAGTTAATACCCTGAAGTATTGACGTTACCAACAGAAGAAGCACCGGCTAACTCTGTGCCAGCAGCCGCGGTAATACAGAGGGTGCGAGCGTTAATCGGAATTACTGGGCGTAAAGCGTACGTAGGCGGTTGAGTCAGTCGGATGTGAAAGCCCAAGGCTTAACCTTGGAACTGCATTCGATACTACTCGGCTAGAGTACAGGAGAGGGAAGCGGAATTCCTAGTGTAGCGGTGAAATGCGTAGAGATTAGGAGGAACACCAGTGGCGAAGGCGGCTTCTTGGACTGATACTGACGCTGAGGTACGAAAGCGTGGGGAGCAAACAGGATTAGATACCCTGGTAGTCCACGCCCTAAACGATGAGAACTAGATGTTGGGGGAATTTAATCCCTTAGTATCGCAGCTAACGCGCTAAGTTCTCCGCCTGGGGAGTACGGCCGCAAGGTTAAAACTCAAAGGAATTGACGGGGGCCCGCACAAGCGGTGGAGCATGTGGTTTAATTCGATGCAACGCGAAGAACCTTACCTGGCCTTGACATCCTTGGAACCTCGCAGAGATGTGAGGGTGCCTTCGGGAACCGAGAGACAGGTGCTGCATGGCTGTCGTCAGCTCGTGTCGTGAGATGTTGGGTTAAGTCCCGCAACGAGCGCAACCCTTATCCCTAGTTGCCAGCGATTCGGTCGGGAACTCTAGAGAGACTGCCGGTGACAAACCGGAGGAAGGTGGGGATGACGTCAAGTCATCATGGCCCTTACGGCCAGGGCTACACACGTGCTACAATGGGGTAGTACAAAGGGTTGCGAACCCGCGAGGGGGTGCTAATCTCACAAAACTACTCGTAGTCCGGATTGGAGTCTGCAACTCGACTCCATGAAGTTGGAATCGCTAGTAATCGCGGATCAGCATGTCGCGGTGAATACGTTCCCGGGCCTTGTACACACCGCCCGTCACACCATGGGAGTGGGCTGTACCAGAAGTAGGTAGTCTAACCGCAAGGGGGACGCTTACCACGGTATGGTTCATGACTGGGGTG

>GuaymasBasin_Dive4872_L-ASV_2187

ATTGAACGCTGGCGGCATGCTTAATACATGCAAGTCGAACGGTAACATGCCCTTCGGGGTGATGACGAGTGGCGGACGGGTGAGTAATGCATAGGAATCTACCCAGTAGACGGGGAACAACTTGGGGAAACTCAAGCTAATACCGCGTAAGCCTTACGGGGGATAGCGGGGGACTTTTTAGGAAGCCTCGCGATATTGGATGAGCCTATGCCGGATTAGCTAGTTGGTGGGGTAAAAGCTTACCAAGGCTACGATCCGTAGCTGGTCTGAGAGGACGATCAGCCACACTGGGACTGAGACACGGCCCAGACTCCTACGGGAGGCAGCAGTGGGGAATATTGGACAATGGGCGAAAGCCTGATCCAGCAATGCCGCGTGTGTGAAGAAGGCCTGCGGGTTGTAAAGCACTTTCAGTTGGGAAGAAAAGCTTTAGGTTAATACCCTGAAGTATTGACGTTACCAACAGAAGAAGCACCGGCTAACTCTGTGCCAGCAGCCGCGGTAATACAGAGGGTGCGAGCGTTAATCGGAATTACTGGGCGTAAAGCGTACGTAGGCGGTTGAGTCAGTCGGATGTGAAAGCCCAAGGCTTAACCTTGGAACTGCATTCGATACTACTCGGCTAGAGTACAGGAGAGGGAAGCGGAATTCCTAGTGTAGCGGTGAAATGCGTAGAGATTAGGAGGAACACCAGTGGCGAAGGCGGCTTCTTGGACTGATACTGACGCTGAGGTACGAAAGCGTGGGGAGCAAACAGGATTAGATACCCTGGTAGTCCACGCCCTAAACGATGAGAACTAGATGTTGGGGGAATTTAATCCCTTAGTATCGCAGCTAACGCGCTAAGTTCTCCGCCTGGGGAGTACGGCCGCAAGGTTAAAACTCAAAGGAATTGACGGGGGCCCGCACAAGCGGTGGAGCATGTGGTTTAATTCGATGCAACGCGAAGAACCTTACCTGGCCTTGACATCCTTGGAACCTCGCAGAGATGTGAGGGTGCCTTCGGGAACCGAGAGACAGGTGCTGCATGGCTGTCGTCAGCTCGTGTCGTGAGATGTTGGGTTAAGTCCCGCAACGAGCGCAACCCTTATCCCTAGTTGCCAGCGATTCGGTCGGGAACTCTAGAGAGACTGCCGGTGACAAACCGGAGGAAGGTGGGGACGACGTCAAGTCATCATGGCCCTTACGGCCAGGGCTACACACGTGCTACAATGGGGTAGTACAAAGGGTTGCGAACCCGCGAGGGGGTGCTAATCTCACAAAACTACTCGTAGTCCGGATTGGAGTCTGCAACTCGACTCCATGAAGTTGGAATCGCTAGTAATCGCGGATCAGCATGTCGCGGTGAATACGTTCCCGGGCCTTGTACACACCGCCCGTCACACCATGGGAGTGGGCTGTACCAGAAGTAGGTAGTCTAACCGCAAGGGGGACGCTTACCACGGTATGGTTCATGACTGGGGTG

>GuaymasBasin_Dive4872_L-ASV_9514

ATTGAACGCTGGCGGCATGCTTAACACATGCAAGTCGAACGGTAACATGCCCTTCGGGGTGATGACGAGTGGCGGACGGGTGAGTAATGCATAGGAATCTACCCAGTAGACGGGGAACAACTTGGGGAAACTCAAGCTAATACCGCGTAAGCCTTACGGGGGATAGCGGGGGACTTTTTAGGAAGCCTCGCGATATTGGATGAGCCTATGCCGGATTAGCTAGTTGGTGGGGTAAAAGCTTACCAAGGCTACGATCCGTAGCTGGTCTGAGAGGACGATCAGCCACACTGGGACTGAGACACGGCCCAGACTCCTACGGGAGGCAGCAGTGGGGAATATTGGACAATGGGCGAAAGCTTGATCCAGCAATGCCGCGTGTGTGAAGAAGGCCTGCGGGTTGTAAAGCACTTTCAGTTGGGAAGAAAAGCTTTAGGTTAATACCCTGAAGTATTGACGTTACCAACAGAAGAAGCACCGGCTAACTCTGTGCCAGCAGCCGCGGTAATACAGAGGGTGCGAGCGTTAATCGGAATTACTGGGCGTAAAGCGTACGTAGGCGGTTGAGTCAGTCGGATGTGAAAGCCCAAGGCTTAACCTTGGAACTGCATTCGATACTACTCGGCTAGAGTACAGGAGAGGGAAGCGGAATTCCTAGTGTAGCGGTGAAATGCGTAGAGATTAGGAGGAACACCAGTGGCGAAGGCGGCTTCTTGGACTGATACTGACGCTGAGGTACGAAAGCGTGGGGAGCAAACAGGATTAGATACCCTGGTAGTCCACGCCCTAAACGATGAGAACTAGATGTTGGGGGAATTTAATCCCTTAGTATCGCAGCTAACGCGCTAAGTTCTCCGCCTGGGGAGTACGGCCGCAAGGTTAAAACTCAAATGAATTGACGGGGGCCCGCACAAGCGGTGGAGCATGTGGTTTAATTCGATGCAACGCGAAGAACCTTACCTGGCCTTGACATCCTTGGAACCTCGCAGAGATGTGAGGGTGCCTTCGGGAACCGAGAGACAGGTGCTGCATGGCTGTCGTCAGCTCGTGTCGTGAGATGTTGGGTTAAGTCCCGCAACGAGCGCAACCCTTATCCCTAGTTGCCAGCGATTCGGTCGGGAACTCTAGAGAGACTGCCGGTGACAAACCGGAGGAAGGTGGGGACGACGTCAAGTCATCATGGCCCTTACGGCCAGGGCTACACACGTGCTACAATGGGGTAGTACAAAGGGTTGCGAACCCGCGAGGGGGTGCTAATCTCACAAAACTACTCGTAGTCCGGATTGGAGTCTGCAACTCGACTCCATGAAGTTGGAATCGCTAGTAATCGCGGATCAGCATGTCGCGGTGAATACGTTCCCGGGCCTTGTACACACCGCCCGTCACACCATGGGAGTGGGCTGTACCAGAAGTAGGTAGTCTAACCGCAAGGGGGACGCTTACCACGGTATGGTTCATGACTGGGGTG

>GuaymasBasin_Dive4872_L-ASV_10233

ATTGAACGCTGGCGGCATGCTTAATACATGCAAGTCGAACGGTAACATGCCCTTCGGGGTGATGACGAGTGGCGGACGGGTGAGTAATGCATAGGAATCTACCCAGTAGACGGGGAACAACTTGGGGAAACTCAAGCTAATACCGCGTAAGCCTTACGGGGGATAGCGGGGGACTTTTTAGGAAGCCTCGCGATATTGGATGAGCCTATGCCGGATTAGCTAGTTGGTGGGGTAAAAGCTTACCAAGGCTACGATCCGTAGCTGGTCTGAGAGGACGATCAGCCACACTGGGACTGAGACACGGCCCAGACTCCTACGGGAGGCAGCAGTGGGGAATATTGCACAATGGGCGAAAGCTTGATCCAGCAATGCCGCGTGTGTGAAGAAGGCCTGCGGGTTGTAAAGCACTTTCAGTTGGGAAGAAAAGCTTTAGGTTAATACCCTGAAGTATTGACGTTACCAACAGAAGAAGCACCGGCTAACTCTGTGCCAGCAGCCGCGGTAATACAGAGGGTGCGAGCGTTAATCGGAATTACTGGGCGTAAAGCGTACGTAGGCGGTTGAGTCAGTCGGATGTGAAAGCCCAAGGCTTAACCTTGGAACTGCATTCGATACTACTCGGCTAGAGTACAGGAGAGGGAAGCGGAATTCCTAGTGTAGCGGTGAAATGCGTAGAGATTAGGAGGAACACCAGTGGCGAAGGCGGCTTCTTGGACTGATACTGACGCTGAGGTACGAAAGCGTGGGGAGCAAACAGGATTAGATACCCTGGTAGTCCACGCCCTAAACGATGAGAACTAGATGTTGGGGGAATTTAATCCCTTAGTATCGCAGCTAACGCGCTAAGTTCTCCGCCTGGGGAGTACGGCCGCAAGGTTAAAACTCAAATGAATTGACGGGGGCCCGCACAAGCGGTGGAGCATGTGGTTTAATTCGATGCAACGCGAAGAACCTTACCTGGCCTTGACATCCTTGGAACCTCGCAGAGATGTGAGGGTGCCTTCGGGAACCGAGAGACAGGTGCTGCATGGCTGTCGTCAGCTCGTGTCGTGAGATGTTGGGTTAAGTCCCGCAACGAGCGCAACCCTTATCCCTAGTTGCCAGCGATTCGGTCGGGAACTCTAGAGAGACTGCCGGTGACAAACCGGAGGAAGGTGGGGACGACGTCAAGTCATCATGGCCCTTACGGCCAGGGCTACACACGTGCTACAATGGGGTAGTACAAAGGGTTGCGAACCCGCGAGGGGGTGCTAATCTCACAAAACTACTCGTAGTCCGGATTGGAGTCTGCAACTCGACTCCATGAAGTTGGAATCGCTAGTAATCGCGGATCAGCATGTCGCGGTGAATACGTTCCCGGGCCTTGTACACACCGCCCGTCACACCATGGGAGTGGGCTGTACCAGAAGTAGGTAGTCTAACCGCAAGGGGGACGCTTACCACGGTATGGTTCATGACTGGGGTG

>GuaymasBasin_Dive4872_L-ASV_9582

ATTGAACGCTGGCGGCATGCTTAATACATGCAAGTCGAACGGTAACATGCCCTTCGGGGTGATGACGAGTGGCGGACGGGTGAGTAATGCATAGGAATCTACCCAGTAGACGGGGAACAACTTGGGGAAACTCAAGCTAATACCGCGTAAGCCTTACGGGGGATAGCGGGGGACTTTTTAGGAAGCCTCGCGATATTGGATGAGCCTATGCCGGATTAGCTAGTTGGTGGGGTAAAAGCTTACCAAGGCTACGATCCGTAGCTGGTCTGAGAGGACGATCAGCCACACTGGGACTGAGACACGGCCCAGACTCCTACGGGAGGCAGCAGTAGGGAATATTGGACAATGGGCGAAAGCTTGATCCAGCAATGCCGCGTGTGTGAAGAAGGCCTGCGGGTTGTAAAGCACTTTCAGTTGGGAAGAAAAGCTTTAGGTTAATACCCTGAAGTATTGACGTTACCAACAGAAGAAGCACCGGCTAACTCTGTGCCAGCAGCCGCGGTAATACAGAGGGTGCGAGCGTTAATCGGAATTACTGGGCGTAAAGCGTACGTAGGCGGTTGAGTCAGTCGGATGTGAAAGCCCAAGGCTTAACCTTGGAACTGCATTCGATACTACTCGGCTAGAGTACAGGAGAGGGAAGCGGAATTCCTAGTGTAGCGGTGAAATGCGTAGAAATTAGGAGGAACACCAGTGGCGAAGGCGGCTTCTTGGACTGATACTGACGCTGAGGTACGAAAGCGTGGGGAGCAAACAGGATTAGATACCCTGGTAGTCCACGCCCTAAACGATGAGAACTAGATGTTGGGGGAATTTAATCCCTTAGTATCGCAGCTAACGCGCTAAGTTCTCCGCCTGGGGAGTACGGCCGCAAGGTTAAAACTCAAATGAATTGACGGGGGCCCGCACAAGCGGTGGAGCATGTGGTTTAATTCGATGCAACGCGAAGAACCTTACCTGGCCTTGACATCCTTGGAACCTCGCAGAGATGTGAGGGTGCCTTCGGGAACCGAGAGACAGGTGCTGCATGGCTGTCGTCAGCTCGTGTCGTGAGATGTTGGGTTAAGTCCCGCAACGAGCGCAACCCTTATCCCTAGTTGCCAGCGATTCGGTCGGGAACTCTAGAGAGACTGCCGGTGACAAACCGGAGGAAGGTGGGGACGACGTCAAGTCATCATGGCCCTTACGGCCAGGGCTACACACGTGCTACAATGGGGTAGTACAAAGGGTTGCGAACCCGCGAGGGGGTGCTAATCTCACAAAACTACTCGTAGTCCGGATTGGAGTCTGCAACTCGACTCCATGAAGTTGGAATCGCTAGTAATCGCGGATCAGCATGTCGCGGTGAATACGTTCCCGGGCCTTGTACACACCGCCCGTCACACCATGGGAGTGGGCTGTACCAGAAGTAGGTAGTCTAACCGCAAGGGGGACGCTTACCACGGTATGGTTCATGACTGGGGTG

>GuaymasBasin_Dive4872_L-ASV_9577

ATTGAACGCTGGCGGCATGCTTAATACATGCAAGTCGAACGGTAACATGCCCTTCGGGGTGATGACGAGTGGCGGACGGGTGAGTAATGCATAGGAATCTACCCAGTAGACGGGGAACAACTTGGGGAAACTCAAGCTAATACCGCGTAAGCCTTACGGGGGATAGCGGGGGACTTTTTAGGAAGCCTCGCGATATTGGATGAGCCTATGCCGGATTAGCTAGTTGGTGGGGTAAAAGCTTACCAAGGCTACGATCCGTAGCTGGTCTGAGAGGATGATCAGCCACACTGGGACTGAGACACGGCCCAGACTCCTACGGGAGGCAGCAGTAGGGAATATTGGACAATGGGCGAAAGCTTGATCCAGCAATGCCGCGTGTGTGAAGAAGGCCTGCGGGTTGTAAAGCACTTTCAGTTGGGAAGAAAAGCTTTAGGTTAATACCCTGAAGTATTGACGTTACCAACAGAAGAAGCACCGGCTAACTCTGTGCCAGCAGCCGCGGTAATACAGAGGGTGCGAGCGTTAATCGGAATTACTGGGCGTAAAGCGTACGTAGGCGGTTGAGTCAGTCGGATGTGAAAGCCCAAGGCTTAACCTTGGAACTGCATTCGATACTACTCGGCTAGAGTACAGGAGAGGGAAGCGGAATTCCTAGTGTAGCGGTGAAATGCGTAGAGATTAGGAGGAACACCAGTGGCGAAGGCGGCTTCTTGGACTGATACTGACGCTGAGGTACGAAAGCGTGGGGAGCAAACAGGATTAGATACCCTGGTAGTCCACGCCCTAAACGATGAGAACTAGATGTTGGGGGAATTTAATCCCTTAGTATCGCAGCTAACGCGCTAAGTTCTCCGCCTGGGGAGTACGGCCGCAAGGTTAAAACTCAAATGAATTGACGGGGGCCCGCACAAGCGGTGGAGCATGTGGTTTAATTCGATGCAACGCGAAGAACCTTACCTGGCCTTGACATCCTTGGAACCTCGCAGAGATGTGAGGGTGCCTTCGGGAACCGAGAGACAGGTGCTGCATGGCTGTCGTCAGCTCGTGTCGTGAGATGTTGGGTTAAGTCCCGCAACGAGCGCAACCCTTATCCCTAGTTGCCAGCGATTCGGTCGGGAACTCTAGAGAGACTGCCGGTGACAAACCGGAGGAAGGTGGGGACGACGTCAAGTCATCATGGCCCTTACGGCCAGGGCTACACACGTGCTACAATGGGGTAGTACAAAGGGTTGCGAACCCGCGAGGGGGTGCTAATCTCACAAAACTACTCGTAGTCCGGATTGGAGTCTGCAACTCGACTCCATGAAGTTGGAATCGCTAGTAATCGCGGATCAGCATGTCGCGGTGAATACGTTCCCGGGCCTTGTACACACCGCCCGTCACACCATGGGAGTGGGCTGTACCAGAAGTAGGTAGTCTAACCGCAAGGGGGACGCTTACCACGGTATGGTTCATGACTGGGGTG

>GuaymasBasin_Dive4872_L-ASV_133

ATTGAACGCTGGCGGCATGCTTAATACATGCAAGTCGAACGGTAACATGCCCTTCGGGGTGATGACGAGTGGCGGACGGGTGAGTAATGCATAGGAATCTACCCAGTAGACGGGGAACAACTTGGGGAAACTCAAGCTAATACCGCGTAAGCCTTACGGGGGATAGCGGGGGACTTTTTAGGAAGCCTCGCGATATTGGATGAGCCTATGCCGGATTAGCTAGTTGGTGGGGTAAAAGCTTACCAAGGCTACGATCCGTAGCTGGTCTGAGAGGACGATCAGCCACACTGGGACTGAGACACGGCCCAGACTCCTACGGGAGGCAGCAGTAGGGAATATTGGACAATGGGCGAAAGCTTGATCCAGCAATGCCGCGTGTGTGAAGAAGGCCTGCGGGTTGTAAAGCACTTTCAGTTGGGAAGAAAAGCTTTAGGTTAATACCCTGAAGTATTGACGTTACCAACAGAAGAAGCACCGGCTAACTCTGTGCCAGCAGCCGCGGTAATACAGAGGGTGCGAGCGTTAATCGGAATTACTGGGCGTAAAGCGTACGTAGGCGGTTGAGTCAGTCGGATGTGAAAGCCCAAGGCTTAACCTTGGAACTGCATTCGATACTACTCGGCTAGAGTACAGGAGAGGGAAGCGGAATTCCTAGTGTAGCGGTGAAATGCGTAGAGATTAGGAGGAACACCAGTGGCGAAGGCGGCTTCTTGGACTGATACTGACGCTGAGGTACGAAAGCGTGGGGAGCAAACAGGATTAGATACCCTGGTAGTCCACGCCCTAAACGATGAGAACTAGATGTTGGGGGAATTTAATCCCTTAGTATCGCAGCTAACGCGCTAAGTTCTCCGCCTGGGGAGTACGGCCGCAAGGTTAAAACTCAAATGAATTGACGGGGGCCCGCACAAGCGGTGGAGCATGTGGTTTAATTCGATGCAACGCGAAGAACCTTACCTGGCCTTGACATCCTTGGAACCTCGCAGAGATGTGAGGGTGCCTTCGGGAACCGAGAGACAGGTGCTGCATGGCTGTCGTCAGCTCGTGTCGTGAGATGTTGGGTTAAGTCCCGCAACGAGCGCAACCCTTATCCCTAGTTGCCAGCGATTCGGTCGGGAACTCTAGAGAGACTGCCGGTGACAAACCGGAGGAAGGTGGGGACGACGTCAAGTCATCATGGCCCTTACGGCCAGGGCTACACACGTGCTACAATGGGGTAGTACAAAGGGTTGCGAACCCGCGAGGGGGTGCTAATCTCACAAAACTACTCGTAGTCCGGATTGGAGTCTGCAACTCGACTCCATGAAGTTGGAATCGCTAGTAATCGCGGATCAGCATGTCGCGGTGAATACGTTCCCGGGCCTTGTACACACCGCCCGTCACACCATGGGAGTGGGCTGTACCAGAAGTAGGTAGTCTAACCGCAAGGGGGACGCTTACCACGGTATGGTTCATGACTGGGGTG

>GuaymasBasin_Dive4872_L-ASV_9396

ATTGAACGCTGGCGGCATGCTTAATACATGCAAGTCGAACGGTAACATGCCCTTCGGGGTGATGACGAGTGGCGGACGGGTGAGTAATGCATAGGAATCTACCCAGTAGACGGGGAACAACTTGGGGAAACTCAAGCTAATACCGCGTAAGCCTTACGGGGGATAGCGGGGGACTTTTTAGGAAGCCTCGCGATATTGGATGAGCCTATGCCGGATTAGCTAGTTGGTGGGGTAAAAGCTTACCAAGGCTACGATCCGTAGCTGGTCTGAGAGGACGATCAGCCACACTGGGACTGAGACACGGCCCAGACTCCTACGGGAGGCAGCAGTAGGGAATATTGGACAATGGGCGAAAGCTTGATCCAGCAATGCCGCGTGTGTGAAGAAGGCCTGCGGGTTGTAAAGCACTTTCAGTTGGGAAGAAAAGCTTTAGGTTAATACCCTGAAGTATTGACGTTACCAACAGAAGAAGCACCGGCTAATTCTGTGCCAGCAGCCGCGGTAATACAGAGGGTGCGAGCGTTAATCGGAATTACTGGGCGTAAAGCGTACGTAGGCGGTTGAGTCAGTCGGATGTGAAAGCCCAAGGCTTAACCTTGGAACTGCATTCGATACTACTCGGATAGAGTACAGGAGAGGGAAGCGGAATTCCTAGTGTAGCGGTGAAATGCGTAGAGATTAGGAGGAACACCAGTGGCGAAGGCGGCTTCTTGGACTGATACTGACGCTGAGGTACGAAAGCGTGGGGAGCAAACAGGATTAGATACCCTGGTAGTCCACGCCCTAAACGATGAGAACTAGATGTTGGGGGAATTTAATCCCTTAGTATCGCAGCTAACGCGCTAAGTTCTCCGCCTGGGGAGTACGGCCGCAAGGTTAAAACTCAAATGAATTGACGGGGGCCCGCACAAGCGGTGGAGCATGTGGTTTAATTCGATGCAACGCGAAGAACCTTACCTGGCCTTGACATCCTTGGAACCTCGCAGAGATGTGAGGGTGCCTTCGGGAACCGAGAGACAGGTGCTGCATGGCTGTCGTCAGCTCGTGTCGTGAGATGTTGGGTTAAGTCCCGCAACGAGCGCAACCCTTATCCCTAGTTGCCAGCGATTCGGTCGGGAACTCTAGAGAGACTGCCGGTGACAAACCGGAGGAAGGTGGGGACGACGTCAAGTCATCATGGCCCTTACGGCCAGGGCTACACACGTGCTACAATGGGGTAGTACAAAGGGTTGCGAACCCGCGAGGGGGTGCTAATCTCACAAAACTACTCGTAGTCCGGATTGGAGTCTGCAACTCGACTCCATGAAGTTGGAATCGCTAGTAATCGCGGATCAGCATGTCGCGGTGAATACGTTCCCGGGCCTTGTACACACCGCCCGTCACACCATGGGAGTGGGCTGTACCAGAAGTAGGTAGTCTAACCGCAAGGGGGACGCTTACCACGGTATGGTTCATGACTGGGGTG

>GuaymasBasin_Dive4872_L-ASV_9095

ATTGAACGCTGGCGGCATGCTTAATACATGCAAGTCGAACGGTAACATGCCCTTCGGGGTGATGACGAGTGGCGGACGGGTGAGTAATGCATAGGAATCTACCCAGTAGACGGGGAACAACTTGGGGAAACTCAAGCTAATACCGCGTAAGCCTTACGGGGGATAGCGGGGGACTTTTTAGGAAGCCTCGCGATATTGGATGAGCCTATGCCGGATTAGCTAGTTGGTGGGGTAAAAGCTTACCAAGGCTACGATCCGTAGCTGGTCTGAGAGGACGATCAGCCACACTGGGACTGAGACACGGCCCAGACTCCTACGGGAGGCAGCAGTAGGGAATATTGGACAATGGGCGAAAGCTTGATCCAGCAATGCCGCGTGTGTGAAGAAGGCCTGCGGGTTGTAAAGCACTTTCAGTTGGGAAGAAAAGCTTTAGGTTAATACCCTGAAGTATTGACGTTACCAACAGAAGAAGCACCGGCTAACTCTGTGCCAGCAGCCGCGGTAATACAGAGGGTGCGAGCGTTAATCGGAATTACTGGGCGTAAAGCGTACGTAGGCGGTTGAGTCAGTCGGATGTGAAAGCCCAAGGCTTAACCTTGGAACTGCATTCGATACTACTCGGCTAGAGTACAGGAGAGGGAAGCGGAATTCCTAGTGTAGCGGTGAAATGCGTAGAGATTAGGAGGAACACCAGTGGCGAAGGCGGCTTCTTGGACTGATACTGACGCTGAGGTACGAAAGCGTGGGGAGCAAACAGGATTAGATACCCTGGTAGTCCACGCCCTAAACGATGAGAACTAGATGTTGGGGGAATTTAATCCCTTAGTATCGCAGCTAACGCGCTAAGTTCTCCGCCTGGGGAGTACGGCCGCAAGGTTAAAACTCAAAGGAATTGACGGGGGCCCGCACAAGCGGTGGAGCATGTGGTTTAATTCGATGCAACGCGAAGAACCTTACCTGGCCTTGACATCCTTGGAACCTCGCAGAGATGTGAGGGTGCCTTCGGGAACCGAGAGACAGGTGCTGCATGGCTGTCGTCAGCTCGTGTCGTGAGATGTTGGGTTAAGTCCCGCAACGAGCGCAACCCTTATCCCTAGTTGCCAGCGATTCGGTCGGGAACTCTAGAGAGACTGCCGGTGACAAACCGGAGGAAGGTGGGGACGACGTCAAGTCATCATGGCCCTTACGGCCAGGGCTACACACGTGCTACAATGGGGTAGTACAAAGGGTTGCGAACCCGCGAGGGGGTGCTAATCTCACAAAACTACTCGTAGTCCGGATTGGAGTCTGCAACTCGACTCCATGAAGTTGGAATCGCTAGTAATCGCGGATCAGCATGTCGCGGTGAATACGTTCCCGGGCCTTGTACACACCGCCCGTCACACCA

>GuaymasBasin_Dive4872_L-ASV_9595

ATTGAACGCTGGCGGCATGCTTAATACATGCAAGTCGAACGGTAACATGCCCTTCGGGGTGATGACGAGTGGCGGACGGGTGAGTAATGCATAGGAATCTACCCAGTAGACGGGGAACAACTTGGGGAAACTCAAGCTAATACCGCGTAAGCCTTACGGGGGATAGCGGGGGACTTTTTAGGAAGCCTCGCGATATTGGATGAGCCTATGCCGGATTAGCTAGTTGGTGGGGTAAAAGCTTACCAAGGCTACGATCCGTAGCTGGTCTGAGAGGACGATCAGCCACACTGGGACTGAGACACGGCCCAGACTCCTACGGGAGGCAGCAGTAGGGAATATTGGACAATGGGCGAAAGCTTGATCCAGCAATGCCGCGTGTGTGAAGAAGGCCTGCGGGTTGTAAAGCACTTTCAGTTGGGAAGAAAAGCTTTAGGTTAATACCCTGAAGTATTGACGTTACCAACAGAAGAAGCACCGGCTAACTCTGTGCCAGCAGCCGCGGTAATACAGAGGGTGCGAGCGTTAATCGGAATTACTGGGCGTAAAGCGTACGTAGGCGGTTGAGTCAGTCGGATGTGAAAGCCCAAGGCTTAACCTTGGAACTGCATTCGATACTACTCGGCTAGAGTACAGGAGAGGGAAGCGGAATTCCTAGTGTAGCGGTGAAATGCGTAGAGATTAGGAGGAACACCAGTGGCGAAGGCGGCTTCTTGGACTGATACTGACGCTGAGGTACGAAAGCGTGGGGAGCAAACAGGATTAGATACCCTGGTAGTCCACGCCCTAAACGATGAGAACTAGATGTTGGGGGAATTTAATCCCTTAGTATCGCAGCTAACGCGCTAAGTTCTCCGCCTGGGGAGTACGGCCGCAAGGTTAAAACTCAAAGGAATTGACGGGGGCCCGCACAAGCGGTGGAGCATGTGGTTTAATTCGATGCAACGCGAAGAACCTTACCTGGCCTTGACATCCTTGGAACCTCGCAGAGATGTGAGGGTGCCTTCGGGAACCGAGAGACAGGTGCTGCATGGCTGTCGTCAGCTCGTGTCGTGAGATGTTGGGTTAAGTCCCGCAACGAGCGCAACCCTTATCCCTAGTTGCCAGCGATTCGGTCGGGAACTCTAGAGAGACTGCCGGTGACAAACCGGAGGAAGGTGGGGACGACGTCAAGTCATCATGGCCCTTACGGCCAGGGCTACACACGTGCTACAATGGGGTAGTACAAAGGGTTGCGAACCCGCGAGGGGGTGCTAATCTCACAAAACTACTCGTAGTCCGGATTGGAGTCTGCAACTCGACTCCATGAAGCTGGAATCGCTAGTAATCGCGGATCAGCATGTCGCGGTGAAT

>GuaymasBasin_Dive4872_L-ASV_12

AGTGAACGCTAGGGGCGTACCTAACACATGCAAGTCGTGCGAGAAACCTACCTTTCGGGGTAGGGAGTAAAGCGGCGAACGGGTGAGTAATGCGTAGGTAACCTGCCTCCGAGAGGGGAATAACCTACCGAAAGGTGGGCTAATGCCCCATAAGCTCACGATCACTACGGTGGTTGTGAGAAAAGATGGCCTCTGCATGCAAGCTGTCGCTCGGAGAGGGGCTTACGTCCTATCAGCTAGTTGGTGAGGTAATGGCTCACCAAGGCTACGACGGGTAGCCGGCCTGAGAGGGTGGTCGGCCACACTGGCACTGAGACACGGGCCAGACTCCTACGGGAGGCAGCAGTGGGGAATATTGGGCAATGGGCGCAAGCCTGACCCAGTAACGCCCCGTGGGTGATGAAGGCCTTCGGGTCGTAAAGCCCTGTCAGGTGGGAAGAATATCTGATGGGTAAATAGCCCATTAGATTGACGGTACCACCAAAGGAAGCACCGGCTAACTCCGTGCCAGCAGCCGCGGTAATACGGAGGGTGCAAGCGTTACTCGGAATCACTGGGCGTAAAGGGCGCGCAGGCGGGAAGGCAAGTTGAGCGTGTAAGCCTGAGGCTCAACCTCAGAATGGCGCTCAAAACTGCCTTTCTTGAGTCCCGGAGAGGCCGGCGGAATTCCCGGTGTAGGGGTGAAATCCGTAGATATCGGGAGGAACACCGGTGGCGAAGGCGGCCGGCTGGACGGGTACTGACGCTGAGGCGCGAAAGCGTGGGGAGCAAACAGGATTAGATACCCTGGTAGTCCACGCCGTAAACGATGAGCACTAGGTGTGGGGGAGATTATCTCTTCCGTGCCGCAGCTAACGCATTAAGTGCTCCGCCTGGGGAGTACGGCCGCAAGGTTAAAACTCAAAGGAATTGACGGGGGCCCGCACAAGCGGTGGAGCATGTGGTTTAATTCGATGCTACGCGAAGAATCTTACCTGGGTTTGACATCCCCGGAACCCTGCCGAAAGGTGGGGGTGCCCCTTCGGGGGAACCGGGTGACAGGTGCTGCATGGCTGTCGTCAGCTCGTGTCGTGAGATGTTGGGTTAAGTCCCGCAACGAGCGCAACCCTTGCCCTTAGTTGCCAGCGGTTCGGCCGGGCACTCTAAGGGGACTGCCGGCGATAAGCCGGAGGAAGGTGGGGATGACGTCAAGTCATCATGGCCCTTATGCCCAGGGCTACACACGTGCTACAATGGCTGGTACAAAGGGTCGCGAAACCGCGAGGTGGAGCTAATCCCAAAAAGCCAGTCCCAGTCCGGATCGAAGGCTGGAACTCGCCTTCGTGAAGGCGGAATCGCTAGTAATGGCGGATCAGAATGCCGCCGTGAATACGTTCCCGGGCCTTGTACACACCGCCCGTCACACCACGAAAGTCGGCTGTACCAGAAGTCGCTGGCCCAACCCCGTAAGGGGAGGGAGGCGCCCAAGGTGTGGTCGGCGATTGGGGTG

>GuaymasBasin_Dive4872_L-ASV_899

AGTGAACGCTAGGGGCGTACCTAACACATGCAAGTCGTGCGAGAAACCTACCTTTCGGGGTAGGGAGTAAAGCGGCGAACGGGTGAGTAATGCGTAGGTAACCTGCCTCCGAGAGGGGAATAACCTACCGAAAGGTGGGCTAATGCCCCATAAGCTCACGATCACTACGGTGGTTGTGAGAAAAGATGGCCTCTGCATGCAAGCTGTCGCTCGGAGAGGGGCTTACGTCCTATCAGCTAGTTGGTGAGGTAATGGCTCACCAAGGCTACGACGGGTAGCCGGCCTGAGAGGGTGGTCGGCCACACTGGCACTGAGACACGGGCCAGACTCCTACGGGAGGCAGCAGTGGGGAATATTGGGCAATGGGCGCAAGCCTGACCCAGTAACGCCCCGTGGGTGATGAAGGCCTTCGGGTCGTAAAGCCCTGTCAGGTGGGAAGAATATCTGATGGGTAAATAGCCCATTAGATTGACGGTACCACCAAAGGAAGCACCGGCTAACTCCGTGCCAGCAGCCGCGGTAATACGGAGGGTGCAAGCGTTACTCGGAATCACTGGGCGTAAAGGGCGCGCAGGCGGGAAGGCAAGTTGAGCGTGTAAGCCTGAGGCTCAACCTCAGAATGGCGCTCAAAACTGCCTTTCTTGAGTCCCGGAGAGGCCGGCGGAATTCCCGGTGTAGGGGTGAAATCCGTAGATATCGGGAGGAACACCGGTGGCGAAGGCGGCCGGCTGGACGGGTACTGACGCTGAGGCGCGAAAGCGTGGGGAGCAAACAGGATTAGATACCCTGGTAGTCCACGCCGTAAACGATGAGCACTAGGTGTGGGGGAGATTATCTCTTCCGTGCCGGAGCTAACGCATTAAGTGCTCCGCCTGGGGAGTACGGCCGCAAGGTTAAAACTCAAAGGAATTGACGGGGGCCCGCACAAGCGGTGGAGCATGTGGTTTAATTCGATGCTACGCGAAGAATCTTACCTGGGTTTGACATCCCCGGAACCCTGCCGAAAGGTGGGGGTGCCCCTTCGGGGGAACCGGGTGACAGGTGCTGCATGGCTGTCGTCAGCTCGTGTCGTGAGATGTTGGGTTAAGTCCCGCAACGAGCGCAACCCTTGCCCTTAGTTGCCAGCGGTTCGGCCGGGCACTCTAAGGGGACTGCCGGCGATAAGCCGGAGGAAGGTGGGGATGACGTCAAGTCATCATGGCCCTTATGCCCAGGGCTACACACGTGCTACAATGGCTGGTACAAAGGGTCGCGAAACCGCGAGGTGGAGCTAATCCCAAAAAGCCAGGCCCAGTCCGGATCGAAGGCTGGAACTCGCCTTCGTGAAGGCGGAATCGCTAGTAATGGCGGATCAGAATGCCGCCGTGAATACGTTCCCGGGCCTTGTACACACCGCCCGTCACACCACGAAAGTCGGCTGTACCAGAAGTCGCTGGCCCAACCCCGTAAGGGGAGGGAGGCGCCCAAGGTGTGGTCGGCGATTGGGGTG

>GuaymasBasin_Dive4872_L-ASV_106

AGTGAACGCTAGGGGCGTACCTAACACATGCAAGTCGTGCGAGAAACCTACCTTTCGGGGTAGGGAGTAAAGCGGCGAACGGGTGAGTAATGCGTAGGTAACCTGCCTCCGAGAGGGGAATAACCTACCGAAAGGTGGGCTAATGCCCCATAAGCTCACGATCACTACGGTGGTTGTGAGAAAAGATGGCCTCTGCATGCAAGCTGTCGCTCGGAGAGGGGCTTACGTCCTATCAGCTAGTTGGTGAGGTAATGGCTCACCAAGGCTACGACGGGTAGCCGGCCTGAGAGGGTGGTCGGCCACACTGGCACTGAGACACGGGCCAGACTCCTACGGGAGGCAGCAGTGGGGAATATTGGGCAATGGGCGCAAGCCTGACCCAGTAACGCCCCGTGGGTGATGAAGGCCTTCGGGTCGTAAAGCCCTGTCAGGTGGGAAGAATATCTGATGGGTAAATAGCCCATTAGATTGACGGTACCACCAAAGGAAGCACCGGCTAACTCCGTGCCAGCAGCCGCGGTAATACGGAGGGTGCAAGCGTTACTCGGAATCACTGGGCGTAAAGGGCGCGCAGGCGGGAAGGCAAGTTGAGCGTGTAAGCCTGAGGCTCAACCTCAGAATGGCGCTCAAAACTGCCTTTCTTGAGTCCCGGAGAGGCCGGCGGAATTCCCGGTGTAGGGGTGAAATCCGTAGATATCGGGAGGAACACCGGTGGCGAAGGCGGCCGGCTGGACGGGTACTGACGCTGAGGCGCGAAAGCGTGGGGAGCAAACAGGATTAGATACCCTGGTAGTCCACGCCGTAAACGATGAGCACTAGGTGTGGGGGAGATTATCTCTTCCGTGCCGCAGCTAACGCATTAAGTGCTCCGCCTGGGGAGTACGGCCGCAAGGTTAAAACTCAAAGGAATTGACGGGGGCCCGCACAAGCGGTGGAGCATGTGGTTTAATTCGATGCAACGCGAAGAACCTTACCTGGGTTTGACATCCCCGGAACCCTGCCGAAAGGTGGGGGTGCCCCTTCGGGGGAACCGGGTGACAGGTGCTGCATGGCTGTCGTCAGCTCGTGTCGTGAGATGTTGGGTTAAGTCCCGCAACGAGCGCAACCCTTGCCCTTAGTTGCCAGCGGTTCGGCCGGGCACTCTAAGGGGACTGCCGGCGATAAGCCGGAGGAAGGTGGGGATGACGTCAAGTCATCATGGCCCTTATGCCCAGGGCTACACACGTGCTACAATGGCTGGTACAAAGGGTCGCGAAACCGCGAGGTGGAGCTAATCCCAAAAAGCCAGTCCCAGTCCGGATCGAAGGCTGGAACTCGCCTTCGTGAAGGCGGAATCGCTAGTAATGGCGGATCAGAATGCCGCCGTGAATACGTTCCCGGGCCTTGTACACACCGCCCGTCACACCACGAAAGTCGGCTGTACCAGAAGTCGCTGGCCCAACCCCGTAAGGGGAGGGAGGCGCCCAAGGTGTGGTCGGCGATTGGGGTG

>GuaymasBasin_Dive4872_L-ASV_17

AGTGAACGCTAGGGGCGTACCTAACACATGCAAGTCGTGCGAGAAACCTATCCTTCAGGGTAGGGAGTAAAGCGGCGGACGGGTGAGTAATGCGTAGGTAACCTGCCTCCGAGAGGGGAATAACCTACCGAAAGGTGGGCTAATGCCCCATAAGCTCACGGCCACTACGGTGGTTGTGAGAAAAGATGGCCTCTGCATGCAAGCTGTCGCTCGGAGAGGGGCTTACGTCCTATCAGCTAGTTGGTGAGGTAATGGCTCACCAAGGCTACGACGGGTAGCCGGCCTGAGAGGGTGGTCGGCCACACTGGCACTGAGACACGGGCCAGACTCCTACGGGAGGCAGCAGTGGGGAATATTGGGCAATGGGCGCAAGCCTGACCCAGTAACGCCCTGTGGGTGATGAAGGCCTTCGGGTCGTAAAGCCCTGTCAGGTGGGAAGAATATCTAATGGATGAATAGTCCATTAGATTGACGGTACCACCAAAGGAAGCACCGGCTAACTCCGTGCCAGCAGCCGCGGTAATACGGAGGGTGCAAGCGTTACTCGGAATTACTGGGCGTAAAGGGCGCGCAGGCGGGAAGGCAAGTTGAGCGTGTAAGCCTGAGGCTCAACCTCAGAATGGCGCTCAAAACTGCCTTTCTTGAGTCCCGGAGAGGCCGGCGGAATTCCCGGTGTAGGGGTGAAATCCGTAGATATCGGGAGGAACACCGGTGGCGAAGGCGGCCGGCTGGACGGGTACTGACGCTGAGGCGCGAAAGCGTGGGGAGCAAACAGGATTAGATACCCTGGTAGTCCACGCCGTAAACGATGAGCACTAGGTGTGGGGGAGATTATCTCTTCCGTGCCGCAGCTAACGCATTAAGTGCTCCGCCTGGGGAGTACGGCCGCAAGGTTAAAACTCAAAGGAATTGACGGGGGCCCGCACAAGCGGTGGAGCATGTGGTTTAATTCGATGCTACGCGAAGAATCTTACCTGGGTTTGACATCCCCGGAACCTTGCCGAAAGGTGAGGGTGCTCCTTCGGGAGAACCGGGTGACAGGTGCTGCATGGCTGTCGTCAGCTCGTGTCGTGAGATGTTGGGTTAAGTCCCGCAACGAGCGCAACCCTTGCCCTTAGTTGCCAGCGGTTTGGCCGGGCACTCTAAGGGGACTGCCGGCGATAAGCCGGAGGAAGGTGGGGATGACGTCAAGTCATCATGGCCCTTATGCCCAGGGCTACACACGTGCTACAATGGCTGGTACAAAGGGTCGCGAAACCGCAAGGTGGAGCTAATCCCAAAAAGCCAGTCCCAGTTCGGATCGAAGGCTGTAACTCGCCTTCGTGAAGGCGGAATCGCTAGTAATGGCGGATCAGAATGCCGCCGTGAATACGTTCCCGGGCCTTGTACACACCGCCCGTCACACCACGAAAGTCGGCTGTACCAGAAGTCGCTGGCCCAACCCCGTAAGGGGAGGGAGGCGCCCAAGGTGTGGTTGGCGATTGGGGTG

>GuaymasBasin_Dive4872_L-ASV_53

AGTGAACGCTAGGGGCGTACCTAACACATGCAAGTCGTGCGAGAAACCTACCCTTCCGGGGTAGGGAGTAAAGCGGCGGACGGGTGAGTAATGCGTAGGTAACCTGCCTCCGAGCGGGGAATAACCTACCGAAAGGTGGGCTAATGCCCCATAAGCTCACGGCCACTACGGTGGTTGTGAGAAAAGATGGCCTCTGCATGCAAGCTGTCGCTCGGAGAGGGGCTTACGTCCTATCAGCTAGTTGGTGAGGTAATGGCTCACCAAGGCTACGACGGGTAGCCGGCCTGAGAGGGTGGTCGGCCACACTGGCACTGAGACACGGGCCAGACTCCTACGGGAGGCAGCAGTGGGGAATATTGGGCAATGGGCGCAAGCCTGACCCAGTAACGCCCCGTGGGTGATGAAGGCCTTCGGGTCGTAAAGCCCTGTCAGGTGGGAAGAATATCTGATGGATGAATAGTCCATTAGATTGACGGTACCACCAGAGGAAGCACCGGCTAACTCCGTGCCAGCAGCCGCGGTAATACGGAGGGTGCGAGCGTTACTCGGAATCACTGGGCGTAAAGGGCGCGTAGGCGGGAAGGCCAGTTGAGCGTGTAAGCCTGAGGCTTAACCTCAGAATGGCGCTCAAAACTGCCTTTCTTGAGTCCCGGAGAGGCCGGCGGAATTCCCGGTGTAGGGGTGAAATCCGTAGATATCGGGAGGAACACCGGTGGCGAAGGCGGCCGGCTGGACGGGTACTGACGCTGAGGCGCGAAAGCGTGGGGAGCAAACAGGATTAGATACCCTGGTAGTCCACGCCGTAAACGATGAGCACTAGGTGTGGGAGAGATCATCTCTTCCGTGCCGCAGCTAACGCATTAAGTGCTCCGCCTGGGGAGTACGGCCGCAAGGTTAAAACTCAAAGGAATTGACGGGGGCCCGCACAAGCGGTGGAGCATGTGGTTTAATTCGATGCTACGCGAAGAATCTTACCTGGGCTTGACATCCCCGGAACCTTGCCGAAAGGTAAGGGTGCTCCTTCGGGAGAACCGGGTGACAGGTGCTGCATGGCTGTCGTCAGCTCGTGTCGTGAGATGTTGGGTTAAGTCCCGCAACGAGCGCAACCCTTGCCCTTAGTTGCCAGCGGTTCGGCCGGGCACTCTAAGGGGACTGCCGGCGATAAGCCGGAGGAAGGTGGGGATGACGTCAAGTCATCATGGCCCTTATGCCCAGGGCTACACACGTGCTACAATGGCTGGTACAAAGGGTCGCGAAACCGCGAGGTGGAGCTAATCCCAAAAAGCCAGTCCCAGTCCGGATCGAAGGCTGGAACTCGCCTTCGTGAAGGCGGAATCGCTAGTAATGGCGGATCAGAATGCCGCCGTGAATACGTTCCCGGGCCTTGTACACACCGCCCGTCACACCACGAAAGTCGGCTGTACCAGAAGTCGCTGGCCCAACCCCGTAAGGGGAGGGAGGCGCCCAAGGTGTGGTCGGCGATTGGGGTG

>GuaymasBasin_Dive4872_L-ASV_180

AGTGAACGCTAGGGGCGTACTTAACACATGCAAGTCGTGCGAGAAACCTACCTTTCGGGGTAGGGAGTAAAGCGGCGAACGGGTGAGTAATGCGTAGGTAACCTACCTCCGAGAGGGGAATAACCTACCGAAAGGTGGGCTAATGCCCCATAAGCTCACGATCACTACGGTGATTGTGAGAAAAGATGGCCTCTGCATGCAAGCTGTCGCTCGGAGAGGGGCTTACGTCCTATCAGCTAGTTGGTGAGGTAATGGCTCACCAAGGCTACGACGGGTAGCCGGCCTGAGAGGGTGGTCGGCCACACTGGCACTGAGACACGGGCCAGACTCCTACGGGAGGCAGCAGTGGGGAATATTGGGCAATGGGCGCAAGCCTGACCCAGTAACGCCCCGTGGGTGATGAAGGCCTTCGGGTCGTAAAGCCCTGTCAGGTGGGAAGAATATCTGATGGATGAATAGTCCATTAGATTGACGGTACCACCAAAGGAAGCACCGGCTAACTCCGTGCCAGCAGCCGCGGTAATACGGAGGGTGCAAGCGTTACTCGGAATTACTGGGCGTAAAGGGCGCGTAGGCGGGAAGGCAAGTTGAGCGTGTAAGCCTGAGGCTTAACCCCAGAATGGCGCTCAAAACTGCCTTTCTTGAGTCCCGGAGAGGCCGGCGGAATTCCCGGTGTAGGGGTGAAATCCGTAGATATCGGGAGGAACACCAGTGGCGAAGGCGGCCGGCTGGACGGGTACTGACGCTGAGGCGCGAAAGCGTGGGGAGCAAACAGGATTAGATACCCTGGTAGTCCACGCCGTAAACGATGAGCACTAGGTGTGGGGGAGATTATCTCTTCCGTGCCGCAGCTAACGCATTAAGTGCTCCGCCTGGGGAGTACGGCCGCAAGGTTAAAACTCAAAGGAATTGACGGGGGCCCGCACAAGCGGTGGAGCATGTGGTTTAATTCGATGCTACGCGAAGAACCTTACCTGGGTTTGACATCCTCGGAACCTTGCCGAAAGGTGAGGGTGCTCCTTCGGGAGAACCGGGTGACAGGTGCTGCATGGCTGTCGTCAGCTCGTGTCGTGAGATGTTGGGTTAAGTCCCGCAACGAGCGCAACCCTTGCCCTTAGTTGCCAGCGGTTTGGCCGGGCACTCTAAGGGGACTGCCGGCGATAAGCCGGAGGAAGGTGGGGATGACGTCAAGTCATCATGGCCCTTATGCCCAGGGCTACACACGTGCTACAATGGCTGGTACAAAGGGTCGCGAAACCGCAAGGTGGAGCTAATCCCAAAAAGCCAGCCCCAGTTCGGATCGAAGGCTGTAACTCGCCTTCGTGAAGGCGGAATCGCTAGTAATGGCGGATCAGAATGCCGCCGTGAATACGTTCCCGGGCCTTGTACACACCGCCCGTCACACCACGAAAGTCGGCTGTACCAGAAGTCGCTGGCCCAACCCCGCAAGGGGAGGGAGGCGCCCAAGGTGTGGTCGGCGATTGGGGTG

>GuaymasBasin_Dive4872_L-ASV_253

AGTGAACGCTAGGGGCGTACTTAACACATGCAAGTCGTGCGAGAAACCTACCTTTCGGGGTAGGGAGTAAAGCGGCGAACGGGTGAGTAATGCGTAGGTAACCTACCTCCGAGAGGGGAATAACCTACCGAAAGGTGGGCTAATGCCCCATAAGCTCACGATCACTACGGTGATTGTGAGAAAAGATGGCCTCTGCATGCAAGCTGTCGCTCGGAGAGGGGCTTACGTCCTATCAGCTAGTTGGTGAGGTAATGGCTCACCAAGGCTACGACGGGTAGCCGGCCTGAGAGGGTGGTCGGCCACACTGGCACTGAGACACGGGCCAGACTCCTACGGGAGGCAGCAGTGGGGAATATTGGGCAATGGGCGCAAGCCTGACCCAGTAACGCCCCGTGGGTGATGAAGGCCTTCGGGTCGTAAAGCCCTGTCAGGTGGGAAGAATATCTGATGGATGAATAGTCCATTAGATTGACGGTACCACCAAAGGAAGCACCGGCTAACTCCGTGCCAGCAGCCGCGGTAATACGGAGGGTGCAAGCGTTACTCGGAATTACTGGGCGTAAAGGGCGCGTAGGCGGGAAGGCAAGTTGAGCGTGTAAGCCTGAGGCTTAACCTCAGAATGGCGCTCAAAACTGCCTTTCTTGAGTCCCGGAGAGGCCGGCGGAATTCCCGGTGTAGGGGTGAAATCCGTAGATATCGGGAGGAACACCAGTGGCGAAGGCGGCCGGCTGGACGGGTACTGACGCTGAGGCGCGAAAGCGTGGGGAGCAAACAGGATTAGATACCCTGGTAGTCCACGCCGTAAACGATGAGCACTAGGTGTGGGGGAGGTTATCTCTTCCGTGCCGCAGCTAACGCATTAAGTGCTCCGCCTGGGGAGTACGGCCGCAAGGTTAAAACTCAAAGGAATTGACGGGGGCCCGCACAAGCGGTGGAGCATGTGGTTTAATTCGATGCTACGCGAAGAACCTTACCTGGGCTTGACATCCCCGGAACCCTGCCGAAAGGTGGGGGTGCCCCTTCGGGGGAACCGGGTGACAGGTGCTGCATGGCTGTCGTCAGCTCGTGTCGTGAGATGTTGGGTTAAGTCCCGCAACGAGCGCAACCCTTGCCCTTAGTTGCCAGCGGTTTAGCCGGGCACTCTAAGGGGACTGCCGGCGATAAGCCGGAGGAAGGTGGGGATGACGTCAAGTCATCATGGCCCTTATGCCCAGGGCTACACACGTGCTACAATGGCTGGTACAAAGGGTCGCGAAGCCGCGAGGTGGAGCTAATCCCAAAAAGCCAGCCCCAGTTCGGATCGAAGGCTGTAACTCGCCTTCGTGAAGGCGGAATCGCTAGTAATGGCGGATCAGAATGCCGCCGTGAATACGTTCCCGGGCCTTGTACACACCGCCCGTCACACCACGAAAGTCGGCTGTACCAGAAGTCGCTGGCCCAACTCCGCAAGGAGAGGGAGGCGCCCAAGGTGTGGTCGGCGATTGGGGTG

>GuaymasBasin_Dive4872_L-ASV_15

AGTGAACGCTAGGGGCGTACTTAACACATGCAAGTCGTGCGAGAAACCTACCTTTCGGGGTAGGGAGTAAAGCGGCGAACGGGTGAGTAATGCGTAGGTAACCTGCCTCCGAGAGGGGAATAACCTACCGAAAGGTGGGCTAATGCCCCATAAGCTCACAATCACTACGGTGATTGTGAGAAAAGATGGCCTCTGCATGCAAGCTGTCGCTCGGAGAGGGGCTTACGTCCTATCAGCTAGTTGGTGAGGTAATGGCTCACCAAGGCTACGACGGGTAGCCGGCCTGAGAGGGTGGTCGGCCACACTGGCACTGAGACACGGGCCAGACTCCTACGGGAGGCAGCAGTGGGGAATATTGGGCAATGGGCGCAAGCCTGACCCAGTAACGCCCCGTGGGTGATGAAGGCCTTCGGGTCGTAAAGCCCTGTCAGGTGGGAAGAATATCTGATGGGTAAATAGCCCATTGGATTGACGGTACCGCCAGAGGAAGCACCGGCTAACTCCGTGCCAGCAGCCGCGGTAATACGGAGGGTGCAAGCGTTACTCGGAATTACTGGGCGTAAAGGGCGCGTAGGCGGGAAGGCAAGTTGAGCGTGTAAGCCTGAGGCTCAACTTCAGAATGGCGCTCAAAACTGCCTTTCTTGAGTCCCGGAGAGGCCGGCGGAATTCCCGGTGTAGGGGTGAAATCCGTAGATATCGGGAGGAACACCGGTGGCGAAGGCGGCCGGCTGGACGGGTACTGACGCTGAGGCGCGAAAGCGTGGGGAGCAAACAGGATTAGATACCCTGGTAGTCCACGCCGTAAACGATGAGCACTAGGTGTGGGGGAGATTATCTCTTCCGTGCCGCAGCTAACGCATTAAGTGCTCCGCCTGGGGAGTACGGCCGCAAGGTTAAAACTCAAAGGAATTGACGGGGGCCCGCACAAGCGGTGGAGCATGTGGTTTAATTCGATGCTACGCGAAGAATCTTACCTGGGTTTGACATCCCCGGAACCTTGCCGAAAGGTGAGGGTGCTCCTTCGGGAGAACCGGGTGACAGGTGCTGCATGGCTGTCGTCAGCTCGTGTCGTGAGATGTTGGGTTAAGTCCCGCAACGAGCGCAACCCTTGCCCTTAGTTGCCAGCGGTTCGGCCGGGCACTCTAAGGGGACTGCCGGCGATAAGCCGGAGGAAGGTGGGGATGACGTCAAGTCATCATGGCCCTTATGCCCAGGGCTACACACGTGCTACAATGGCTGGTACAAAGGGTCGCGAAACCGCGAGGTGGAGCTAATCCCAAAAAGCCAGTCCCAGTTCGGATCGAAGGCTGTAACTCGCCTTCGTGAAGGCGGAATCGCTAGTAATGGCGGATCAGAATGCCGCCGTGAATACGTTCCCGGGCCTTGTACACACCGCCCGTCACACCACGAAAGTCGGCTGTACCAGAAGTCGCTGGCCCAACTCCGCAAGGAGAGGGAAGCGCCCAAGGTGTGGTCGGCGATTGGGGTG

>GuaymasBasin_Dive4872_L-ASV_7

AGTGAACGCTAGGGGCGTACCTAACACATGCAAGTCGTGCGAGAAACCTACCCTTCGGGGTAGGGAGTAAAGCGGCGGACGGGTGAGTAATGCGTAGGTAACCTGCCTCCGAGAGGGGAATAACCTACCGAAAGGTGGGCTAATGCCCCATAAGCTCACGGCCACTACGGTGGTTGTGAGAAAAGATGGCCTCTGCATGCAAGCTGTCGCTCGGAGAGGGGCTTACGTCCTATCAGCTAGTTGGTGAGGTAATGGCTCACCAAGGCTACGACGGGTAGCCGGCCTGAGAGGGTGGTCGGCCACACTGGCACTGAGACACGGGCCAGACTCCTACGGGAGGCAGCAGTGGGGAATATTGGGCAATGGGCGCAAGCCTGACCCAGTAACGCCCCGTGGGTGATGAAGGCCTTCGGGTCGTAAAGCCCTGTCAGGTGGGAAGAATATCTGATGGGTAAATAGCCCATTGGATTGACGGTACCACCAGAGGAAGCACCGGCTAACTCCGTGCCAGCAGCCGCGGTAATACGGAGGGTGCGAGCGTTACTCGGAATTACTGGGCGTAAAGGGCGCGTAGGCGGGAAGGCAAGTTGAGCGTGTAAGCCTGAGGCTCAACCTCAGAATGGCGCTCAAAACTGCCTTTCTTGAGTCCCGGAGAGGCCGGCGGAATTCCCGGTGTAGGGGTGAAATCCGTAGATATCGGGAGGAACACCGGTGGCGAAGGCGGCCGGCTGGACGGGTACTGACGCTGAGGCGCGAAAGCGTGGGGAGCAAACAGGATTAGATACCCTGGTAGTCCACGCCGTAAACGATGAGCACTAGGTGTGGGGGAGGTTATCTCTTCCGTGCCGAAGCTAACGCATTAAGTGCTCCGCCTGGGGAGTACGGCCGCAAGGTTAAAACTCAAAGGAATTGACGGGGGCCCGCACAAGCGGTGGAGCATGTGGTTTAATTCGATGCTACGCGAAGAACCTTACCTGGGCTTGACATCCCCGGAACCCTGCCGAAAGGTGGGGGTGCCCCTTCGGGGGAACCGGGTGACAGGTGCTGCATGGCTGTCGTCAGCTCGTGTCGTGAGATGTTGGGTTAAGTCCCGCAACGAGCGCAACCCCTGCCCTTAGTTGCCAGCGGTTCGGCCGGGCACTCTAAGGGGACTGCCGGCGATAAGCCGGAGGAAGGTGGGGATGACGTCAAGTCATCATGGCCCTTATGCCCAGGGCTACACACGTGCTACAATGGCTGGTACAAAGGGTCGCGAAACCGCGAGGTGGAGCTAATCCCAAAAAGCCAGCCCCAGTTCGGATCGAAGGCTGTAACTCGCCTTCGTGAAGGCGGAATCGCTAGTAATGGCGGATCAGAATGCCGCCGTGAATACGTTCCCGGGCCTTGTACACACCGCCCGTCACACCACGAAAGTCGGCTGTACCAGAAGTCGCTGGCCCAACCCCGCAAGGGGAGGGAGGCGCCCAAGGTGTGGTCGGCGATTGGGGTG

>GuaymasBasin_Dive4872_L-ASV_128

AGTGAACGCTAGGGGCGTACCTAACACATGCAAGTCGTGCGAGAAACCTACCCTTCGGGGTAGGGAGTAAAGCGGCGGACGGGTGAGTAATGCGTAGGTAACCTGCCTCCGAGAGGGGAATAACCTACCGAAAGGTGGGCTAATGCCCCATAAGCTCACGGCCACTACGGTGGTTGTGAGAAAAGATGGCCTCTGCATGCAAGCTGTCGCTCGGAGAGGGGCTTACGTCCTATCAGCTAGTTGGTGAGGTAATGGCTCACCAAGGCTACGACGGGTAGCCGGCCTGAGAGGGTGGTCGGCCACACTGGCACTGAGACACGGGCCAGACTCCTACGGGAGGCAGCAGTGGGGAATATTGGGCAATGGGCGCAAGCCTGACCCAGTAACGCCCCGTGGGTGATGAAGGCCTTCGGGTCGTAAAGCCCTGTCAGGTGGGAAGAATATCTGATGGGTAAATAGCCCATTGGATTGACGGTACCACCAGAGGAAGCACCGGCTAACTCCGTGCCAGCAGCCGCGGTAATACGGAGGGTGCGAGCGTTACTCGGAATTACTGGGCGTAAAGGGCGCGTAGGCGGGAAGGCAAGTTGAGCGTGTAAGCCTGAGGCTCAACCTCAGAATGGCGCTCAAAACTGCCTTTCTTGAGTCCCGGAGAGGCCGGCGGAATTCCCGGTGTAGGGGTGAAATCCGTAGATATCGGGAGGAACACCGGTGGCGAAGGCGGCCGGCTGGACGGGTACTGACGCTGAGGCGCGAAAGCGTGGGGAGCAAACAGGATTAGATACCCTGGTAGTCCACGCCGTAAACGATGAGCACTAGGTGTGGGGGAGGTTATCTCTTCCGTGCCGAAGCTAACGCATTAAGTGCTCCGCCTGGGGAGTACGGCCGCAAGGTTAAAACTCAAAGGAATTGACGGGGGCCCGCACAAGCGGTGGAGCATGTGGTTTAATTCGATGCTACGCGAAGAATCTTACCTGGGCTTGACATCCCCGGAACCCTGCCGAAAGGTGGGGGTGCCCCTTCGGGGGAACCGGGTGACAGGTGCTGCATGGCTGTCGTCAGCTCGTGTCGTGAGATGTTGGGTTAAGTCCCGCAACGAGCGCAACCCCTGCCCTTAGTTGCCAGCGGTTCGGCCGGGCACTCTAAGGGGACTGCCGGCGATAAGCCGGAGGAAGGTGGGGATGACGTCAAGTCATCATGGCCCTTATGCCCAGGGCTACACACGTGCTACAATGGCTGGTACAAAGGGTCGCGAAACCGCGAGGTGGAGCTAATCCCAAAAAGCCAGCCCCAGTTCGGATCGAAGGCTGTAACTCGCCTTCGTGAAGGCGGAATCGCTAGTAATGGCGGATCAGAATGCCGCCGTGAATACGTTCCCGGGCCTTGTACACACCGCCCGTCACACCACGAAAGTCGGCTGTACCAGAAGTCGCTGGCCCAACTCCGCAAGGAGAGGGAGGCGCCCAAGGTGTGGTCGGCGATTGGGGTG

>GuaymasBasin_Dive4872_L-ASV_3

AGTGAACGCTAGGGGCGTACCTAACACATGCAAGTCGTGCGAGAAACCTACCCTTCGGGGTAGGGAGTAAAGCGGCGGACGGGTGAGTAATGCGTAGGTAACCTGCCTCCGAGAGGGGAATAACCTACCGAAAGGTGGGCTAATGCCCCATAAGCTCACGGCCACTACGGTGGTTGTGAGAAAAGATGGCCTCTGCATGCAAGCTGTCGCTCGGAGAGGGGCTTACGTCCTATCAGCTAGTTGGTGAGGTAATGGCTCACCAAGGCTACGACGGGTAGCCGGCCTGAGAGGGTGGTCGGCCACACTGGCACTGAGACACGGGCCAGACTCCTACGGGAGGCAGCAGTGGGGAATATTGGGCAATGGGCGCAAGCCTGACCCAGTAACGCCCCGTGGGTGATGAAGGCCTTCGGGTCGTAAAGCCCTGTCAGGTGGGAAGAATATCTGATGGGTAAATAGCCCATTGGATTGACGGTACCACCAGAGGAAGCACCGGCTAACTCCGTGCCAGCAGCCGCGGTAATACGGAGGGTGCGAGCGTTACTCGGAATTACTGGGCGTAAAGGGCGCGTAGGCGGGAAGGCAAGTTGAGCGTGTAAGCCTGAGGCTCAACCTCAGAATGGCGCTCAAAACTGCCTTTCTTGAGTCCCGGAGAGGCCGGCGGAATTCCCGGTGTAGGGGTGAAATCCGTAGATATCGGGAGGAACACCGGTGGCGAAGGCGGCCGGCTGGACGGGTACTGACGCTGAGGCGCGAAAGCGTGGGGAGCAAACAGGATTAGATACCCTGGTAGTCCACGCCGTAAACGATGAGCACTAGGTGTGGGGGAGGTTATCTCTTCCGTGCCGAAGCTAACGCATTAAGTGCTCCGCCTGGGGAGTACGGCCGCAAGGTTAAAACTCAAAGGAATTGACGGGGGCCCGCACAAGCGGTGGAGCATGTGGTTTAATTCGATGCTACGCGAAGAATCTTACCTGGGCTTGACATCCCCGGAACCCTGCCGAAAGGTGGGGGTGCCCCTTCGGGGGAACCGGGTGACAGGTGCTGCATGGCTGTCGTCAGCTCGTGTCGTGAGATGTTGGGTTAAGTCCCGCAACGAGCGCAACCCCTGCCCTTAGTTGCCAGCGGTTCGGCCGGGCACTCTAAGGGGACTGCCGGCGATAAGCCGGAGGAAGGTGGGGATGACGTCAAGTCATCATGGCCCTTATGCCCAGGGCTACACACGTGCTACAATGGCTGGTACAAAGGGTCGCGAAACCGCGAGGTGGAGCTAATCCCAAAAAGCCAGCCCCAGTTCGGATCGAAGGCTGTAACTCGCCTTCGTGAAGGCGGAATCGCTAGTAATGGCGGATCAGAATGCCGCCGTGAATACGTTCCCGGGCCTTGTACACACCGCCCGTCACACCACGAAAGTCGGCTGTACCAGAAGTCGCTGGCCCAACCCCGCAAGGGGAGGGAGGCGCCCAAGGTGTGGTCGGCGATTGGGGTG

>GuaymasBasin_Dive4872_L-ASV_130

AACGAACGCTGGCGGCGTGCCTAACACATGCAAGTCGAACGAGAAACTCCGGGCTTGCTCGGAGAAGTACAGTGGCGCACGGGTGAGTAACGCGTGGGTAACCTACCCTTGAATCTGGGATAACCCCGCGAAAGCGGGACTAATACCGGATATGGTTCCATAAGCCTCGGCTCTTGGAATTAAAGGTGACCTCTTCATGAAAGTTGCCGTTCAAGGACGGGCCCGCGTACCATTAGCTTGTTGGTAGGGTAATGGCCTACCAAGGCTACGATGGTTAGCTGGTCTGAGAGGATGATCAGTCACACTGGAACTGGAACACGGTCCAGACTCCTACGGGAGGCAGCAGTGAGGAATTTTGCGCAATGGGGGAAACCCTGACGCAGCAACGCCGCGTGAGCGAAGAAGGCCTTCGGGTCGTAAAGCTCTGTCAAGTGGGAAAAAAGCCTTTCGGTTAATACCCGGGAGGTCTGATGGTACCACTGGAGGAAGCACCGGCTAACTCCGTGCCAGCAGCCGCGGTAATACGGAGGGTGCAAGCGTTGTTCGGAATTACTGGGCGTAAAGCGCGTGCAGGCGGTTTGGTAAGTCAGATGTGAAAGCCCTGGGCTCAACCCGGGAAGTGCATTTGAAACTGCCTTTCTTGAGTATGGGAGAGGAGAGTGGAATTCCCAGTGTAGAGGTGAAATTCGTAGATATTGGGAGGAACACCCGTGGCGAAGGCGACTCTCTGGACCAATACTGACGCTGAGACGCGAAAGCGTGGGGAGCAAACAGGATTAGATACCCTGGTAGTCCACGCCGTAAACGATGAGAACTAGGTGTAGCGGGTATTGACCCCTGCTGTGCCGAAGTTAACGCATTAAGTTCTCCGCCTGGGGAGTACGGCCGCAAGGCTAAAACTCAAAGGAATTGACGGGGGCCCGCACAAGCGGTGGAGCATGTGGTTTAATTCGACGCAACGCGAAGAACCTTACCTGGGTTTGACATCCTTTGACCGTCTGGGAAACCAGATTTTTCCGACTCTGTCGGAACAGAGTGACAGGTGCTGCATGGCTGTCGTCAGCTCGTGTCGTGAGATGTTGGGTTAAGTCCCGCAACGAGCGTAACCCTTGTCTTTAGTTGCCACTATTAAGTTAGGCACTCTAAAGAGACTGCCTCGGTTAACGGGGAGGAAGGTGGGGATGACGTCAAGTCCTCATGGCCTTTATACTCAGGGCTACACACGTGCTACAATGGGCTGTACAAAGGGTTGCCACTCCGCGAGGAGGAGCTAATCCCAAAAAGCAGTCCTCAGTTCGGATTGGAGTCTGCAACTCGACTCCATGAAGGTGGAATCGCTAGTAATCGTGGATCAGCATGCCACGGTGAATACGTTCCCGGGCCTTGTACACACCGCCCGTCACACCACGAAAGTCGACTGTACCAGAAGTTGCTGGGCCAACCCCATTTGGGGAGGCAGGTACCTAAGGTACGGCCGGTAATTGGGGTG

>GuaymasBasin_Dive4872_L-ASV_138

AACGAACGCTGGCGGCGTGCCTAACACATGCAAGTCGAACGAGAAACTCCGGGCTTGCTCGGAGGAGTAAAGTGGCGCACGGGTGAGTAACGCGTGGGTAACCCACCCTTGAAACCGGGATAACCCCGCGAAAGCGGGACTAATACCGGATACGGTTTTTTAGGCTCCGGCCTTTGGAATTAAAGGTGACCTCTTCATGAAAGTTGCCGTTCAGGGACGGGCTCGCGTACCATTAGCTTGTTGGTAGGGTAATGGCCTACCAAGGCTACGATGGTTAGCTGGTCTGAGAGGATGATCAGCCACACTGGAACTGGAACACGGTCCAGACTCCTACGGGAGGCAGCAGTGAGGAATTTTGCGCAATGGGGGAAACCCTGACGCAGCAACGCCGCGTGAGCGAAGAAGGCCTTCGGGTCGTAAAGCTCTGTCAAGTGGGAAAAAAGCCTTTCGGTTAATACCCGGAAGGTCTGATGGTACCACTGGAGGAAGCACCGGCTAACTCCGTGCCAGCAGCCGCGGTAATACGGAGGGTGCAAGCGTTGTTCGGAATTACTGGGCGTAAAGCGCGTGCAGGCGGTTTGGTAAGTCAGATGTGAAAGCCCTGGGCTCAACCCGGGAAGTGCATTTGAAACTGCCTTTCTTGAGTATGGGAGAGGAGAGTGGAATTCCCAGTGTAGAGGTGAAATTCGTAGATATTGGGAGGAACACCCGTGGCGAAGGCGACTCTCTGGACCAATACTGACGCTGAGACGCGAAAGCGTGGGGAGCAAACAGGATTAGATACCCTGGTAGTCCACGCCGTAAACGATGAGAACTAGGTGTAGCGGGTATTGACCCCTGCTGTGCCGAAGTTAACGCATTAAGTTCTCCGCCTGGGGAGTACGGCCGCAAGGCTAAAACTCAAAGGAATTGACGGGGGCCCGCACAAGCGGTGGAGCATGTGGTTTAATTCGACGCAACGCGAAGAACCTTACCTGGGTTTGACATCCTCTGACCGCCTTGGAAACAAGGTTTTTCCGGTTCTGCCGGAACAGAGTGACAGGTGCTGCATGGCTGTCGTCAGCTCGTGTCGTGAGATGTTGGGTTAAGTCCCGCAACGAGCGTAACCCTTGTCTTTAGTTGCCATTATTAAGTTAGGCACTCTAAAGAGACTGCCTCGGTTAACGGGGAGGAAGGTGGGGATGACGTCAAGTCCTCATGGCCTTTATACCCAGGGCTACACACGTGCTACAATGGGCTGTACAAAGGGTTGCCACTCCGCGAGGAGGCGCTAATCCCAAAAAGCAGTCCTCAGTTCGGATTGGAGTCTGCAACTCGACTCCATGAAGGTGGAATCGCTAGTAATCGTGGATCAGCACGCCACGGTGAATACGTTCCCGGGCCTTGTACACACCGCCCGTCACACCACGAAAGTCGACTGTACCAGAAGTTGCTGGGCCAACCACGCCTTGGCGTGGAGGCAGGTACCTAAGGTACGGCCGGTAATTGGGGTG

>GuaymasBasin_Dive4872_L-ASV_93

AACGAACGCTGGCGGCGTGCCTAACACATGCAAGTCGAACGAGAAACTCCGGGCTTGCTCGGAGGAGTAAAGTGGCGCACGGGTGAGTAACGCGTGGGTAACCTGCCCCTGAAACTGGGATAACCCGCCGAAAGGCGGACTAATACCGGATATTGTCCTTTTAGCTGTGGCTTTTGGGATGAAAGGCAACCTCTTCATGAAAGTTGTCGTTCGGGGAGGGGCTCGCGTACCATTAGCTTGTTGGTGGGGTAATGGCCTACCAAGGCTACGATGGTTAGCTGGTCTGAGAGGATGATCAGCCACACTGGAACTGAAACACGGTCCAGACTCCTACGGGAGGCAGCAGTGAGGAATTTTGCGCAATGGGGGAAACCCTGACGCAGCAACGCCGCGTGAGTGAAGAAGGCCTTTGGGTCGTAAAGCTCTGTCAAGTGGGAAAAAACGCTTTCGATTAATAGTCGAAAGAATTGATGGTACCACTGGAGGAAGCACCGGCTAACTCCGTGCCAGCAGCCGCGGTAATACGGAGGGTGCAAGCGTTGTTCGGAATTACTGGGCGTGAAGAGCGTGTAGGCGGTTTGGCAAGTCAGATGTGAAAGCCCTGGGCTTAACCCAGGAAGTGCATTTGAAACTGCCAGACTAGAGTATGGGAGAGGAGAGTGGAATTCCCAGTGTAGAGGTGAAATTCGTAGATATTGGGAGGAACACCCGTGGCGAAGGCGACTCTCTGGACCAATACTGACGCTGAGACGCGAAAGCGTGGGGAGCAAACAGGATTAGATACCCTGGTAGTCCACGCCGTAAACGATGAGAACTAGGTGTAGCGGGTATTGACCCCTGCTGTGCCGAAGTTAACGCATTAAGTTCTCCGCCTGGGGAGTACGGTCGCAAGGCTAAAACTCAAAGGAATTGACGGGGGCCCGCACAAGCGGTGGAGCATGTGGTTTAATTCGACGCAACGCGAAGAACCTTACCTGGGTTTGACATCCTCTGACCACCACGGAAACGTGGTTTTCCTGACTTGTCAGGACAGAGTGACAGGTGCTGCATGGCTGTCGTCAGCTCGTGTCGTGAGATGTTGGGTTAAGTCCCGCAACGAGCGCAACCCTTGTCTTTAGTTGCTAGTATTAATTTACGCACTCTAAAGAGACTGCCTCGGTTAACGGGGAGGAAGGTGGGGATGACGTCAAGTCCTCATGGCCTTTATATCCAGGGCTACACACGTGCTACAATGGGCTGTACAAAGGGTCGCTAACTCGCGAGAGTACGCTAATCCCAAAAAACAGTTCTCAGTTCGGATTGGAGTCTGCAACTCGACTCCATGAAGTTGGAATCGCTAGTAATCGTGGATCAGCATGCCACGGTGAATACGTTCCCGGGCCTTGTACACACCGCCCGTCACACCACGAAAGTCGACTGTACCAGAAGTTGCTGGGCCAACCCCGCTTGCGGGGAGGTAGGTACCTAAGGTACGGCCGGTAATTGGGGTG

>GuaymasBasin_Dive4872_L-ASV_27

AACGAACGCTGGCGGCGTGCCTAACACATGCAAGTCGAACGAGAAATCGGGCACTCAGGCAACTGAGTGCCGGAGAGTAAAGTGGCGCACGGGTGAGTAACGCGTGGGTAACCCGCCCTTGAATTTGGGATAACCTCGTGAAAACGGGACTAATACCGAATATTGTCCTGAGAACCATGGTTCTTGGGATGAAAGGTGACCTCTTCATGAAAGTTGCCGTTCAGGGAAGGGCCCGCGTACCATTAGCTTGTTGGTGGGGTAATGGCCTACCAAGGCGACGATGGTTAGCTGGTCTGAGAGGATGATCAGCCACACTGGAACTGAAACACGGTCCAGACTCCTACGGGAGGCAGCAGTGAGGAATTTTGCGCAATGGGGGAAACCCTGACGCAGCAACGCCGCGTGAGTGAAGAAGGCCTTCGGGTCGTAAAGCTCTGTCAAGTGGGAAAAAATGTCCCGGGTTAATAGCCCGGGAAATTGATGGTACCACTGGAGGAAGCACCGGCTAACTCCGTGCCAGCAGCCGCGGTAATACGGAGGGTGCAAGCGTTGTTCGGAATTACTGGGCGTAAAGAGCGTGTAGGCGGTTTGGCAAGTCAGATGTGAAAGCCCCGGGCTCAACCCGGGAAGTGCATTTGAAACTGCTATACTTGAGTATGGGAGAGGAGAGTGGAATTCCCAGTGTAGAGGTGAAATTCGTAGATATTGGGAGGAACACCCGTGGCGAAGGCGACTCTCTGGACCAATACTGACGCTGAGACGCGAAAGCGTGGGGAGCAAACAGGATTAGATACCCTGGTAGTCCACGCTGTAAACGATGAGAACTAGGTGTAGCGGGTATTGACCCCTGCTGTGCCGAAGTTAACGCATTAAGTTCTCCGCCTGGGGAGTACGGCCGCAAGGCTAAAACTCAAAGGAATTGACGGGGGCCCGCACAAGCGGTGGAGCATGTGGTTTAATTCGACGCAACGCGAAGAACCTTACCTGGGTTTGACATCCTCTGACCGTCTGGGAAACCAGATTTTTCCGGCTCTGCCGGAACAGAGTGACAGGTGCTGCATGGCTGTCGTCAGCTCGTGTCGTGAGATGTTGGGTTAAGTCCCGCAACGAGCGCAACCCTTATTTTTAGTTGCCAGTATTAAGTTAGGCACTCTAAAGAGACTGCCTCGGTTAACGAGGAGGAAGGTGGGGATGACGTCAAGTCCTCATGGCCTTTATACCCAGGGCTACACACGTGCTACAATGGGCTGTACAAAGGGTCGCCAATCCGCGAGGATGCGCTAATCCCAAAAAGCAGCTCTCAGTTCGGATTGGAGTCTGCAACTCGACTCCATGAAGGTGGAATCGCTAGTAATCGTGGATCAGCATGCCACGGTGAATACGTTCCCGGGCCTTGTACACACCGCCCGTCACACCACGAAAGTCGACTGTACCAGAAGTTGCTGGGCTAACTCCCGATTTATCGGGGGAGGCAGGTACCTAAGGTACGGCCGGTAATTGGGGTG

>GuaymasBasin_Dive4872_L-ASV_183

AATGAACGCTGGCGGCGTGCCTAACACATGCAAGTCGAACGAGAAATTTCCGCTTCGGCGGAAAGAGTAAAGTGGCGCACGGGTGAGTAACGCGTGGGTAACCTGCCTCTGAATCTGGGATAACCCCGTGAAAACGGGACTAATACCGGATAATGTCCCTGTGGCTGTGGCCATTGGGATGAAAGGTGACCTCTTTATGAAAGTTGCCGTTCGGAGAGGGGCTCGCGTACCATTAGCTTGTTGGTGGGGTAATGGCCTACCAAGGCTACGATGGTTAGCTGGTCTGAGAGGATGATCAGCCACACTGGAACTGGAACACGGTCCAGACTCCTACGGGAGGCAGCAGTGAGGAATTTTGCGCAATGGGGGAAACCCTGACGCAGCAACGCCGCGTGAGTGATGAAGATTTTCGGATCGTAAAGCTCTGTCAAGCGGGAAAAAATACTTTCGGTCAATAGCCGGAAGGACTGATGGTACCGCTGGAGGAAGCACCGGCTAACTCCGTGCCAGCAGCCGCGGTAATACGGAGGGTGCAAGCGTTGTTCGGATTTACTGGGCGTAAAGAGCGTGTAGGCGGTTTAGCAAGTCAGATGTGAAAGCCCTGGGCTCAACTCAGGAAGTGCATTTGAAACTGCTATACTAGAGTATGGGAGAGGAGATTGGAATTCCCAGTGTAGAGGTGAAATTCGTAGATATTGGGAGGAACACCGGTGGCGAAGGCGACTCTCTGGACCGATACTGACGCTGAGACGCGAAAGCGTGGGGAGCAAACAGGATTAGATACCCTGGTAGTCCACGCCGTAAACGATGAGAACTAGGTGTAGCGGGTATTGACCCCTGCTGTGCCGCAGCTAACGCATTAAGTTCTCCGCCTGGGGAGTACGGCCGCAAGGCTAAAACTCAAAGGAATTGACGGGGGCCCGCACAAGCGGTGGAGCATGTGGTTTAATTCGACGCAACGCGAAGAACCTTACCTGGATTTGACATCCTCTTGCCACCCTGGAAACAGGGTTTTTCCGACTTGTCGGAAGAGAGTGACAGGTGCTGCATGGCTGTCGTCAGCTCGTGTCGTGAGATGTTGGGTTAAGTCCCGCAACGAGCGCAACCCTTGTCTCTAGTTGCCAGTATTAAGTTAGGGACTCTAGAGAGACTGCCTCGGTTAACGGGGAGGAAGGTGGGGATGACGTCAAGTCCTCATGGCCTTTATATCCAGGGCTACACACGTGCTACAATGGGCTGTACAAAGGGTAGCTAACTCGCGAGAGTATGCCAATCCCAAAAAACAGTTCTCAGTTCGGATTGGAGTCTGCAACTCGACTCCATGAAGTTGGAATCGCTAGTAATCGTGGATCAGCATGCCACGGTGAATACGTTCCCGGGCCTTGTACACACCGCCCGTCACACCACGAAAGTCGACTGTACCAGAAGTTGCCGGGCTAACCCTTCGGGGAGGCAGGTACCTAAGGTACGGCTGGTAATTGGGGTG

>GuaymasBasin_Dive4872_L-ASV_150

AATGAACGCTGGCGGCGTGCCTAACACATGCAAGTCGAACGAGAAATCAGGCACTCAGGCAACTGAGTGCCGGAGAGTAAAGTGGCGCACGGGTGAGTAACGCGTGGGTAACCTGCCCTTGAAACCGGGATAACATCGCGAAAGCGTTGCTAATACCGGATACTGTCCCAAGAACTATGGTTCCTGGGATGAAAGGTGACCTCTCCGTGGAAGTTGCCGTTCAAGGAGGGGCCCGCGTACCATTAGCTTGTTGGTAGGGTAATGGCCTACCAAGGCGACGATGGTTAGCTGGTCTGAGAGGATGATCAGCCACACTGGAACTGAAACACGGTCCAGACTCCTACGGGAGGCAGCAGTGAGGAATTTTGCGCAATGGGGGAAACCCTGACGCAGCAACGCCGCGTGAGTGAAGAAGGCCTTCGGGTCGTAAAGCTCTGTCAAGGGGGAAAAAAGTCCTGCAGTGAATAGTTGTAGGATCTGATGGTACCTCTAGAGGAAGCACCGGCTAACTCCGTGCCAGCAGCCGCGGTAATACGGAGGGTGCGAGCGTTGTTCGGAATTACTGGGCGTAAAGAGCGTGTAGGCGGTTTGGCAAGTCAGATGTGAAAGCCTTCCGCTCAACGGAAGAAGTGCATTTGAAACTGCCATTCTAGAGTATGGGAGAGGAGAGTGGAATTCCCGGTGTAGAGGTGAAATTCGTAGATATCGGGAGGAACACCTGTGGCGAAGGCGACTCTCTGGACCAATACTGACGCTGAGACGCGAAAGCGTGGGGAGCAAACAGGATTAGATACCCTGGTAGTCCACGCCGTAAACGATGAGCACTAGGTGTAGCGGGTATTGACCCCTGCTGTGCCGTAGCTAACGCATTAAGTGCTCCGCCTGGGGAGTACGGCCGCAAGGCTAAAACTCAAAGGAATTGACGGGGGCCCGCACAAGCGGTGGAGCATGTGGTTTAATTCGACGCAACGCGAAGAACCTTACCTGGGCTTGACATCCTCTGACAGCCCCGGAAACGGGGTCTTTCCTGCTTGCAGGAACAGAGTGACAGGTGCTGCATGGCTGTCGTCAGCTCGTGTCGTGAGATGTTGGGTTAAGTCCCGCAACGAGCGCAACCCCTGTGTTTAGTTGCCAGTATTGAGTTAGGCACTCTAAACAGACTGCCTCGGTTAACGAGGAGGAAGGTGGGGATGACGTCAAGTCCTCATGGCCTTTATGCCCAGGGCTACACACGTGCTACAATGGGCTGTACAAAGGGTTGCTACTCCGCGAGGAGGCGCTAATCTCAAAAAGCAGCTCTCAGTTCGGATTGGAGTCTGCAACTCGACTCCATGAAGTCGGAATCGCTAGTAATCGTGGATCAGCATGCCACGGTGAATACGTTCCCGGGCCTTGTACACACCGCCCGTCACACCACGAAAGCTGGCTGTACCAGAAGTTGCTGGGCTAACCCCGCACCTACTTTGCCAATAACTTGATTAGAGGTATTATCTCGAGGGGTCGTTGGTAAAGTAGGTGCGGGGAGGCAGGTACCTAAGGTACGGTTGGTAATTGGGGTG

>GuaymasBasin_Dive4872_L-ASV_40

AATGAACGCTGGCGGCGTGCCTAACACATGCAAGTCGAACGAGAAACTCCGAGCTTGCTCGGAGGAGTAAAGTGGCGCACGGGTGAGTAACGCGTGGGTAACCTGCCCTAGAATCTGGGATAACTTCGCGAAAGCGTTGCTAATACCGGATATCGTCCCGAGAACCGTGGTTTTTGGGATGAAAGATGACCTCTTCATGAAAGTTGTCGTTCAAGGAGGGGCCCGCGTACCATTAGCTTGTTGGTGGGGTAATGGCCTACCAAGGTGACGATGGTTAGCTGGTCTGAGAGGATGATCAGCCACACTGGAACTGAAACACGGTCCAGACTCCTACGGGAGGCAGCAGTGAGGAATTTTGCGCAATGGGGGAAACCCTGACGCAGCAACGCCGCGTGAGTGAAGAAGGCTTTCGGGTCGTAAAGCTCTGTCAAGTGGGAAAAAACGCTTCCGGTCAATAGCCGGAAGAATTGATGGTACCACTGGAGGAAGCACCGGCTAACTCCGTGCCAGCAGCCGCGGTAATACGGAGGGTGCAAGCGTTGTTCGGAATTATTGGGCGTAAAGAGCGTGTAGGCGGTTTGGTAAGTCAGATGTGAAAGCCCTGGGCTCAACCCAGGAAGGGCATTTGATACTGCCTGACTTGAGTATGGGAGAGGAGAGTGGAATTCCCAGTGTAGAGGTGAAATTCGTAGATATTGGGAGGAACACCCGTGGCGAAGGCGACTCTCTGGACCAATACTGACGCTGAGACGCGAAAGCGTGGGGAGCAAACAGGATTAGATACCCTGGTAGTCCACGCCGTAAACGATGAGAACTAGGTGTAGCGGGTATTGACCCCTGCTGTGCCGAAGTTAACGCATTAAGTTCTCCGCCTGGGGAGTACGGCCGCAAGGCTAAAACTCAAAGGAATTGACGGGGACCCGCACAAGCGGTGGAGCATGTGGTTTAATTCGACGCAACGCGAAGAACCTTACCTGGATTTGACATCCTCTGACAGCCCCAGAAACGGGGTTTATCCGACTTGTCGGAACAGAGTGACAGGTGCTGCATGGCTGTCGTCAGCTCGTGTCGTGAGATGTTGGGTTAAGTCCCGCAACGAGCGCAACCCTTGTTTTTAGTTGCCAGTATTAAGTTAGGCACTCTAGAGATACTGCCTCGGTTAACGGGGAGGAAGGTGGGGATGACGTCAAGTCCTCATGGCCTTTATATCCAGGGCTACACACGTGCTACAATGGGCTGTACAAAGGGTCGCAACTCCGCGAGGAGAAGCTAATCTCAAAAAGCAGTTCTCAGTTCGGATTGGAGTCTGCAACTCGACTCCATGAAGTTGGAATCGCTAGTAATCGTGGATCAGCATGCCACGGTGAATACGTTCCCGGGTCTTGTACACACCGCCCGTCACACCACGAAAGTCGGCTGTACCAGAAGTTGCTGGGCTAACCCTTCGGGGAGGCAGGTACCTAAGGTACGGTCGGTAATTGGGGTG

>GuaymasBasin_Dive4872_L-ASV_104

AATGAACGCTGGCGGCGTGCCTAACACATGCAAGTCGAACGAGAAACTTCGAGCTTGCTCGAAGGAGTAAAGTGGCGCACGGGTGAGTAACGCGTGGGTAACCTGCCCTTGAATTTGGGATAACCCCGTGAAAACGGGGCTAATACCGGATATTATCACGAGAACTATGGTTTTCGGGATAAAAGGTGACCTCTTCATGAAAGTTGCCGTTCATAGAGGGGCCCGCGTACCATTAGCTTGTTGGCAGGGTAATGGCCTACCAAGGCTACGATGGTTAGCTGGTCTGAGAGGATGATCAGCCACACTGGAACTGAAACACGGTCCAGACTCCTACGGGAGGCAGCAGTGAGGAATTTTGCGCAATGGGGGAAACCCTGACGCAGCAACGCCGCGTGAGTGAAGAAGGCTTTCGGGTCGTAAAGCTCTGTCAAAGGGGAAAAAATCCCTGTGATAAATAGTCACAGGGGCTGATGGTACCCTTAGAGGAAGCACCGGCTAACTCCGTGCCAGCAGCCGCGGTAATACGGAGGGTGCAAGCGTTGTTCGGATTTACTGGGCGTAAAGAGCGTGTAGGCGGTTTGGTAAGTCAGATGTGAAAGTCTTCCGCTCAACGGGAGAAGTGCATTTGAAACTGCCATTCTAGAGTATGGGAGAGGAGAGTGGAATTCCCAGTGGAGAGGTGAAATTCGTAGATATTGGGAGGAACACCCGTGGCGAAGGCGGCTCTCTGGACCAATACTGACGCTGAGACGCGAAAGCGTGGGGAGCAAACAGGATTAGATACCCTGGTAGTCCACGCCGTAAACGATGAGAACTAGGTGTAGCGGGTATTGACCCCTGCTGTGCCGTAGCTAACGCATTAAGTTCTCCGCCTGGGGAGTACGGCCGCAAGGCTAAAACTCAAAGGAATTGACGGGGGCCCGCACAAGCGGTGGAGCATGTGGTTTAATTCGACGCAACGCGAAGAACCTTACCTGGATTTGACATCCTCTGACAGTCCCGGAAACGGGATCTTTTCGACTTGTCGAAACAGAGTGACAGGTGCTGCATGGCTGTCGTCAGCTCGTGTCGTGAGATGTTGGGTTAAGTCCCGCAACGAGCGCAACCCTTGTCTTTAGTTGCCAGTATTAAGTTAGGAACTCTAAAGATACCGCCTCGGTTAACGGGGAGGAAGGTGGGGATGACGTCAAGTCCTCATGGCCTTTATGTCCAGGGCTACACACGTGCTACAATGGGCTGTACAAAGGGCAGCAAATCCGCGAGGAGGCGCTAATCTCAAAAAGCAGCTCTCAGTTCGGATTGGAGTCTGCAACTCGACTCCATGAAGGTGGAATCGCTAGTAATCGTGGATCAGCATGCCACGGTGAATACGTTCCCGGGCCTTGTACACACCGCCCGTCACACCACGAAAGTCGACTGTACCAGAAGTTGCTGGGCTAACCCCCGATTTATCGGGGGGGGCAGGCACCTAAGGTACGGCCGGTAATTGGGGTG

>GuaymasBasin_Dive4872_L-ASV_56

AATGAACGCTGGGGGCGCGCTTAACACATGCAAGTCGAACGAGAAATCTCGAGCTTGCTCGAGAAAGTAAAGTGGCGCACGGGTGAGTAACGCGTGGGTAACCTGTCCTTAAATTTGGGATAATCCTCCGAAAGGTGGCCTAATACCAAATACAGTCCTGAGAACCAAGGTTCTTTGGATGAAAGCTGACCTCTTCATGAAAGTTGGCGTTTAAGGAGGGGCTCGCGTACCATTAGCTTGTTGGTGGGGTAATGGCTTACCAAGGCTACGATGGTTAGCTGGTCTGAGAAGATGGCCAGCCACACTGGAACTGAAACACGGTCCAGACTCCTACGGGAGGCAGCAGTGAGGAATCTTGCGCAATGGGGGAAACCCTGACGCAGTGACGCCGTGTGAGTGAAGAAGGCCTTCGGGTCGTAAAGCTCTGTCAAGTGGGAATAAAGTCCTGCGGTGAATACCCGAGGGATTTGAAGGTACCACTAAAGGAAGCACCGGCTAACTCCGTGCCAGCAGCCGCGGTAATACGGAGGGTGCAAGCGTTATTCGGATTTATTGGGCGTAAAGAGCGTGTAGGCGGTTGGGATAGTCAGACGTGAAAGCCTTCTGCTCAACAGAAGAAGTACGTCTGAAACTGCCCAACTTGAGTACGAGAGAGGAAAGTGGAATTCCCAGTGTAGAGGTGAAATTCGTAGATATTGGGAGGAACACCTGTGGCGAAGGCGACTTTCTGGATCGATACTGACGCTGAGACGCGAAAGCGTGGGTAGCAAACAGGATTAGATACCCTGGTAGTCCACGCTGTAAACGATGGGAACTAGGTGTAGCGGGTATTGATCCCTGCTGTGCCGAAGCTAACGCATTAAGTTCCCCGCCTGGGGAGTACGGTCGCAAGGCTAAAACTCAAAGGAATTGACGGGGGCCCGCACAAGCGGTGGAGCATGTGGTTTAATTCGATGCAACGCGAAGAACCTTACCTGGGTTTGACATCCTTTGACAGTCCCTGAAAGGGGACCTTACCGATTCTTCGGAACAAAGTGACAGGTGCTGCATGGCTGTCGTCAGCTCGTGTCGTGAGATGTTGGGTTAAGTCCCGCAACGAGCGCAACCCTTGTCTTTAGTTGCCACTATTAAGTTAGGCACTCTAAAGATACTGCCCCGGTTAACGGGGAGGAAGGTGGGGATGACGTCAAGTCCTCATGGCCTTTATACCCAGGGCTACACACGTGCTACAATGGGCTGTACAAAGGGCTGCGACTCTGTAAGGAGAAGCTAATCCCAAAAAGTAGTCCTAAGTTCGGATTGGAGTCTGCAACTCGACTCCATGAAGCTGGAATCGCTAGTAATCGTGGATCAGCATGCCACGGTGAATACGTTCCCGGGCCTTGTACACACCGCCCGTCACACCACGAAAGCTGGTTGTACCAGAAGTTGCTGGGCTAACCCCTGAGCTTGTTGAAGGGGAGGCAGGCACCTAAGGTATGGCTGGTGATTGGGGTG

>GuaymasBasin_Dive4872_L-ASV_201

AATGAACGCTGGCGGCGCGCCTAACACATGCAAGTCGAACGAGAAACTCCGAGCTTGCTCGGAGGAGTAAAGTGGCGCACGGGTGAGTAACGCGTGGGTAACCTGCCCTTAAATCCGGGATAATCCAACGAAAGATGGCCTAATACCGGATGTTGTCCTGAGAACCACGGTTTTCGGGATGAAAGCTGACCTCTTCACGAAAGTTGGCGTTTAGGGAGGGGCTCGCGTACCATTAGCTTGTTGGTAGGGTAATGGCCTACCAAGGCTACGATGGTTAGCTGGTCTGAGAAGATGATCAGCCACACTGGAACTGGAACACGGTCCAGACTCCTACGGGAGGCAGCAGTGAGGAATCTTGCGCAATGGGGGAAACCCTGACGCAGCGACGCCGCGTGAGTGAAGAAGGCCTTCGGGTCGTAAAGCTCTGTCAAGTGGGAATAAAATCCAGCGGTGAATAGCCGTAAGGATCTGAAGGTACCACTAGAGGAAGCACCGGCTAACTCCGTGCCAGCAGCCGCGGTAATACGGAGGGTGCGAGCGTTATTCGGATTTATTGGGCGTAAAGAGCGTGTAGGTGGTTGGGTAAGTCAGATGTGAAAGCCTTTTGCTCAACAAAAGAAGTGCATCTGAAACTGCTCAACTTGAGTACGAGAGAGGAGAGTGGAATTCCCAGTGTAGAGGTGAAATTCGTAGATATTGGGAGGAACACCTGTGGCGAAGGCGACTCTCTGGATCGATACTGACGCTGAGACGCGAAAGCGTGGGGAGCAAACAGGATTAGATACCCTGGTAGTCCACGCCGTAAACGATGGGAACTAGGTGTAGCGGGTATTGATCCCTGCTGTGCCGAAGCTAACGCATTAAGTTCCCCGCCTGGGGAGTACGGTCGCAAGGCTAAAACTCAAAGGAATTGACGGGGGCCCGCACAAGCGGTGGAGCATGTGGTTTAATTCGATGCAACGCGAAGAACCTTACCTGGGTTTGACATCCTCTGACAGTCCCTGAAAGGGGATCTTCTCGATTTTTCGGGACAGAGTGACAGGTGCTGCATGGCTGTCGTCAGCTCGTGTCGTGAGATGTTGGGTTAAGTCCCGCAACGAGCGCAACCCTTGTCTTTAGTTGCCAGTATTAAGTTAGGCACTCTAAAGATACTGCCACGGTTAACGTGGAGGAAGGTGGGGACGACGTCAAGTCCTCATGGCCTTTATACCCAGGGCTACACACGTGCTACAATGGGCTGTACAAAGGGTTGCAACGCCGTGAGGCGAAGCTAATCCCAAAAAATAGTCCTCAGTTCGGATTGGAGTCTGCAACTCGACTCCATGAAGTTGGAATCGCTAGTAATCGTGGATCAGCATGCCACGGTGAATACGTTCCCGGGCCTTGTACACACCGCCCGTCACACCACGAAAGTTGGCTGTACCAGAAGTTGTTGAGCTAACCCCTGAGCTTGTCGAAGGGGAGGCAGGCACCTAAGGTATGGTTGATAATTGGGGTG

>GuaymasBasin_Dive4872_L-ASV_190

AACGAACGCTGGCGGCGTGCCTAACACATGCAAGTCGAACGAGAAACCCCGGGCTTGCTCGGGGGAGTAAAGTGGCGAACGGGTGAGTAACGCGTGGGTAACCTACCCCTGAATCTGGGATAACTTCGCGAAAGCGTTGCTAATACCGGATAATGTCCTAGGAACTGTGGTTCCTGGGATGAAAGGTGACCTCTCCATGCAAGTTGCCGTTCAGGGATGGGCCCGCGTCCCATTAGCTTGTTGGTAGGGTAATGGCCTACCAAGGCTACGATGGGTAGCTGGTCTGAGAGGATGATCAGCCACACTGGCACTGAAACACGGGCCAGACTCCTACGGGAGGCAGCAGTGAGGAATTTTGCGCAATGGGGGAAACCCTGACGCAGCAACGCCGCGTGAGTGAAGAAGGCCTTCGGGTTGTAAAGCTCTGTCAGGTGGGAAAAAAGTCCTACTGTGAATAGCGGTAGGATCTGATGGTACCACCAGAGGAAGCACCGGCTAACTCCGTGCCAGCAGCCGCGGTAATACGGAGGGTGCAAGCGTTGTTCGGAATTACTGGGCGTAAAGCGGGTGTAGGCGGTTTGGCAAGTCAGGTGTGAAAGCCCGGGGCTCAACCCCGGAAGTGCACTTGATACTGCCATTCTTGAGTACGGGAGAGGAGAGTGGAATTCCCGGTGTAGAGGTGAAATTCGTAGATATCGGGAGGAACACCCGTGGCGAAGGCGACTCTCTGGACCGATACTGACGCTGAGACCCGAAAGCGTGGGGAGCAAACAGGATTAGATACCCTGGTAGTCCACGCCGTAAACGATGAGAACTAGGTGTAGCGGGTATTGACCCCTGCTGTGCCGAAGCTAACGCATTAAGTTCTCCGCCTGGGGAGTACGGCCGCAAGGTTAAAACTCAAAGGAATTGACGGGGGCCCGCACAAGCGGTGGAGCATGTGGTTTAATTCGACGCAACGCGAAGAACCTTACCTGGGCTTGACATCCCCTGACAGCCCTGGAAACAGGGTCTTCCCGGTTCTGCCGGGACAGGGTGACAGGTGCTGCATGGCTGTCGTCAGCTCGTGTCGTGAGATGTTGGGTTAAGTCCCGCAACGAGCGCAACCCTTGTCTCTAGTTGCCAGTATTAGGTTAGGCACTCTAGAGAGACTGCCTGGGTTAACCAGGAGGAAGGTGGGGATGACGTCAAGTCCTCATGGCCCTTATGCCCAGGGCTACACACGTGCTACAATGGGCCGTACAAAGGGTTGCGAATCCGCGAGGAGGAGCTAATCCCAAAAAGCGGCTCTCAGTTCGGATTGGAGTCTGCAACTCGACTCCATGAAGGCGGAATCGCTAGTAATCGTGGATCAGCATGCCACGGTGAATACGTTCCCGGGCCTTGTACACACCGCCCGTCACACCACGAAAGTCGACTGTACCAGAAGTCGTCGGGCTAACCCCTTTTGGGGAGGCAGGCGCCTAAGGTACGGCCGGTAATTGGGGTG

>GuaymasBasin_Dive4872_L-ASV_207

AACGAACGCTGGTGGCGTGCCTAACACATGCAAGTCGAACGAGAAACTTCACCTTCGGGTGGAGGAGTAAAGTGGCGCACGGGTGAGTAACACGTGGGTAACCTGCCCTTGAGCCCGGGATAACCTCGCGAAAGCGGGGCTAATACCGGATAATGTTTCTGAGGCTTCGGTCTCAGGGATGAAAGGTGACCGCTCCATGGAAGTTGCCACTCAAGGAGGGGCCCGCGGCCTATTAGCTTGTTGGTGGGGTAACGGCTCACCAAGGCAATGATGGGTAGCTGGTCTGAGAGGATGATCAGCCACACTGGCACTGAAACACGGGCCAGACTCCTACGGGAGGCAGCAGTGAGGAATATTGCGCAATGGGGGAAACCCTGACGCAGCAACGCCGTGTGAGCGAAGAAGGCCTTCGGGTCGTAAAGCTCTGTCAGGTGGGAAAAAAGTCTTATGGTTAATAGCCATAGGGTCTGATGGTACCACCAGAGGAAGCACCGGCTAACTCCGTGCCAGCAGCCGCGGTAATACGGAGGGTGCAAGCGTTGTTCGGAATCACTGGGCGTAAAGCGTGTGTAGGCGGTCTGGCAAGTCAGATGTGAAAGACTCCTGCTCAACAGGAGAAGTGCATCTGATACTGTCAGGCTGGAGTATAGGAGAGGAGAGCGGAATTCCCGGTGTAGAGGTGAAATTCGTAGATATCGGGAGGAACACCGGTGGCGAAGGCGGCTCTCTGGCCTAATACTGACGCTGAGACACGAAAGCGTGGGGAGCAAACAGGATTAGAGACCCTGGTAGTCCACGCTGTAAACGGTGAGAACTAGGTGTGGTGGGGTTAAATCCTGCTGTGCCGAAGCTAACGCATTAAGTTCTCCGCCTGGGAACTACGGCCGCAAGGCTAAAACTCAAAGGAATTGACGGGGGCCCGCACAAGCGGTGGAGCATGTGGTTTAATTCGACGCAACGCGAAGAACCTTACCTGGGCTTGACATCCCCTGCCAGCTCTGGAAACAGGGTCTTCCCGGCTTTGCTGGGACAGGGTGACAGGTGCTGCATGGCTGTCGTCAGCTCGTGCCGTGAGGTGTTGGGTTAAGTCCCGCAACGAGCGCAACCCCTGCCCTTAGTTGCCAGTATTAAGTTAGGCACTCTAAGGGGACTGCCTGGGTTAACCAGGAGGAAGGTGGGGACGACGTCAAGTCATCATGGCCTTTATGCCCAGGGCTACACACGTGCTACAATGGGCGGTACAAAGGGTTGCGACACCGCGAGGTGAAGCTAATCCCAAAAAGCCGCCCCCAGTTCGGATTGGAGTCTGCAACTCGACTCCATGAAGGCGGAATCGCTAGTAATCGTGGATCAGAATGCCACGGTGAATACGTTCCCGGGCCTTGTACACACCGCCCGTCACACCACGAAAGTCAGCTGTACCAGAAGTTGCTGGGCTAACCCCTTTATAGGGGAGGCAGGCACCGAAGGTATGGCTGATAATTGGGGTG

>GuaymasBasin_Dive4872_L-ASV_116

AACGAACGCTGGCGGCGTGCTTAACACATGCAAGTCGTACGAGAAACCGGAACTTCGGTTCCGGGAGTAAAGTGGCGCACGGGTGAGTAACGCGTAGGTAATCTACCCTTGGGTTTGGAATAACTCGCCGAAAGGCGTGCTAATACCGGATAAAATCTTTCGGAGGGATCTGGAAGATCAAAGGTGGCCTCTCCATGGATGCTACTGCCTGAGGATGAGCCTGCGTCCCATTAGCTTGTTGGTAGGGTAATGGCCTACCAAGGCTACGATGGGTAGCTGGTCTGAGAGGATGATCAGTCACACTGGAACTGGAACACGGTCCAGACTCCTACGGGAGGCAGCAGTGAGGAATATTGCGCAATGGGGGAAACCCTGACGCAGCGACGCCGCGTGGGCGAAGAAGGCCTTCGGGTCGTAAAGCCCTGTCTAGAGGGAAGAAATTCTCAATGGAATAATACCCATTGAGATTGACGGTACCTCTGGAGGAAGCACCGGCTAACTCCGTGCCAGCAGCCGCGGTAATACGGAGGGTGCTAGCGTTGTTCGGAATTACTGGGCGTAAAGCGGGTGTAGGCGGTTTGTTAAGTCAGATGTGAAAGCCCACGGCTCAACCGTGGAAGTGCATCTGAAACTGGCAGACTTGAGTACCGGAGAGGGAAGTGGAATTCCTGGTGTAGGGGTGAAATCCGTAGATATCAGGAGGAACACCGGTGGCGAAGGCGACTTCCTGGACGGATACTGACGCTGAGATCCGAAAGCGTGGGGAGCAAACAGGATTAGATACCCTGGTAGTCCACGCCCTAAACGATGGGCACTAGGTGCAGGGGGTGTTGACCCCTCCTGTGCCGCAGCTAACGCATTAAGTGCCCCGCCTGGGGAGTACGGCCGCAAGGTTGAAACTCAAAGGAATTGACGGGGGCCCGCACAAGCGGTGGAGCATGTGGTTTAATTCGACGCAACGCGAAGAACCTTACCTGGGCTTGACATCCCGGGAACTCTGTGGAAACACGGAGGTGCCCCTTCGGGGGAACCTGGTGACAGGTGCTGCATGGCTGTCGTCAGCTCGTGTCGTGAGATGTTGGGTTAAGTCCCGCAACGAGCGCAACCCTTGCCTTTAGTTGCCATCATTAAGTTGGGCACTCTAGAGGGACTGCCGGTGTTAAACCGGAGGAAGGTGGGGATGACGTCAAGTCCTCATGGCCCTTATGTCCAGGGCTACACACGTGCTACAATGGTCGGTACAAAGGGCAGCAAACTCGCGAGAGTAAGCAAATCCCAAAAAGCCGATCTCAGTTCGGATCGAAGTCTGCAACTCGACTTCGTGAAGGTGGAATCGCTAGTAATCCCGGATCAGCATGCCGGGGTGAATACGTTCCCGGGCCTTGTACACACCGCCCGTCACACCACGAAAGTTGGCTGTACCAGAAGTCGTTGGGCTAACCCGCAAGGGAGGCAGGCGCCCAAGGTATGGTTAGTGATTGGGGTG

>GuaymasBasin_Dive4872_L-ASV_243

AATGAACGCTGGCGGCGTGCTTAACACATGCAAGTCGTACGAGAAACCGGAACCTCGGTTCCGGGAGTAAAGTGGCGCACGGGTGAGTAACGCGTAGGTAATCTACCCTTGGGTTTGGAATAACCCGCCGAAAGGCGGACTAATACCGGATAAGATCTTTCAGATGGATCTGAAAGATCAAAGGTGGCCTCTCCATGGATGCTACTGCCTGAGGATGAGCCTGCGTCCCATTAGCTTGTTGGTAGGGTAATGGCCTACCAAGGCTACGATGGGTAGCTGGTCTGAGAGGATGATCAGCCACACTGGAACTGGAACACGGTCCAGACTCCTACGGGAGGCAGCAGTGAGGAATATTGCGCAATGGGGGAAACCCTGACGCAGCGACGCCGCGTGGGTGAAGAAGGCCTTCGGGTCGTAAAGCCCTGTCTAGAGGGAAGAAATTCTCAATGGAATAATACCTGTTGAGATTGACGGTACCTCTGGAGGAAGCACCGGCTAACTCCGTGCCAGCAGCCGCGGTAATACGGAGGGTGCGAGCGTTGTTCGGAATTACTGGGCGTAAAGCGGGTGTAGGCGGTTTGTTAAGTCAGATGTGAAAGCCCACGGCTCAACCGTGGAAGTGCATCTGAAACTGGCAGACTTGAGTACCGGAGAGGGAAGTGGAATTCCTGGTGTAGGGGTGAAATCCGTAGATATCAGGAGGAACACCGGTGGCGAAGGCGACTTCCTGGACGGATACTGACGCTGAGATCCGAAAGCGTGGGGAGCAAACAGGATTAGATACCCTGGTAGTCCACGCCCTAAACGATGGGCACTAGGTGCAGGGGGTGTTGACCCCTCCTGTGCCGCAGCTAACGCATTAAGTGCCCCGCCTGGGGAGTACGGCCGCAAGGTTGAAACTCAAAGGAATTGACGGGGGCCCGCACAAGCGGTGGAGCATGTGGTTTAATTCGACGCAACGCGAAGAACCTTACCTGGGCTTGACATCCCGGGAACTCTGTGGAAACATGGAGGTGCCCCTTCGGGGGAACCTGGTGACAGGTGCTGCATGGCTGTCGTCAGCTCGTGTCGTGAGATGTTGGGTTAAGTCCCGCAACGAGCGCAACCCTTGCCTTTAGTTGCCATCATTAAGTTGGGCACTCTAGAGGGACTGCCGGTGTTAAACCGGAGGAAGGTGGGGATGACGTCAAGTCCTCATGGCCCTTATGTCCAGGGCTACACACGTGCTACAATGGTCGGTACAAAGGGCAGCAAACTCGCGAGAGTAAGCAAATCCCAAAAAGCCGATCTCAGTTCGGATCGAAGTCTGCAACTCGACTTCGTGAAGGTGGAATCGCTAGTAATCCCGGATCAGCATGCCGGGGTGAATACGTTCCCGGGCCTTGTACACACCGCCCGTCACACCACGAAAGTTGGCTGTACCAGAAGTCGTTGGGCTAACCCGCAAGGGAAGCAGGCGCCCAAGGTATGGTCAGTGATTGGGGTG

>GuaymasBasin_Dive4872_L-ASV_290

AATGAACGCTGGCGGCGTGCTTAACACATGCAAGTCGTACGAGAAACCGGAACTTCGGTTCCGGGAGTAAAGTGGCGCACGGGTGAGTAACGCGTAGGTAATCTACCCTTGGGTTTGGAATAACCCGCCGAAAGGCGGACTAATACCGGATAAGATCTTTCAGATGGATCTGAAAGATCAAAGGTGGCCTCTCCATGGATGCTACTGCCTGAGGATGAGCCTGCGTCCCATTAGCTTGTTGGTAGGGTAATGGCCTACCAAGGCTACGATGGGTAGCTGGTCTGAGAGGATGATCAGCCACACTGGAACTGGAACACGGTCCAGACTCCTACGGGAGGCAGCAGTGAGGAATATTGCGCAATGGGGGAAACCCTGACGCAGCGACGCCGCGTGGGTGAAGAAGGCCTTCGGGTCGTAAAGCCCTGTCTAGAGGGAAGAAATTCTCAATGGAATAATACCTGTTGAGATTGACGGTACCTCTGGAGGAAGCACCGGCTAACTCCGTGCCAGCAGCCGCGGTAATACGGAGGGTGCGAGCGTTGTTCGGAATTACTGGGCGTAAAGCGGGTGTAGGCGGTTTGTTAAGTCAGATGTGAAAGCCCACGGCTCAACCGTGGAAGTGCATCTGAAACTGGCAGACTTGAGTACCGGAGAGGGAAGTGGAATTCCTGGTGTAGGGGTGAAATCCGTAGATATCAGGAGGAACACCGGTGGCGAAGGCGACTTCCTGGACGGATACTGACGCTGAGATCCGAAAGCGTGGGGAGCAAACAGGATTAGATACCCTGGTAGTCCACGCCCTAAACGATGGGCACTAGGTGCAGGGGGTGTTGACCCCTCCTGTGCCGCAGCTAACGCATTAAGTGCCCCGCCTGGGGAGTACGGCCGCAAGGTTGAAACTCAAAGGAATTGACGGGGGCCCGCACAAGCGGTGGAGCATGTGGTTTAATTCGACGCAACGCGAAGAACCTTACCTGGGCTTGACATCCCGGGAACTCTGTGGAAACATGGAGGTGCCCCTTCGGGGGAACCTGGTGACAGGTGCTGCATGGCTGTCGTCAGCTCGTGTCGTGAGATGTTGGGTTAAGTCCCGCAACGAGCGCAACCCTTGCCTTTAGTTGCCATCATTAAGTTGGGCACTCTAGAGGGACTGCCGGTGTTAAACCGGAGGAAGGTGGGGATGACGTCAAGTCCTCATGGCCCTTATGTCCAGGGCTACACACGTGCTACAATGGTCGGTACAAAGGGCAGCAAACTCGCGAGAGTAAGCAAATCCCAAAAAGCCGATCTCAGTTCGGATCGAAGTCTGCAACTCGACTTCGTGAAGGTGGAATCGCTAGTAATCCCGGATCAGCATGCCGGGGTGAATACGTTCCCGGGCCTTGTACACACCGCCCGTCACACCACGAAAGTTGGCTGTACCAGAAGTCGTTGGGCTAACCCGCAAGGGAAGCAGGCGCCCAAGGTATGGTCAGTGATTGGGGTG

>GuaymasBasin_Dive4872_L-ASV_1074

AATGAACGCTGGCGGCGTGCTTAACACATGCAAGTCGTACGAGAAACCGGGACTTCGGTTCTGGGAGTAAAGTGGCGCACGGGTGAGTAACGCGTAGGTAATCTACCCTTGGGTTTGGAATAACCCGCCGAAAGGCGGGCTAATACCGGATAAAATCTTTCGGAGGGATCCGGCGGATCAAAGGTGGCCTCTCCATGGATGCTACTGCCTGAGGATGAGCCTGCGTCCCATTAGCTTGTTGGTTGGGTAATGGCCTACCAAGGCTACGATGGGTAGCTGGTCTGAGAGGATGATCAGTCACACTGGAACTGGAACACGGTCCAGACTCCTACGGGAGGCAGCAGTGAGGAATATTGCGCAATGGGGGAAACCCTGACGCAGCGACGCCGCGTGGGTGAAGAAGGCCTTCGGGTCGTAAAGCCCTGTCTAGAGGGAAGAACTTCTCAAGGGAACAATACCCGTTGAGATTGACGGTACCTCTGGAGGAAGCACCGGCTAACTCCGTGCCAGCAGCCGCGGTAATACGGAGGGTGCGAGCGTTGTTCGGAATTACTGGGCGTAAAGTGGGTGTAGGCGGCTTGTTAAGTCAGATGTGAAAGCCCACGGCTCAACCGTGGAAGTGCATCTGAAACTGGCAGGCTTGAGTACCGGAGAGGGAAGTGGAATTCCTGGTGTAGGGGTGAAATCCGTAGATATCAGGAGGAACACCGGTGGCGAAGGCGACTTCCTGGACGGATACTGACGCTGAGATCCGAAAGCGTGGGGAGCAAACAGGATTAGATACCCTGGTAGTCCACGCCCTAAACGATGGGCACTAGGTGCAGGGGGTGTTGACCCCTTCTGTGCCGCAGCTAACGCATTAAGTGCCCCGCCTGGGGAGTACGGCCGCAAGGTTGAAACTCAAAGGAATTGACGGGGGCCCGCACAAGCGGTGGAGCATGTGGTTTAATTCGACGCAACGCGAAGAACCTTACCTGGGCTTGACATCCCGGGAACTCTGTGGAAACATGGAGGTGCCCCTTCGGGGGAACCCGGTGACAGGTGCTGCATGGCTGTCGTCAGCTCGTGTCGTGAGATGTTGGGTTAAGTCCCGCAACGAGCGCAACCCTTGCCTTTAGTTGCCATCATTAAGTTGGGCACTCTAGAGGGACTGCCGGTGTTAAACCGGAGGAAGGTGGGGATGACGTCAAGTCCTCATGGCCCTTATGTCCAGGGCTACACACGTGCTACAATGGTCGGTACAAAGGGCAGCAAACTCGCGAGAGCAAGCAAATCCCCAAAAGCCGATCTCAGTTCGGATCGAAGTCTGCAACTCGACTTCGTGAAGGTGGAATCGCTAGTAATCCCGGATCAGCATGCCGGGGTGAATACGTTCCCGGGCCTTGTACACACCGCCCGTCACACCACGAAAGTTGGCTGTACCAGAAGTCGTTGGGCTAACCCGCAAGGGAGGCAGGCGCCCAAGGTATGGTCAGTGATTGGGGTG

>GuaymasBasin_Dive4872_L-ASV_608

AACGAACGCTGGCGGCGTGCCTAACACATGCAAGTCGTACGAGAAACCGGGACTTCGGTTCCGGGAGTAAAGTGGCGCACGGGTGAGTAACGCGTAGGTAATCTACCCTTGGGTCTGGAATAACCCGCCGAAAGGCGTGCTAATACCGGATAAAATCCTTCGGAGGGATCCGGCGGATCAAAGGTGGCCTCTCCATGGATGCTACTGCCTGAGGATGAGCCTGCGTCCCATTAGCTTGTTGGTAGGGTAATGGCCTACCAAGGCTACGATAGGTAGCTGGTCTGAGAGGATGATCAGCCACACTGGGACTGGAACACGGTCCAGACTCCTACGGGAGGCAGCAGTGAGGAATATTGCGCAATGGGGGAAACCCTGACGCAGCGACGCCGCGTGGGTGAAGAAGGCCTTCGGGTCGTAAAGCCCTGTCTGGAGGGAAGAACTTCTCAATGGAATAATACCTGTTGAGATTGACGGTACCTCTGGAGGAAGCACCGGCTAACTCCGTGCCAGCAGCCGCGGTAATACGGAGGGTGCGAGCGTTGTTCGGAATTACTGGGCGTAAAGGGGATGTAGGCGGTTTGTTAAGTCAGATGTGAAAGCCCACGGCTCAACCGTGGAAGTGCATCTGAAACTGGCAGACTTGAGTACCGGAGAGGGAAGTGGAATTCCTGGTGTAGGGGTGAAATCCATAGATATCAGGAGGAACACCGGTGGCGAAGGCGACTTCCTGGACGGATACTGACGCTGAGATCCGAAAGCGTGGGGAGCAAACAGGATTAGATACCCTGGTAGTCCACGCCGTAAACGATGGGCACTAGGTGCAGGGGGTGTTGACCCCTCCTGTGCCGCAGCTAACGCATTAAGTGCCCCGCCTGGGGAGTACGGCCGCAAGGTTGAAACTCAAAGGAATTGACGGGGGCCCGCACAAGCGGTGGAGCATGTGGTTTAATTCGACGCAACGCGAAGAACCTTACCTGGGCTTGACATCCCGGGAACTCTGTGGAAACACGGGGGTGCCCCTTCGGGGGAACCTGGTGACAGGTGCTGCATGGCTGTCGTCAGCTCGTGTCGTGAGATGTTGGGTTAAGTCCCGCAACGAGCGCAACCCTTGCCTTTAGTTGCCATCATTAAGTTGGGCACTCTAGAGGGACTGCCGGTGCTAAACCGGAGGAAGGTGGGGATGACGTCAAGTCCTCATGGCCCTTATGCCCAGGGCTACACACGTGCTACAATGGTCGGTACAAAGGGCAGCGAACTCGCGAGAGCAAGCAAATCCCCAAAAGCCGATCTCAGTTCGGATCGAAGTCTGCAACTCGACTTCGTGAAGGTGGAATCGCTAGTAATCCCGGATCAGCATGCCGGGGTGAATACGTTCCCGGGCCTTGTACACACCGCCCGTCACACCACGAAAGTTGGCTGTACCAGAAGTCGTTGGGCTAACCCGCAAGGGAGGCAGGCGCCCAAGGTATGGTCAGTGATTGGGGTG

>GuaymasBasin_Dive4872_L-ASV_609

AACGAACGCTGGCGGCGTGCCTAACACATGCAAGTCGTACGAGAAACCGGGACTTCGGTTCCGGGAGTAAAGTGGCGCACGGGTGAGTAACGCGTAGGTAATCTACCCTTGGGTCTGGAATAACCCGCCGAAAGGCGTGCTAATACCGGATAAAATCCTTCGGAGGGATCCGGCGGATCAAAGGTGGCCTCTCCATGGATGCTACTGCCTGAGGATGAGCCTGCGTCCCATTAGCTTGTTGGTAGGGTAATGGCCTACCAAGGCTACGATAGGTAGCTGGTCTGAGAGGATGATCAGCCACACTGGGACTGGAACACGGTCCAGACTCCTACGGGAGGCAGCAGTGAGGAATATTGCGCAATGGGGGAAACCCTGACGCAGCGACGCCGCGTGGGTGAAGAAGGCCTTCGGGTCGTAAAGCCCTGTCTGGAGGGAAGAACTTCTCAATGGAATAATACCTGTTGAGATTGACGGTACCTCTGGAGGAAGCACCGGCTAACTCCGTGCCAGCAGCCGCGGTAATACGGAGGGTGCAAGCGTTGTTCGGAATTACTGGGCGTAAAGGGGATGTAGGCGGTTTGTTAAGTCAGATGTGAAAGCCCACGGCTCAACCGTGGAAGTGCATCTGAAACTGGCAGACTTGAGTACCGGAGAGGGAAGTGGAATTCCTGGTGTAGGGGTGAAATCCATAGATATCAGGAGGAACACCGGTGGCGAAGGCGACTTCCTGGACGGATACTGACGCTGAGATCCGAAAGCGTGGGGAGCAAACAGGATTAGATACCCTGGTAGTCCACGCCCTAAACGATGGGCACTAGGTGCAGGGGGTGTTGACCCCTCCTGTGCCGCAGCTAACGCATTAAGTGCCCCGCCTGGGGAGTACGGCCGCAAGGTTGAAACTCAAAGGAATTGACGGGGGCCCGCACAAGCGGTGGAGCATGTGGTTTAATTCGACGCAACGCGAAGAACCTTACCTGGGCTTGACATCCCGGGAACTCTGTGGAAACACGGGGGTGCCCCTTCGGGGGAACCTGGTGACAGGTGCTGCATGGCTGTCGTCAGCTCGTGTCGTGAGATGTTGGGTTAAGTCCCGCAACGAGCGCAACCCTTGCCTTTAGTTGCCATCATTAAGTTGGGCACTCTAGAGGGACTGCCGGTGCTAAACCGGAGGAAGGTGGGGATGACGTCAAGTCCTCATGGCCCTTATGCCCAGGGCTACACACGTGCTACAATGGTCGGTACAAAGGGCAGCGAACTCGCGAGAGCAAGCAAATCCCCAAAAGCCGATCTCAGTTCGGATCGAAGTCTGCAACTCGACTTCGTGAAGGTGGAATCGCTAGTAATCCCGGATCAGCATGCCGGGGTGAATACGTTCCCGGGCCTTGTACACACCGCCCGTCACACCACGAAAGTTGGCTGTACCAGAAGTCGTTGGGCTAACCCGCAAGGGAGGCAGGCGCCCAAGGTATGGTCAGTGATTGGGGTG

>GuaymasBasin_Dive4872_L-ASV_51

AACGAACGCTGGCGGCGTGCCTAACACATGCAAGTCGTACGAGAAACCGGGACTTCGGTTCCGGGAGTAAAGTGGCGCACGGGTGAGTAACGCGTAGGTAATCTACCCTTGGGTCTGGAATAACCCGCCGAAAGGCGTGCTAATACCGGATAAAATCCTTCGGAGGGATCCGGCGGATCAAAGGTGGCCTCTCCATGGATGCTACTGCCTGAGGATGAGCCTGCGTCCCATTAGCTTGTTGGTAGGGTAATGGCCTACCAAGGCTACGATAGGTAGCTGGTCTGAGAGGATGATCAGCCACACTGGGACTGGAACACGGTCCAGACTCCTACGGGAGGCAGCAGTGAGGAATATTGCGCAATGGGGGAAACCCTGACGCAGCGACGCCGCGTGGGTGAAGAAGGCCTTCGGGTCGTAAAGCCCTGTCTGGAGGGAAGAACTTCTCAATGGAATAATACCTGTTGAGATTGACGGTACCTCTGGAGGAAGCACCGGCTAACTCCGTGCCAGCAGCCGCGGTAATACGGAGGGTGCGAGCGTTGTTCGGAATTACTGGGCGTAAAGGGGATGTAGGCGGTTTGTTAAGTCAGATGTGAAAGCCCACGGCTCAACCGTGGAAGTGCATCTGAAACTGGCAGACTTGAGTACCGGAGAGGGAAGTGGAATTCCTGGTGTAGGGGTGAAATCCATAGATATCAGGAGGAACACCGGTGGCGAAGGCGACTTCCTGGACGGATACTGACGCTGAGATCCGAAAGCGTGGGGAGCAAACAGGATTAGATACCCTGGTAGTCCACGCCCTAAACGATGGGCACTAGGTGCAGGGGGTGTTGACCCCTCCTGTGCCGCAGCTAACGCATTAAGTGCCCCGCCTGGGGAGTACGGCCGCAAGGTTGAAACTCAAAGGAATTGACGGGGGCCCGCACAAGCGGTGGAGCATGTGGTTTAATTCGACGCAACGCGAAGAACCTTACCTGGGCTTGACATCCCGGGAACTCTGTGGAAACACGGGGGTGCCCCTTCGGGGGAACCTGGTGACAGGTGCTGCATGGCTGTCGTCAGCTCGTGTCGTGAGATGTTGGGTTAAGTCCCGCAACGAGCGCAACCCTTGCCTTTAGTTGCCATCATTAAGTTGGGCACTCTAGAGGGACTGCCGGTGCTAAACCGGAGGAAGGTGGGGATGACGTCAAGTCCTCATGGCCCTTATGCCCAGGGCTACACACGTGCTACAATGGTCGGTACAAAGGGCGGCGAACTCGCGAGAGCAAGCAAATCCCCAAAAGCCGATCTCAGTTCGGATCGAAGTCTGCAACTCGACTTCGTGAAGGTGGAATCGCTAGTAATCCCGGATCAGCATGCCGGGGTGAATACGTTCCCGGGCCTTGTACACACCGCCCGTCACACCACGAAAGTTGGCTGTACCAGAAGTCGTTGGGCTAACCCGCAAGGGAGGCAGGCGCCCAAGGTATGGTCAGTGATTGGGGTG

>GuaymasBasin_Dive4872_L-ASV_5

AACGAACGCTGGCGGCGTGCCTAACACATGCAAGTCGTACGAGAAACCGGGACTTCGGTTCCGGGAGTAAAGTGGCGCACGGGTGAGTAACGCGTAGGTAATCTACCCTTGGGTCTGGAATAACCCGCCGAAAGGCGTGCTAATACCGGATAAAATCCTTCGGAGGGATCCGGCGGATCAAAGGTGGCCTCTCCATGGATGCTACTGCCTGAGGATGAGCCTGCGTCCCATTAGCTTGTTGGTAGGGTAATGGCCTACCAAGGCTACGATAGGTAGCTGGTCTGAGAGGATGATCAGCCACACTGGGACTGGAACACGGTCCAGACTCCTACGGGAGGCAGCAGTGAGGAATATTGCGCAATGGGGGAAACCCTGACGCAGCGACGCCGCGTGGGTGAAGAAGGCCTTCGGGTCGTAAAGCCCTGTCTGGAGGGAAGAACTTCTCAATGGAATAATACCTGTTGAGATTGACGGTACCTCTGGAGGAAGCACCGGCTAACTCCGTGCCAGCAGCCGCGGTAATACGGAGGGTGCGAGCGTTGTTCGGAATTACTGGGCGTAAAGGGGATGTAGGCGGTTTGTTAAGTCAGATGTGAAAGCCCACGGCTCAACCGTGGAAGTGCATCTGAAACTGGCAGACTTGAGTACCGGAGAGGGAAGTGGAATTCCTGGTGTAGGGGTGAAATCCATAGATATCAGGAGGAACACCGGTGGCGAAGGCGACTTCCTGGACGGATACTGACGCTGAGATCCGAAAGCGTGGGGAGCAAACAGGATTAGATACCCTGGTAGTCCACGCCCTAAACGATGGGCACTAGGTGCAGGGGGTGTTGACCCCTCCTGTGCCGCAGCTAACGCATTAAGTGCCCCGCCTGGGGAGTACGGCCGCAAGGTTGAAACTCAAAGGAATTGACGGGGGCCCGCACAAGCGGTGGAGCATGTGGTTTAATTCGACGCAACGCGAAGAACCTTACCTGGGCTTGACATCCCGGGAACTCTGTGGAAACACGGGGGTGCCCCTTCGGGGGAACCTGGTGACAGGTGCTGCATGGCTGTCGTCAGCTCGTGTCGTGAGATGTTGGGTTAAGTCCCGCAACGAGCGCAACCCTTGCCTTTAGTTGCCATCATTAAGTTGGGCACTCTAGAGGGACTGCCGGTGCTAAACCGGAGGAAGGTGGGGATGACGTCAAGTCCTCATGGCCCTTATGCCCAGGGCTACACACGTGCTACAATGGTCGGTACAAAGGGCAGCGAACTCGCGAGAGCAAGCAAATCCCCAAAAGCCGATCTCAGTTCGGATCGAAGTCTGCAACTCGACTTCGTGAAGGTGGAATCGCTAGTAATCCCGGATCAGCATGCCGGGGTGAATACGTTCCCGGGCCTTGTACACACCGCCCGTCACACCACGAAAGTTGGCTGTACCAGAAGTCGTTGGGCTAACCCGCAAGGGAGGCAGGCGCCCAAGGTATGGTCAGTGATTGGGGTG

>GuaymasBasin_Dive4872_L-ASV_605

AACGAACGCTGGCGGCGTGCCTAACACATGCAAGTCGTACGAGAAACCGGGACTTCGGTTCCGGGAGTAAAGTGGCGCACGGGTGAGTAACGCGTAGGTAATCTACCCTTGGGTCTGGAATAACCCGCCGAAAGGCGTGCTAATACCGGATAAAATCCTTCGGAGGGATCCGGCGGATCAAAGGTGGCCTCTCCATGGATGCTACTGCCTGAGGATGAGCCTGCGTCCCATTAGCTTGTTGGTAGGGTAATGGCCTACCAAGGCTACGATAGGTAGCTGGTCTGAGAGGATGATCAGCCACACTGGGACTGGAACACGGTCCAGACTCTTACGGGAGGCAGCAGTGAGGAATATTGCGCAATGGGGGAAACCCTGACGCAGCGACGCCGCGTGGGTGAAGAAGGCCTTCGGGTCGTAAAGCCCTGTCTGGAGGGAAGAACTTCTCAATGGAATAATACCTGTTGAGATTGACGGTACCTCTGGAGGAAGCACCGGCTAACTCCGTGCCAGCAGCCGCGGTAATACGGAGGGTGCGAGCGTTGTTCGGAATTACTGGGCGTAAAGGGGATGTAGGCGGTTTGTTAAGTCAGATGTGAAAGCCCACGGCTCAACCGTGGAAGTGCATCTGAAACTGGCAGACTTGAGTACCGGAGAGGGAAGTGGAATTCCTGGTGTAGGGGTGAAATCCATAGATATCAGGAGGAACACCGGTGGCGAAGGCGACTTCCTGGACGGATACTGACGCTGAGATCCGAAAGCGTGGGGAGCAAACAGGATTAGATACCCTGGTAGTCCACGCCCTAAACGATGGGCACTAGGTGCAGGGGGTGTTGACCCCTCCTGTGCCGCAGCTAACGCATTAAGTGCCCCGCCTGGGGAGTACGGCCGCAAGGTTGAAACTCAAAGGAATTGACGGGGGCCCGCACAAGCGGTGGAGCATGTGGTTTAATTCGACGCAACGCGAAGAACCTTACCTGGGCTTGACATCCCGGGAACTCTGTGGAAACACGGGGGTGCCCCTTCGGGGGAACCTGGTGACAGGTGCTGCATGGCTGTCGTCAGCTCGTGTCGTGAGATGTTGGGTTAAGTCCCGCAACGAGCGCAACCCTTGCCTTTAGTTGCCATCATTAAGTTGGGCACTCTAGAGGGACTGCCGGTGCTAAACCGGAGGAAGGTGGGGATGACGTCAAGTCCTCATGGCCCTTATGCCCAGGGCTACACACGTGCTACAATGGTCGGTACAAAGGGCAGCGAACTCGCGAGAGCAAGCAAATCCCCAAAAGCCGATCTCAGTTCGGATCGAAGTCTGCAACTCGACTTCGTGAAGGTGGAATCGCTAGTAATCCCGGATCAGCATGCCGGGGTGAATACGTTCCCGGGCCTTGTACACACCGCCCGTCACACCACGAAAGTTGGCTGTACCAGAAGTCGTTGGGCTAACCCGCAAGGGAGGCAGGCGCCCAAGGTATGGTCAGTGATTGGGGTG

>GuaymasBasin_Dive4872_L-ASV_920

AACGAACGCTGGCGGCGTGCCTAACACATGCAAGTCGTACGAGAAACCGGGACTTCGGTTCCGGGAGTAAAGTGGCGCACGGGTGAGTAACGCGTAGGTAATCTACCCTTGGGTCTGGAATAACCCGCCGAAAGGCGTGCTAATACCGGATAAAATCCTTCGGAGGGATCCGGCGGATCAAAGGTGGCCTCTCCATGGATGCTACTGCCTGAGGATGAGCCTGTGTCCCATTAGCTTGTTGGTAGGGTAATGGCCTACCAAGGCTACGATAGGTAGCTGGTCTGAGAGGATGATCAGCCACACTGGGACTGGAACACGGTCCAGACTCCTACGGGAGGCAGCAGTGAGGAATATTGCGCAATGGGGGAAACCCTGACGCAGCGACGCCGCGTGGGTGAAGAAGGCCTTCGGGTCGTAAAGCCCTGTCTGGAGGGAAGAACTTCTCAATGGAATAATACCTGTTGAGATTGACGGTACCTCTGGAGGAAGCACCGGCTAACTCCGTGCCAGCAGCCGCGGTAATACGGAGGGTGCGAGCGTTGTTCGGAATTACTGGGCGTAAAGGGGATGTAGGCGGTTTGTTAAGTCAGATGTGAAAGCCCACGGCTCAACCGTGGAAGTGCATCTGAAACTGGCAGACTTGAGTACCGGAGAGGGAAGTGGAATTCCTGGTGTAGGGGTGAAATCCATAGATATCAGGAGGAACACCGGTGGCGAAGGCGACTTCCTGGACGGATACTGACGCTGAGATCCGAAAGCGTGGGGAGCAAACAGGATTAGATACCCTGGTAGTCCACGCCCTAAACGATGGGCACTAGGTGCAGGGGGTGTTGACCCCTCCTGTGCCGCAGCTAACGCATTAAGTGCCCCGCCTGGGGAGTACGGCCGCAAGGTTGAAACTCAAAGGAATTGACGGGGGCCCGCACAAGCGGTGGAGCATGTGGTTTAATTCGACGCAACGCGAAGAACCTTACCTGGGCTTGACATCCCGGGAACTCTGTGGAAACACGGGGGTGCCCCTTCGGGGGAACCTGGTGACAGGTGCTGCATGGCTGTCGTCAGCTCGTGTCGTGAGATGTTGGGTTAAGTCCCGCAACGAGCGCAACCCTTGCCTTTAGTTGCCATCATTAAGTTGGGCACTCTAGAGGGACTGCCGGTGCTAAACCGGAGGAAGGTGGGGATGACGTCAAGTCCTCATGGCCCTTATGCCCAGGGCTACACACGTGCTACAATGGTCGGTACAAAGGGCAGCGAACTCGCGAGAGCAAGCAAATCCCCAAAAGCCGATCTCAGTTCGGATCGAAGTCTGCAACTCGACTTCGTGAAGGTGGAATCGCTAGTAATCCCGGATCAGCATGCCGGGGTGAATACGTTCCCGGGCCTTGTACACACCGCCCGTCACACCACGAAAGTTGGCTGTACCAGAAGTCGTTGGGCTAACCCGCAAGGGAGGCAGGCGCCCAAGGTATGGTCAGTGATTGGGGTG

>GuaymasBasin_Dive4872_L-ASV_818

AACGAACGCTGGCGGCGTGCTTAACACATGCAAGTCGAACGCGAAAGTTTTCTTCGGAAGATGAGTAGAGTGGCGCACGGGTGAGTAACGCGTAAATAATCTACCCCTGCATCCGGGATAACCCACCGAAAGGTGTGCTAATACCGGATACGTTCTCTTTATCGCGAGATAGAGAGAAGAAAGGTGGCCTCTGATATAAGCTACTGTGCGGGGAGGAGTTTGCGTACCATTAGCTAGTTGGTAGGGTAATGGCCTACCAAGGCAACGATGGTTAGCGGGTCTGAGAGGATGATCCGCCACACTGGAACTGGAACACGGACCAGACTCCTACGGGAGGCAGCAGTGAGGAATATTGCGCAATGGGGGCAACCCTGACGCAGCGACGCCGCGTGGATGATGAAGGCCTTCGGGTCGTAAAATCCTGTCAGATGGAAAGAAATGTTATATGGATAATACCTGTATAGCTTGACGGTACCATCAAAGGAAGCACCGGCTAACTCCGTGCCAGCAGCCGCGGTAATACGGAGGGTGCAAGCGTTGTTCGGAATTACTGGGCGTAAAGCGCGCGTAGGTGGTCTGTTATGTCAGATGTGAAAGTCCACGGCTCAACCGTGGAAGTGCATTTGAAACTGGCAGACTTGAGTACTGGAGGGGGTAGTGGAATTCCCGGTGTAGAGGTGAAATTCGTAGATATCGGGAGGAATACCGGTGGCGAAGGCGACTACCTGGCCAGATACTGACACTGAGGTGCGAAAGCGTGGGGAGCGAACAGGATTAGATACCCTGGTAGTCCACGCCGTAAACGATGTCAACTAGGTGTTGGGATGGTTAATCGTCTCATTGCCGCAGCTAACGCATTAAGTTGACCGCCTGGGGAGTACGGTCGCAAGATTAAAACTCAAAGGAATTGACGGGGGCCCGCACAAGCGGTGGAGTATGTGGTTTAATTCGACGCAACGCGCAGAACCTTACCTGGTCTTGACATCCCGGGAAGCTTCAGGAAACTGGAGTGTGCCTCTTGAGGAACCCGGTGACAGGTGCTGCATGGCTGTCGTCAGCTCGTGTCGTGAGATGTTGGGTTAAGTCCCGCAACGAGCGCAACCCTTGTCTTTAGTTGCCATCATTAAGTTGGGCACTCTAAAGAGACTGCCGGTGTCAAACCGGAGGAAGGTGGGGATGACGTCAAGTCCTCATGGCCTTTATGACCAGGGCTACACACGTACTACAATGGCATAGACAAAGGGCAGCGACATCGCAAGGTGAAGCGAATCCCATAAACTATGTCTCAGTCCGGATTGGAGTCTGCAACTCGACTCCATGAAGTTGGAATCGCTAGTAATCGTAGATCAGCATGCTACGGTGAATACGTTCCCGGGCCTTGTACACACCGCCCGTCACACCACGGGAGTTGGTTGTACCAGAAGCAGTTGAGCGAACTATTCGTAGACGCAGGCTGCCAAGGTATGATTGGTAACTGGGGTG

>GuaymasBasin_Dive4872_L-ASV_693

AACGAACGCTGGCGGCGTGCTTAACACATGCAAGTCGAACGCGAAAGTTTTCTTCGGAAGATGAGTAGAGTGGCGCACGGGTGAGTAACGCGTAAATAATCTACCCCTGCATTCGGGATAACCCACCGAAAGGTGTGCTAATACCGGATACGTTCTCTTTATCGCGAGATAAGGAGAAGAAAGGTGGCCTCTGATATAAGCTACTGTGCGGGGAGGAGTTTGCGTACCATTAGCTAGTTGGTAGGGTAATGGCCTACCAAGGCAACGATGGTTAGCGGGTCTGAGAGGATGATCCGCCACACTGGAACTGGAACACGGACCAGACTCCTACGGGAGGCAGCAGTGAGGAATATTGCGCAATGGGGGCAACCCTGACGCAGCGACGCCGCGTGGATGATGAAGGCCTTCGGGTCGTAAAATCCTGTCAGATGGAAAGAAATGTTATATGGATAATACCTGTATAGCTTGACGGTACCATCAAAGGAAGCACCGGCTAACTCCGTGCCAGCAGCCGCGGTAATACGGAGGGTGCAAGCGTTGTTCGGAATTACTGGGCGTAAAGCGCGCGTAGGTGGTCTGTTATGTCAGATGTGAAAGTCCACGGCTCAACCGTGGAAGTGCATTTGAAACTGGCAGACTTGAGTACTGGAGGGGGTAGTGGAATTCCCGGTGTAGAGGTGAAATTCGTAGATATCGGGAGGAATACCGGTGGCGAAGGCGACTACCTGGCCAGATACTGACACTGAGGTGCGAAAGCGTGGGGAGCGAACAGGATTAGATACCCTGGTAGTCCACGCCGTAAACGATGTCAACTAGGTGTTGGGATGGTTAATCGTCTCATTGCCGCAGCTAACGCATTAAGTTGACCGCCTGGGGAGTACGGTCGCAAGATTAAAACTCAAAGGAATTGACGGGGGCCCGCACAAGCGGTGGAGTATGTGGTTTAATTCGACGCAACGCGCAGAACCTTACCTGGTCTTGACATCCCGGGAAGCTCCAGGAAACTGGAGTGTGCCTCTTGAGGAACCCGGTGACAGGTGCTGCATGGCTGTCGTCAGCTCGTGTCGTGAGATGTTGGGTTAAGTCCCGCAACGAGCGCAACCCTTGTCTTTAGTTGCCATCATTAAGTTGGGCACTCTAAAGAGACTGCCGGTGTCAAACCGGAGGAAGGTGGGGATGACGTCAAGTCCTCATGGCCTTTATGACCAGGGCTACACACGTACTACAATGGCATAGACAAAGGGCAGCGACATCGCGAGGTGAAGCGAATCCCATAAACTATGTCTCAGTCCGGATTGGAGTCTGCAACTCGACTCCATGAAGTTGGAATCGCTAGTAATCGTAGATCAGCATGCTACGGTGAATACGTTCCCGGGCCTTGTACACACCGCCCGTCACACCACGGGAGTTGGTTGTACCAGAAGCAGTTGAGCGAACTATTCGTAGACGCAGGCTGCCAAGGTATGATTGGTAACTGGGGTG

>GuaymasBasin_Dive4872_L-ASV_386

AACGAACGCTGGCGGCGTGCTTAACACATGCAAGTCGAACGCGAAAGTTTTCTTTCGAGAAAATGAGTAGAGTGGCGCACGGGTGAGTAACGCGTAAATAATCTACCCCTGCATCTGGGATAACCCACCGAAAGGTGTGCTAATACCGGATACGTTCTTTTGATCGCGAGATCTTAAGAAGAAAGGTGGCCTCTGATATAAGCTACTGTGCGGGGAGGAGTTTGCGTACCATTAGCTAGTTGGTAGGGTAATGGCCTACCAAGGCAACGATGGTTAGCGGGTCTGAGAGGATGATCCGCCACACTGGAACTGGAACACGGACCAGACTCCTACGGGAGGCAGCAGTGAGGAATATTGCGCAATGGGGGCAACCCTGACGCAGCGACGCCGCGTGGATGATGAAGGCCTTCGGGTCGTAAAATCCTGTCAGATGGAAAGAAATGTATATGTATTAATACTATGTATGCTTGACGGTACCATCAAAGGAAGCACCGGCTAACTCCGTGCCAGCAGCCGCGGTAATACGGAGGGTGCAAGCGTTGTTCGGAATTACTGGGCGTAAAGCGCGCGTAGGTGGTCTGTTATGTCAGATGTGAAAGTCCACGGCTCAACCGTGGAAGTGCATTTGAAACTGGCAGACTTGAGTACTGGAGGGGGTAGTGGAATTCCCGGTGTAGAGGTGAAATTCGTAGATATCGGGAGGAATACCGGTGGCGAAGGCGACTACCTGGCCAGATACTGACACTGAGGTGCGAAAGCGTGGGGAGCGAACAGGATTAGATACCCTGGTAGTCCACGCCGTAAACGATGTCAACTAGGTGTTGGGATGGTTAATCGTCTCATTGCCGCAGCTAACGCATTAAGTTGACCGCCTGGGGAGTACGGTCGCAAGATTAAAACTCAAAGGAATTGACGGGGGCCCGCACAAGCGGTGGAGTATGTGGTTTAATTCGACGCAACGCGCAGAACCTTACCTGGTCTTGACATCCCGGGAAGCTCTAGGAAACTAGAGCGTGCCTCTTGAGGAACCCGGTGACAGGTGCTGCATGGCTGTCGTCAGCTCGTGTCGTGAGATGTTGGGTTAAGTCCCGCAACGAGCGCAACCCTTGTCTTTAGTTGCCATCATTAAGTTGGGCACTCTAAAGAGACTGCCGGTGTCAAACCGGAGGAAGGTGGGGATGACGTCAAGTCCTCATGGCCTTTATGACCAGGGCTACACACGTACTACAATGGCATAGACAAAGGGCAGCGACATCGCGAGGTGAAGCGAATCCCATAAACTATGTCTCAGTCCGGATTGGAGTCTGCAACTCGACTCCATGAAGTTGGAATCGCTAGTAATCGTAGATCAGCATGCTACGGTGAATACGTTCCCGGGCCTTGTACACACCGCCCGTCACACCACGGGAGTTGGTTGTACCAGAAGCAGTTGAGCGAACTATTTATAGACGCAGGCTGCCAAGGTATGATTGGTAACTGGGGTG

>GuaymasBasin_Dive4872_L-ASV_560

AACGAACGCTGGCGGCGTGCTTAACACATGCAAGTCGAACGCGAAAGTTTTCTTCGGAAGATGAGTAGAGTGGCGCACGGGTGAGTAACGCGTAAATAATCTACCCCTGCATCTGGGATAACCCACCGAAAGGTGTGCTAATACCGGATACGTTCTTTTGATCGCGAGATCTTAAGAAGAAAGGTGGCCTCTGATATAAGCTACTGTGCGGGGAGGAGTTTGCGTACCATTAGCTAGTTGGTAGGGTAATGGCCTACCAAGGCAACGATGGTTAGCGGGTCTGAGAGGATGATCCGCCACACTGGAACTGGAACACGGACCAGACTCCTACGGGAGGCAGCAGTGAGGAATATTGCGCAATGGGGGCAACCCTGACGCAGCGACGCCGCGTGGATGATGAAGGCCTTCGGGTCGTAAAATCCTGTCAGATGGAAAGAAATGTTATATGGATAATACCTGTATAGCTTGACGGTACCATCAAAGGAAGCACCGGCTAACTCCGTGCCAGCAGCCGCGGTAATACGGAGGGTGCAAGCGTTGTTCGGAATTACTGGGCGTAAAGCGCGCGTAGGTGGTCTGTTATGTCAGATGTGAAAGTCCACGGCTCAACCGTGGAAGTGCATTTGAAACTGGCAGACTTGAGTACTGGAGGGGGTAGTGGAATTCCCGGTGTAGAGGTGAAATTCGTAGATATCGGGAGGAATACCGGTGGCGAAGGCGACTACCTGGCCAGATACTGACACTGAGGTGCGAAAGCGTGGGGAGCGAACAGGATTAGATACCCTGGTAGTCCACGCCGTAAACGATGTCAACTAGGTGTTGGGATGGTTAATCGTCTCATTGCCGCAGCTAACGCATTAAGTTGACCGCCTGGGGAGTACGGTCGCAAGATTAAAACTCAAAGGAATTGACGGGGGCCCGCACAAGCGGTGGAGTATGTGGTTTAATTCGACGCAACGCGCAGAACCTTACCTGGTCTTGACATCCCGGGAAGCTTCAGGAAACTGGAGTGTGCCTCTTGAGGAACCCGGTGACAGGTGCTGCATGGCTGTCGTCAGCTCGTGTCGTGAGATGTTGGGTTAAGTCCCGCAACGAGCGCAACCCTTGTCTTTAGTTGCCATCATTAAGTTGGGCACTCTAAAGAGACTGCCGGTGTCAAACCGGAGGAAGGTGGGGATGACGTCAAGTCCTCATGGCCTTTATGACCAGGGCTACACACGTACTACAATGGCATAGACAAAGGGCAGCGACATCGCGAGGTGAAGCGAATCCCATAAACTATGTCTCAGTCCGGATTGGAGTCTGCAACTCGACTCCATGAAGTTGGAATCGCTAGTAATCGTAGATCAGCATGCTACGGTGAATACGTTCCCGGGCCTTGTACACACCGCCCGTCACACCACGGGAGTTGGTTGTACCAGAAGCAGTTGAGCGAACTATTTATAGACGCAGGCTGCCAAGGTATGATTGGTAACTGGGGTG

>GuaymasBasin_Dive4872_L-ASV_924

AACGAACGCTGGCGGCGTGCTTAACACATGCAAGTCGAACGCGAAAGTTTTCTTCGGAAGATGAGTAGAGTGGCGCACGGGTGAGTAACGCGTAAATAATCTACCCCTGCATTCGGGATAACCCACCGAAAGGTGTGCTAATACCGGATACGTTCTTTTGATCGCGAGATTAAGAGAAGAAAGGTGGCCTCTGATATAAGCTACTGTGCGGGGAGGAGTTTGCGTACCATTAGCTAGTTGGTAGGGTAATGGCCTACCAAGGCAACGATGGTTAGCGGGTCTGAGAGGATGATCCGCCACACTGGAACTGGAACACGGACCAGACTCCTACGGGAGGCAGCAGTGAGGAATATTGCGCAATGGGGGAAACCCTGACGCAGCGACGCCGCGTGGATGATGAAGGCCTTCGGGTCGTAAAATCCTGTCAGATGGAAAGAAATGTATATGTATTAATACTATGTATGCTTGACGGTACCATCAAAGGAAGCACCGGCTAACTCCGTGCCAGCAGCCGCGGTAATACGGAGGGTGCAAGCGTTGTTCGGAATTACTGGGCGTAAAGCGCGCGTAGGTGGTCTGTTATGTCAGATGTGAAAGTCCACGGCTCAACCGTGGAAGTGCATTTGAAACTGGCAGACTTGAGTACTGGAGGGGGTAGTGGAATTCCCGGTGTAGAGGTGAAATTCGTAGATATCGGGAGGAATACCGGTGGCGAAGGCGACTACCTGGCCAGATACTGACACTGAGGTGCGAAAGCGTGGGGAGCGAACAGGATTAGATACCCTGGTAGTCCACGCCGTAAACGATGTCAACTAGGTGTTGGGATGGTTAATCGTCTCATTGCCGCAGCTAACGCATTAAGTTGACCGCCTGGGGAGTACGGTCGCAAGATTAAAACTCAAAGGAATTGACGGGGGCCCGCACAAGCGGTGGAGTATGTGGTTTAATTCGACGCAACGCGCAGAACCTTACCTGGTCTTGACATCCCGAGAAGCTTTAGGAAACTAGAGTGTGCCTCTTGAGGAACTCGGTGACAGGTGCTGCATGGCTGTCGTCAGCTCGTGTCGTGAGATGTTGGGTTAAGTCCCGCAACGAGCGCAACCCTTGTCTTTAGTTGCCATCATTAAGTTGGGCACTCTAAAGAGACTGCCGGTGTCAAACCGGAGGAAGGTGGGGATGACGTCAAGTCCTCATGGCCTTTATGACCAGGGCTACACACGTACTACAATGGCATAGACAAAGGGCAGCGACATCGCGAGGTGAAGCGAATCCCATAAACTATGTCCCAGTCCGGATTGGAGTCTGCAACTCGACTCCATGAAGTTGGAATCGCTAGTAATCGTAGATCAGCATGCTACGGTGAATACGTTCCCGGGCCTTGTACACACCGCCCGTCACACCACGGGAGTTGGTTGTACCAGAAGCAGTTGAGCGAACTATTCGTAGACGCAGGCTGCCAAGGTATGATTGGTAACTGGGGTG

>GuaymasBasin_Dive4872_L-ASV_441

AACGAACGCTGGCGGCGTGCTTAACACATGCAAGTCGAACGCGAAAGTTTTCTTCGGAAGATGAGTAGAGTGGCGCACGGGTGAGTAACGCGTAAATAATCTACCCCTGCATTCGGGATAACCCACCGAAAGGTGTGCTAATACCGGATACGTTCTTTTGATCGCGAGATTAAGAGAAGAAAGGTGGCCTCTGATATAAGCTACTGTGCGGGGAGGAGTTTGCGTACCATTAGCTAGTTGGTAGGGTAATGGCCTACCAAGGCAACGATGGTTAGCGGGTCTGAGAGGATGATCCGCCACACTGGAACTGGAACACGGACCAGACTCCTACGGGAGGCAGCAGTGAGGAATATTGCGCAATGGGGGCAACCCTGACGCAGCGACGCCGCGTGGATGATGAAGGCCTTCGGGTCGTAAAATCCTGTCAGATGGAAAGAAATGTATATGTATTAATACTATGTATGCTTGACGGTACCATCAAAGGAAGCACCGGCTAACTCCGTGCCAGCAGCCGCGGTAATACGGAGGGTGCAAGCGTTGTTCGGAATTACTGGGCGTAAAGCGCGCGTAGGTGGTCTGTTATGTCAGATGTGAAAGTCCACGGCTCAACCGTGGAAGTGCATTTGAAACTGGCAGACTTGAGTACTGGAGGGGGTAGTGGAATTCCCGGTGTAGAGGTGAAATTCGTAGATATCGGGAGGAATACCGGTGGCGAAGGCGACTACCTGGCCAGATACTGACACTGAGGTGCGAAAGCGTGGGGAGCGAACAGGATTAGATACCCTGGTAGTCCACGCCGTAAACGATGTCAACTAGGTGTTGGGATGGTTAATCGTCTCATTGCCGCAGCTAACGCATTAAGTTGACCGCCTGGGGAGTACGGTCGCAAGATTAAAACTCAAAGGAATTGACGGGGGCCCGCACAAGCGGTGGAGCATGTGGTTTAATTCGACGCAACGCGCAGAACCTTACCTGGTCTTGACATCCCGAGAAGCTTTAGGAAACTAGAGTGTGCCTCTTGAGGAACTCGGTGACAGGTGCTGCATGGCTGTCGTCAGCTCGTGTCGTGAGATGTTGGGTTAAGTCCCGCAACGAGCGCAACCCTTGTCTTTAGTTGCCATCATTAAGTTGGGCACTCTAAAGAGACTGCCGGTGTCAAACCGGAGGAAGGTGGGGATGACGTCAAGTCCTCATGGCCTTTATGACCAGGGCTACACACGTACTACAATGGCATAGACAAAGGGCAGCGACATCGCGAGGTGAAGCGAATCCCATAAACTATGTCCCAGTCCGGATTGGAGTCTGCAACTCGACTCCATGAAGTTGGAATCGCTAGTAATCGTAGATCAGCATGCTACGGTGAATACGTTCCCGGGCCTTGTACACACCGCCCGTCACACCACGGGAGTTGGTTGTACCAGAAGCAGTTGAGCGAACTATTCGTAGACGCAGGCTGCCAAGGTATGATTGGTAACTGGGGTG

>GuaymasBasin_Dive4872_L-ASV_85

AACGAACGCTGGCGGCGTGCTTAACACATGCAAGTCGAACGCGAAAGTTTTCTTCGGAAGATGAGTAGAGTGGCGCACGGGTGAGTAACGCGTAAATAATCTACCCCTGCATTCGGGATAACCCACCGAAAGGTGTGCTAATACCGGATACGTTCTTTTGATCGCGAGATTAAGAGAAGAAAGGTGGCCTCTGATATAAGCTACTGTGCGGGGAGGAGTTTGCGTACCATTAGCTAGTTGGTAGGGTAATGGCCTACCAAGGCAACGATGGTTAGCGGGTCTGAGAGGATGATCCGCCACACTGGAACTGGAACACGGACCAGACTCCTACGGGAGGCAGCAGTGAGGAATATTGCGCAATGGGGGCAACCCTGACGCAGCGACGCCGCGTGGATGATGAAGGCCTTCGGGTCGTAAAATCCTGTCAGATGGAAAGAAATGTATATGTATTAATACTATGTATGCTTGACGGTACCATCAAAGGAAGCACCGGCTAACTCCGTGCCAGCAGCCGCGGTAATACGGAGGGTGCAAGCGTTGTTCGGAATTACTGGGCGTAAAGCGCGCGTAGGTGGTCTGTTATGTCAGATGTGAAAGTCCACGGCTCAACCGTGGAAGTGCATTTGAAACTGGCAGACTTGAGTACTGGAGGGGGTAGTGGAATTCCCGGTGTAGAGGTGAAATTCGTAGATATCGGGAGGAATACCGGTGGCGAAGGCGACTACCTGGCCAGATACTGACACTGAGGTGCGAAAGCGTGGGGAGCGAACAGGATTAGATACCCTGGTAGTCCACGCCGTAAACGATGTCAACTAGGTGTTGGGATGGTTAATCGTCTCATTGCCGCAGCTAACGCATTAAGTTGACCGCCTGGGGAGTACGGTCGCAAGATTAAAACTCAAAGGAATTGACGGGGGCCCGCACAAGCGGTGGAGTATGTGGTTTAATTCGACGCAACGCGCAGAACCTTACCTGGTCTTGACATCCCGAGAAGCTTTAGGAAACTAGAGTGTGCCTCTTGAGGAACTCGGTGACAGGTGCTGCATGGCTGTCGTCAGCTCGTGTCGTGAGATGTTGGGTTAAGTCCCGCAACGAGCGCAACCCTTGTCTTTAGTTGCCATCATTAAGTTGGGCACTCTAAAGAGACTGCCGGTGTCAAACCGGAGGAAGGTGGGGATGACGTCAAGTCCTCATGGCCTTTATGACCAGGGCTACACACGTACTACAATGGCATAGACAAAGGGCAGCGACATCGCGAGGTGAAGCGAATCCCATAAACTATGTCCCAGTCCGGATTGGAGTCTGCAACTCGACTCCATGAAGTTGGAATCGCTAGTAATCGTAGATCAGCATGCTACGGTGAATACGTTCCCGGGCCTTGTACACACCGCCCGTCACACCACGGGAGTTGGTTGTACCAGAAGCAGTTGAGCGAACTATTCGTAGACGCAGGCTGCCAAGGTATGATTGGTAACTGGGGTG

>GuaymasBasin_Dive4872_L-ASV_680

AACGAACGCTGGCGGCGTGCTTAACACATGCAAGTCGAACGCGAAAGTTTTCTTCGGAAGACGAGTAGAGTGGCGCACGGGTGAGTAACGCGTAAATAATCTACCCCTGCATTCGGGATAACCCACCGAAAGGTGTGCTAATACCGGATACGTTCTTTTGATCGCGAGATTAAGAGAAGAAAGGTGGCCTCTGATATAAGCTACTGTGCGGGGAGGAGTTTGCGTACCATTAGCTAGTTGGTAGGGTAATGGCCTACCAAGGCAACGATGGTTAGCGGGTCTGAGAGGATGATCCGCCACACTGGAACTGGAACACGGACCAGACTCCTACGGGAGGCAGCAGTGAGGAATATTGCGCAATGGGGGCAACCCTGACGCAGCGACGCCGCGTGGATGATGAAGGCCTTCGGGTCGTAAAATCCTGTCAGATGGAAAGAAATGTATATGTATTAATACTATGTATGCTTGACGGTACCATCAAAGGAAGCACCGGCTAACTCCGTGCCAGCAGCCGCGGTAATACGGAGGGTGCAAGCGTTGTTCGGAATTACTGGGCGTAAAGCGCGCGTAGGTGGTCTGTTATGTCAGATGTGAAAGTCCACGGCTCAACCGTGGAAGTGCATTTGAAACTGGCAGACTTGAGTACTGGAGGGGGTAGTGGAATTCCCGGTGTAGAGGTGAAATTCGTAGATATCGGGAGGAATACCGGTGGCGAAGGCGACTACCTGGCCAGATACTGACACTGAGGTGCGAAAGCGTGGGGAGCGAACAGGATTAGATACCCTGGTAGTCCACGCCGTAAACGATGTCAACTAGGTGTTGGGATGGTTAATCGTCTCATTGCCGCAGCTAACGCATTAAGTTGACCGCCTGGGGAGTACGGTCGCAAGATTAAAACTCAAAGGAATTGACGGGGGCCCGCACAAGCGGTGGAGTATGTGGTTTAATTCGACGCAACGCGCAGAACCTTACCTGGTCTTGACATCCCGAGAAGCTTTAGGAAACTAGAGTGTGCCTCTTGAGGAACTCGGTGACAGGTGCTGCATGGCTGTCGTCAGCTCGTGTCGTGAGATGTTGGGTTAAGTCCCGCAACGAGCGCAACCCTTGTCTTTAGTTGCCATCATTAAGTTGGGCACTCTAAAGAGACTGCCGGTGTCAAACCGGAGGAAGGTGGGGATGACGTCAAGTCCTCATGGCCTTTATGACCAGGGCTACACACGTACTACAATGGCATAGACAAAGGGCAGCGACATCGCGAGGTGAAGCGAATCCCATAAACTATGTCCCAGTCCGGATTGGAGTCTGCAACTCGACTCCATGAAGTTGGAATCGCTAGTAATCGTAGATCAGCATGCTACGGTGAATACGTTCCCGGGCCTTGTACACACCGCCCGTCACACCACGGGAGTTGGTTGTACCAGAAGCAGTTGAGCGAACTATTCGTAGACGCAGGCTGCCAAGGTATGATTGGTAACTGGGGTG

>GuaymasBasin_Dive4872_L-ASV_945

AACGAACGCTGGCGGCGTGCTTAACACATGCAAGTCGAACGCGAAAGTTTTCTTCGGAAAATGAGTAGAGTGGCGCACGGGTGAGTAACGCGTAAATAATCTACCCCTGCATCCGGGATAACCCACCGAAAGGTGTGCTAATACCGGATACGTTCTTTTGATCGCGAGATCTTAAGAAGAAAGGTGGCCTCTGATATAAGCTACTGTGCGGGGAGGAGTTTGCGTACCATTAGCTAGTTGGCAGGGTAATGGCCTACCAAGGCAACGATGGTTAGCGGGTCTGAGAGGATGATCCGCCACACTGGAACTGGAACACGGACCAGACTCCTACGGGAGGCAGCAGTGAGGAATATTGCGCAATGGGGGCAACCCTGACGCAGCGACGCCGCGTGGATGATGAAGGCCTTCGGGTCGTAAAATCCTGTCAGATGGAAAGAAGTGCATATGGATTAATACTTCATATGTTTGACGGTACCATCAAAGGAAGCACCGGCTAACTCCGTGCCAGCAGCCGCGGTAATACGGAGGGTGCAAGCGTTGTTCGGAATTACTGGGCGTAAAGCGCGCGTAGGTGGTCTGTTATGTCAGATGTGAAAGTCCACGGCTCAACCGTGGAAGTGCATTTGAAACTGGCAGACTTGAGTACTGGAGGGGGTAGTGGAATTCCCGGTGTAGAGGTGAAATTCGTAGATATCGGGAGGAATACCGGTGGCGAAGGCGACTACCTGGCCAGATACTGACACTGAGGTGCGAAAGCGTGGGGAGCAAACAGGATTAGATACCCTGGTAGTCCACGCCGTAAACGATGTCAACTAGGTGTTGGGATGGTTAATCGTCTCATTGCCGGAGCTAACGCATTAAGTTGACCGCCTGGGGAGTACGGTCGCAAGATTAAAACTCAAAGGAATTGACGGGGGCCCGCACAAGCGGTGGAGTATGTGGTTTAATTCGACGCAACGCGCAGAACCTTACCTGGTCTTGACATCCCGAGAATCTTCTGGAAACAGAAGAGTGCCTCTTGAGGAACTCGGTGACAGGTGCTGCATGGCTGTCGTCAGCTCGTGTCGTGAGATGTTGGGTTAAGTCCCGCAACGAGCGCAACCCTTGTCTTTAGTTGCCATCATTTAGTTGGGCACTCTAAAGAGACTGCCGGTGTCAAACCGGAGGAAGGTGGGGATGACGTCAAGTCCTCATGGCCTTTATGACCAGGGCTACACACGTACTACAATGGCATAGACAAAGGGCAGCGACATCGCGAGGTGAAGCGAATCCCATAAACTATGTCTCAGTCCGGATTGGAGTCTGCAACTCGACTCCATGAAGTTGGAATCGCTAGTAATCGTAGATCAGCATGCTACGGTGAATACGTTCCCGGGCCTTGTACACACCGCCCGTCACACCACGGGAGTTGGTTGTACCAGAAGCAGTTGAGCGAACCCCGCTACTAAATTATGAGAGGAGATGTTTTTAGCAGGAGCATTTGCTTTATAATTTAGTAGCGGGGGCGCAGGCTGCCAAGGTATGATTGGTAACTGGGGTG

>GuaymasBasin_Dive4872_L-ASV_1970

AACGAACGCTGGCGGCGTGCTTAACACATGCAAGTCGAACGCGAAAGTTTTCTTCGGAAGATGAGTAGAGTGGCGCACGGGTGAGTAACGCGTAAATAATCTACCCTTGCATTTGGGATAACCAACCGAAAGGTTGGCTAATACCGGATACGTTCTTTTGATCGCGAGATCTTAAGAAGAAAGGTGGCCTCTGATATAAGCTACTGTGCAGGGAGGAGTTTGCGTACCATTAGCTAGTTGGTAGGGTAATGGCCTACCAAGGCAACGATGGTTAGCGGGTCTGAGAGGATGATCCGCCACACTGGAACTGGAACACGGACCAGACTCCTACTGGAGGCAGCAGTGAGGAATATTGCGCAATGGGGGCAACCCTGACGCAGCGACGCCGCGTGGATGATGAAGGCCTTCGGGTCGTAAAATCCTGTCAGATGGAAAGAAATGTTATATGGCTAATATCTGTATAGCTTGACGGTACCATCAAAGGAAGCACCGGCTAACTCCGTGCCAGCAGCCGCGGTAATACGGAGGGTGCAAGCGTTGTTCGGAATTACTGGGCGTAAAGCGCGCGTAGGTGGTTTGTTATGTCAGATGTGAAAGTCCACGGCTCAACCGTGGAAGTGCATTTGAAACTGGCAGACTTGAGTACTGGAGGGGGTAGTGGAATTCCCGGTGTAGAGGTGAAATTCGTAGATATCGGGAGGAATACCGGTGGCGAAGGCGACTACCTGGCCAGATACTGACACTGAGGTGCGAAAGCGTGGGGAGCAAACAGGATTAGATACCCTGGTAGTCCACGCCGTAAACGATGTCAACTAGGTGTTGGGATGGTTAATCGTCTCATTGCCGGAGCTAACGCATTAAGTTGACCGCCTGGGGAGTACGGTCGCAAGATTAAAACTCAAAGGAATTGACGGGGGCCCGCACAAGCGGTGGAGCATGTGGTTTAATTCGACGCAACGCGCAGAACCTTACCTGGTCTTGACATCCCGAGAATCCCTGAGAAATCAGGGAGTGCCTCTTGAGGAACTCGGTGACAGGTGCTGCATGGCTGTCGTCAGCTCGTGTCGTGAGATGTTGGGTTAAGTCCCGCAACGAGCGCAACCCTTGTCTTTAGTTGCCATCATTAAGTTGGGCACTCTAAAGAGACTGCCGGTGTCAAACCGGAGGAAGGTGGGGATGACGTCAAGTCCTCATGGCCTTTATGACCAGGGCTACACACGTACTACAATGGCATAGACAAAGGGCAGCGACATCGCGAGGTGAAGCGAATCCCATAAACTATGTCCCAGTCCGGATTGGAGTCTGCAACTCGACTCCATGAAGTTGGAATCGCTAGTAATCGTAGATCAGCATGCTACGGTGAATACGTTCCCGGGCCTTGTACACACCGCCCGTCACACCACGGGAGTTGGTTGTACCAGAAGCAGTTGAGCGAACTATTCGTAGGCGCAGGCTGCCAAGGTATGATTGGTAACTGGGGTG

>GuaymasBasin_Dive4872_L-ASV_385

AACGAACGCTGGCGGCGTGCTTAACACATGCAAGTCGAACGCGAAAGTTTTCTTCGGAAGATGAGTAGAGTGGCGCACGGGTGAGTAACGCGTAAATAATCTACCCTTGCATCTGGGATAACCCACCGAAAGGTGTGCTAATACCGGATACGTTCTTTTGATCGCGAGATTGGAAGAAGAAAGGTGGCCTCTGATATAAGCTACTGTGCAGGGAGGAGTTTGCGTACCATTAGCTAGTTGGTAGGGTAATGGCCTACCAAGGCAACGATGGTTAGCGGGTCTGAGAGGATGATCCGCCACACTGGAACTGGAACACGGACCAGACTCCTACGGGAGGCAGCAGTGAGGAATATTGCGCAATGGGGGCAACCCTGACGCAGCGACGCCGCGTGGATGATGAAGGCCTTCGGGTCGTAAAATCCTGTCAGATGGAAAGAAGTGTTATATGGTTAATAACTGTATAGCTTGACGGTACCATCAAAGGAAGCACCGGCTAACTCCGTGCCAGCAGCCGCGGTAATACGGAGGGTGCAAGCGTTGTTCGGAATTACTGGGCGTAAAGCGCGCGTAGGTGGTCTGTTATGTCAGATGTGAAAGTCCACGGCTCAACCGTGGAAGTGCATTTGAAACTGGCAGACTTGAGTACTGGAGGGGGTAGTGGAATTCCCGGTGTAGAGGTGAAATTCGTAGATATCGGGAGGAATACCGGTGGCGAAGGCGACTACCTGGCCAGATACTGACACTGAGGTGCGAAAGCGTGGGGAGCAAACAGGATTAGATACCCTGGTAGTCCACGCCGTAAACGATGTCAACTAGGTGTTGGGATGGTTAATCGTCTCATTGCCGGAGCTAACGCATTAAGTTGACCGCCTGGGGAGTACGGTCGCAAGATTAAAACTCAAAGGAATTGACGGGGGCCCGCACAAGCGGTGGAGTATGTGGTTTAATTCGACGCAACGCGCAGAACCTTACCTGGTCTTGACATCCCGAGAATCCCTGAGAAATCAGGGAGTGCCTCTTGAGGAGCTCGGTGACAGGTGCTGCATGGCTGTCGTCAGCTCGTGTCGTGAGATGTTGGGTTAAGTCCCGCAACGAGCGCAACCCTTGTCTTTAGTTGCCATCATTAAGTTGGGCACTCTAAAGAGACTGCCGGTGTCAAACCGGAGGAAGGTGGGGATGACGTCAAGTCCTCATGGCCTTTATGACCAGGGCTACACACGTACTACAATGGCATAGACAAAGGGCAGCGACATCGCGAGGTGAAGCGAATCCCGTAAACTATGTCTCAGTCCGGATTGGAGTCTGCAACTCGACTCCATGAAGCTGGAATCGCTAGTAATCGTGGATCAGCATGCCACGGTGAATACGTTCCCGGGCCTTGTACACACCGCCCGTCACACCACGGGAGTTGGTTGTACCAGAAGCAGTTGAGCGAACTATTCGTAGACGCAGGCTGCCAAGGTATGATTGGTAACTGGGGTG
