## Supplementary Figures for "Complex bacterial diversity of Guaymas Basin hydrothermal sediments revealed by synthetic long-read sequencing (LoopSeq)"

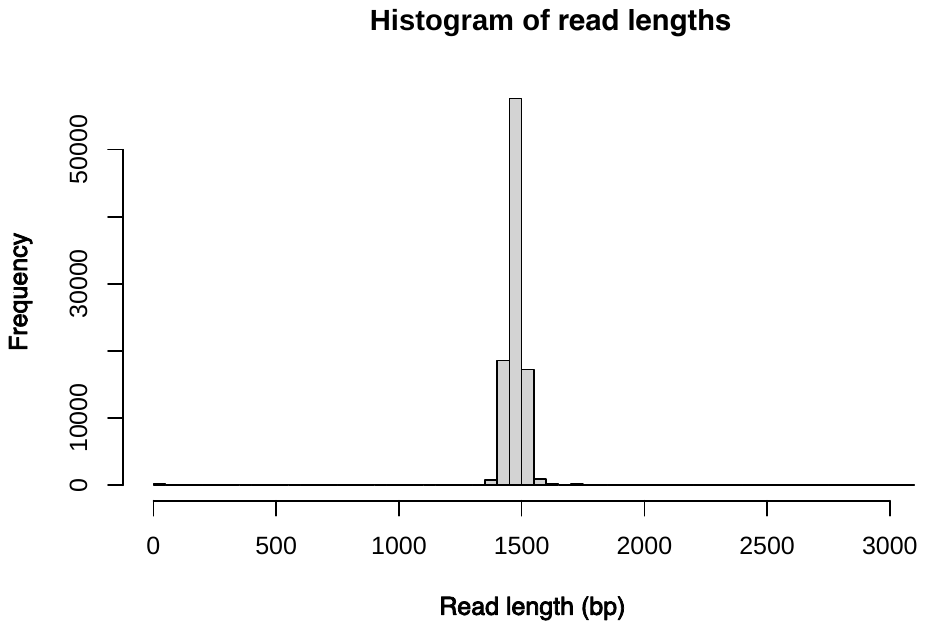

**Supplementary Figure 1.** Histogram of sequence read lengths (in base pairs) as generated by the 27F and 1492R 16S rRNA gene sequence primers utilized by LoopSeq.

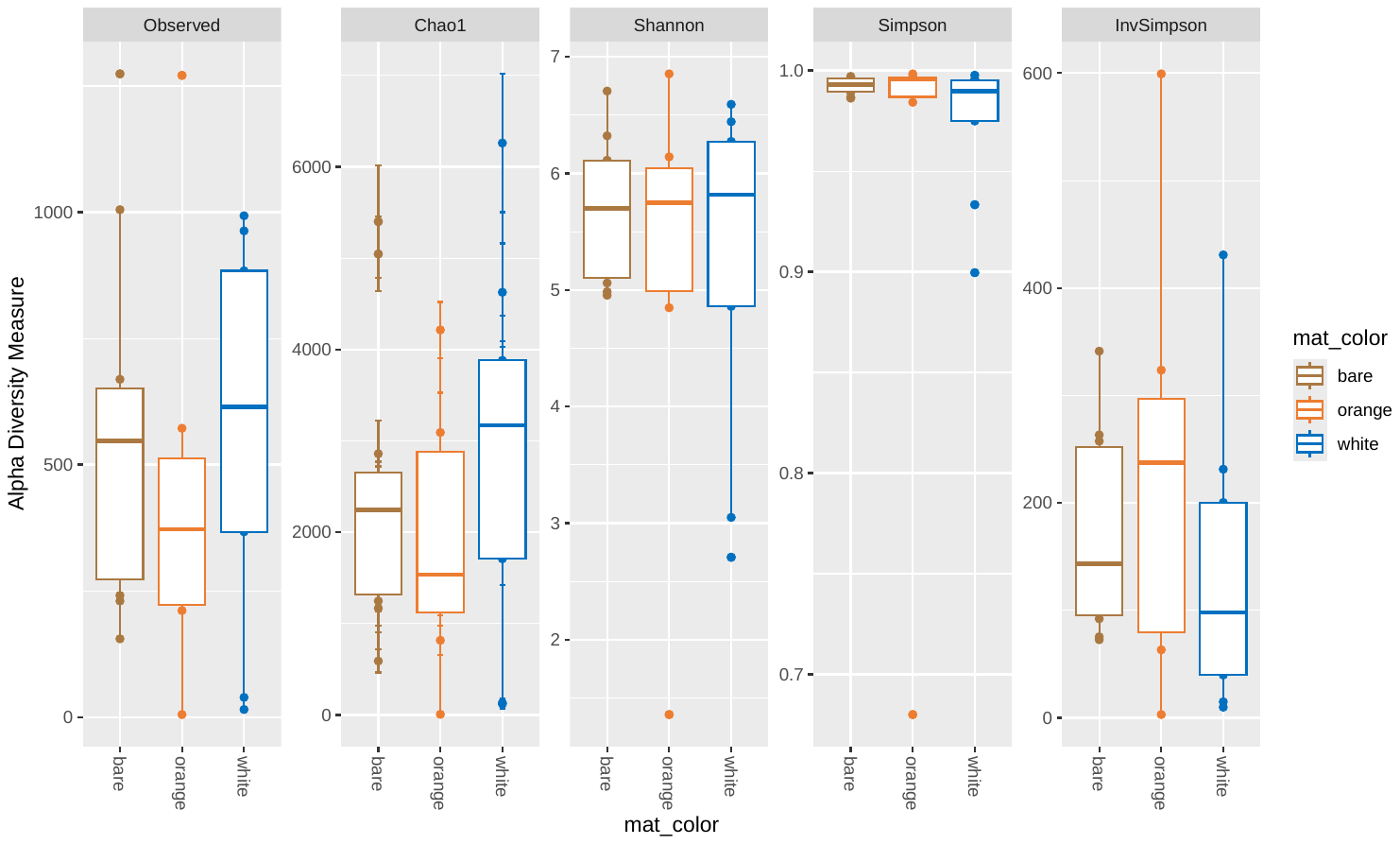

**Supplementary Figure 2.** Alpha diversity figure using the Observed, Chao1, Shannon, Simpson, and Inverse Simpson metrics for the sequencing datasets obtained at the three coring sites (Core 4872-01: the bare sediment site, Core 4872-06: the white mat site, and Core 4872-14: the orange mat site).

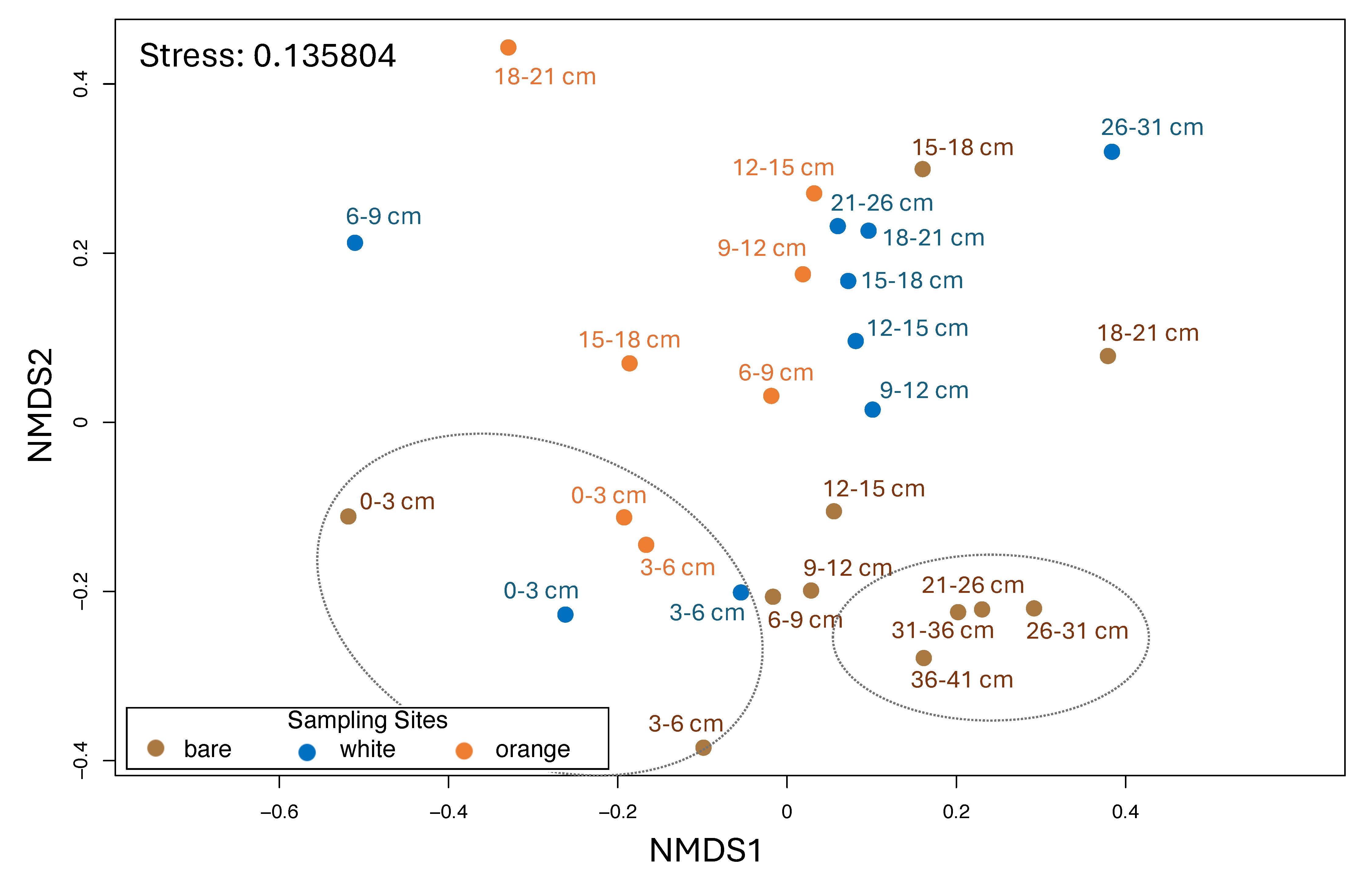

**Supplementary Figure 3.** Non-metric multi-dimensional scaling (NMDS) plot using the Bray-Curtis dissimilarity method. Two clusters are highlighted using stippled lines: I) a cluster of surficial (top 6 cmbsf) samples from all three sites, and II) a cluster of deeper (≥ 21 cmbsf) samples from the bare sediment site.

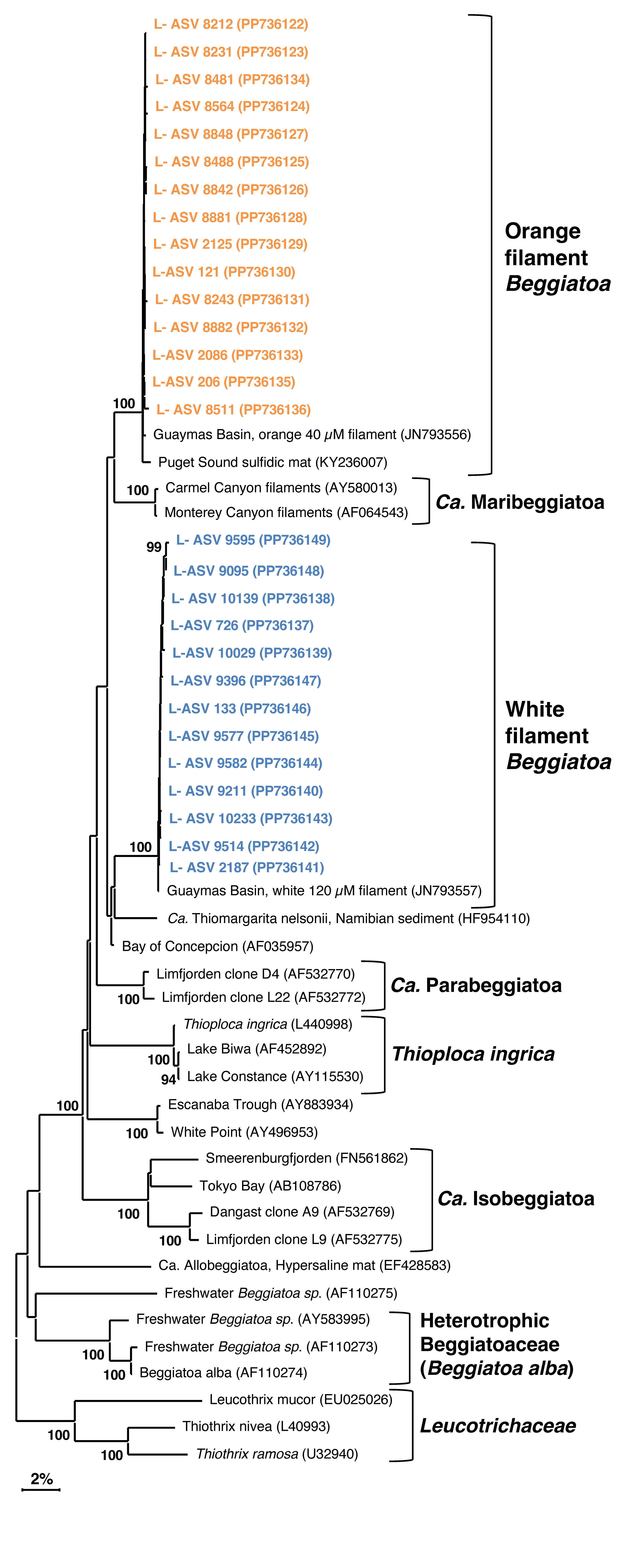

**Supplementary Figure 4.** Expanded distance phylogeny of *Beggiatoaceae* (rooted by *Leucotrichaecea*) depicting all L-ASVs recovered at each mat site (orange, white).

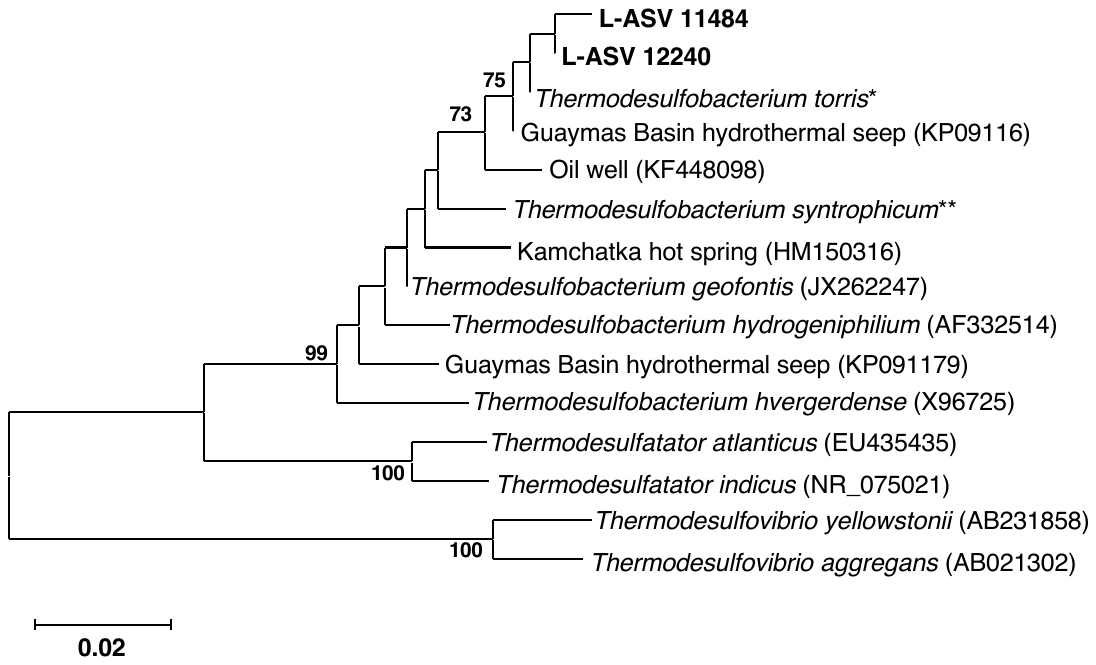

**Supplementary Figure 5.** Distance phylogeny of *Thermodesulfobacterium* and related lineages.

*Thermodesulfobacterium torris 16S rRNA sequence was obtained from annotated genome (BioSample ID: SAMN27514933).

**Thermodesulfobacterium syntrophicum 16S rRNA sequence was obtained from annotated genome (BioSample ID: SAMN29995626).
